# Design and development of lysyl tRNA synthetase inhibitors, for the treatment of tuberculosis

**DOI:** 10.1101/2025.05.13.646596

**Authors:** Susan H. Davis, Michael Mathieson, Kirsteen I. Buchanan, Alice Dawson, Alasdair Smith, Mattia Cocco, Fabio K. Tamaki, John Post, Beatriz Baragaña, Chimed Jansen, Michael Kiczun, Fabio Zuccotto, Gavin Wood, Paul Scullion, Peter C. Ray, Ola Epemolu, Eva Maria Lopez-Román, Laura Guijarro López, Curtis A. Engelhart, Jia Kim, Paula A. Pino, Dirk Schnappinger, Kevin D. Read, Lourdes Encinas, Robert H Bates, Paul G Wyatt, Simon R. Green, Laura A.T. Cleghorn

## Abstract

There is currently a public health crisis due to the rise of multi-drug-resistant tuberculosis cases, as well as the rise in number of deaths from tuberculosis. To achieve the United Nations Sustainable Development Goal of ending the tuberculosis epidemic by 2030, new treatments are urgently required. We previously reported the discovery of **49**, a pre-clinical candidate that acted through inhibition of the *Mycobacterium tuberculosis* lysyl-tRNA synthetase (LysRS). In this report, the full medicinal chemistry program is reviewed from the original hit through to the optimised lead. The work was guided by the first crystal structure of *M. tuberculosis* LysRS. The physicochemical and pharmacokinetic properties were optimised to afford compounds suitable for evaluation in mouse efficacy models of tuberculosis and with the potential for clinical development.

## INTRODUCTION

Despite *Mycobacterium tuberculosis* being known as the causative agent of tuberculosis (TB) since 1882, when it was first identified^1^, treatment of TB is still challenging and lengthy. The number of annual deaths from TB had been in decline between 2005 and 2019, but since the COVID-19 pandemic, the TB incidence rate (new cases per 100 000 population per year) rose by 3.6% between 2020 and 2021; reversing declines of about 2% per year, for most of the previous two decades^2^. The post-pandemic rise in cases has now slowed and started to stabilize. In 2023 the estimated total was 10.8 million, a small increase from 10.7 million in 2022 although still much higher than 10.1 million in 2020^3^. Treatment for drug-susceptible TB is a combination of four frontline drugs: isoniazid, rifampicin, ethambutol and pyrazinamide for a total of 6 months. In 2021 there was a rise in the number of multidrug resistant TB cases that required further treatment with second-line drugs^2^. In 2022, the WHO recommended the 6-month regimen of Bedaquiline, Pretomanid, Linezolid and Moxifloxacin, in cases where there is no known fluoroquinolone resistance; a considerable improvement on traditional 18+ month second line treatments^4^. However, resistance to some of these new drugs is already emerging^5–6^. As such, there is still an urgent demand for alternative TB treatments.

One approach to combating rising clinical resistance, is to identify compounds with novel modes of action, thereby circumventing pre-existing resistance. Aminoacyl-tRNA synthetases are an essential part of the protein biosynthesis machinery, responsible for charging tRNA with the appropriate cognate amino acid for incorporation into growing polypeptide chains. Compounds that target leucyl tRNA synthetase have already entered clinical development for TB treatment^7^. This report focusses on lysyl tRNA synthetase (LysRS), the enzyme that charges lysyl-tRNA with lysine. Molecules have been reported previously targeting LysRS in *Plasmodium* that block parasite growth^8–9^. *LysS* the gene that encodes LysRS is known to be essential for *M. tuberculosis*^10–12^ and has been classified in the top 200 most vulnerable genes to target for inhibition of *M. tuberculosis* growth^13^. We recently published a manuscript on the validation of LysRS as a target for *M. tuberculosis* drug development, including the identification of a compound suitable for preclinical candidate development **49**^14^. Herein, we describe the detailed medicinal chemistry program that underpinned the optimisation of a singleton hit into a preclinical candidate.

## RESULTS AND DISCUSSION

### Hit identification and initial scoping

The starting point, **1** was identified simultaneously as a modest inhibitor of *M. tuberculosis* growth (MIC 20 µM) and a weak inhibitor of *Plasmodium falciparum* LysRS (IC_50_ 5 µM). To exploit this finding, a medicinal chemistry programme was initiated to develop a novel series of *M. tuberculosis* LysRS inhibitors. The hit had favourable druglike properties and an excellent physicochemical profile including impressive solubility (>250 µM). When tested in a *M. tuberculosis* LysRS assay, **1** had modest activity (IC_50_ = 42 µM) which correlated well with its MIC activity (**Table 1**). Compound **1** also had activity against *M. tuberculosis* growing inside macrophages; in fact, the intra and extracellular MIC data tracked very well for the whole synthetic program (see data and PAINS summary in the supplemental information). The crystal structure of *M. tuberculosis* LysRS complexed with lysine, was obtained in-house (PDB:7qh8)^14^ which allowed the synthetic program to be explored utilising structure based design. A key area of focus was to improve both potency against LysRS and selectivity over the human ortholog, KARS1. In the early stages of the project, a modelling comparison between LysRS and KARS1, highlighted a significant point of change within the R_1_ pocket. Specifically, the larger and more hydrohphobic Met271 in LysRS (Thr337 in KARS1) was predicted to “seal-off” the bottom of the R_1_ pocket in LysRS, radically condensing the space available to accommodate ligands. It was hypothesised that this key difference could provide an opportunity to introduce selectivity over KARS1.

**Table 1:**
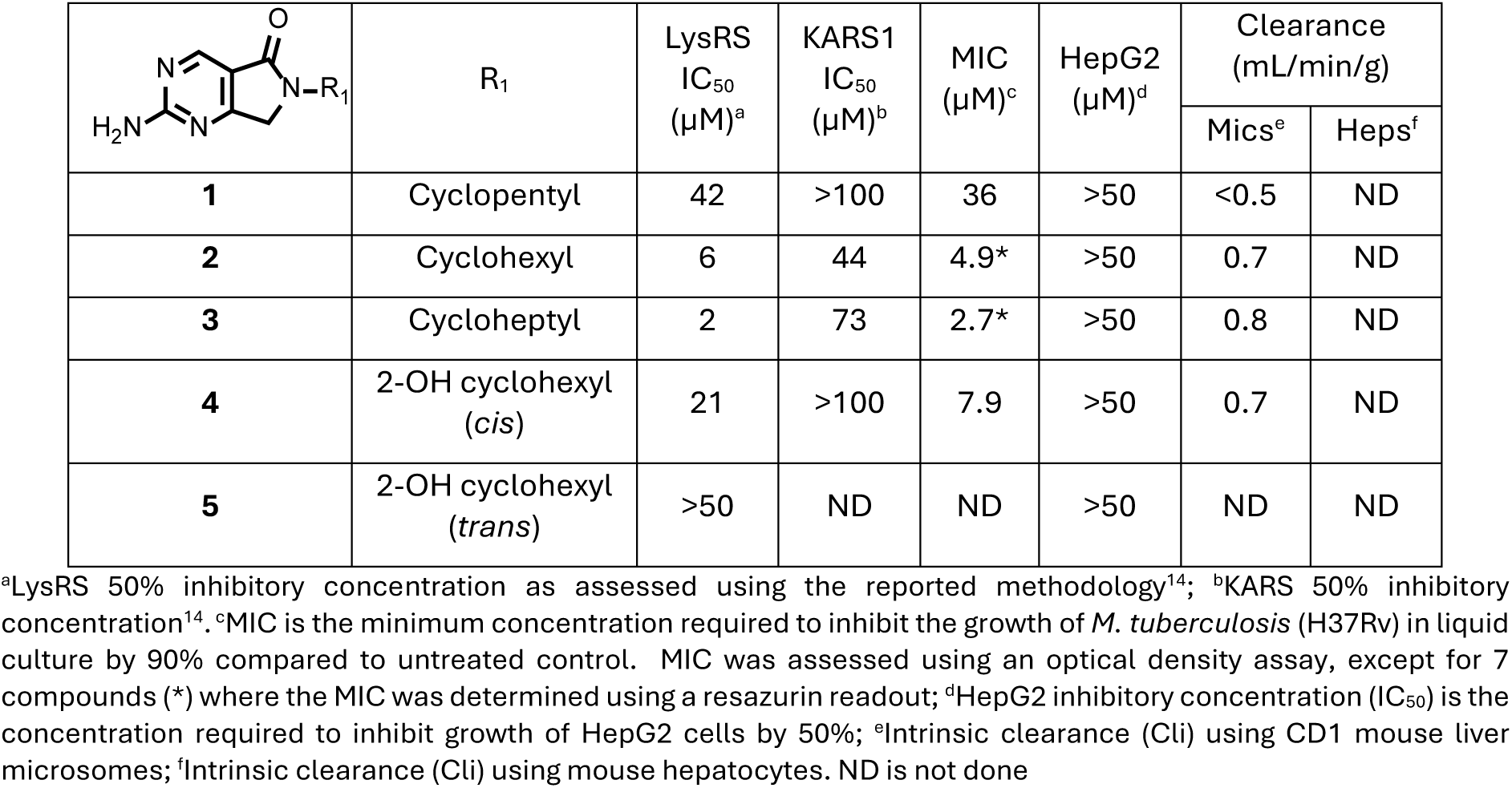
Early SAR modifications.

|  | R<sub>1</sub> | LysRS<br>IC<sub>50</sub><br>(μM)<sup>a</sup> | KARS1<br>IC<sub>50</sub><br>(μM)<sup>b</sup> | MIC<br>(μM)<sup>c</sup> | HepG2<br>(μM)<sup>d</sup> | Clearance<br>(mL/min/g) |  |
| --- | --- | --- | --- | --- | --- | --- | --- |
|  |  |  |  |  |  | Mics<sup>e</sup> | Heps<sup>f</sup> |
| **1** | Cyclopentyl | 42 | >100 | 36 | >50 | <0.5 | ND |
| **2** | Cyclohexyl | 6 | 44 | 4.9* | >50 | 0.7 | ND |
| **3** | Cycloheptyl | 2 | 73 | 2.7* | >50 | 0.8 | ND |
| **4** | 2-OH cyclohexyl<br>(*cis*) | 21 | >100 | 7.9 | >50 | 0.7 | ND |
| **5** | 2-OH cyclohexyl<br>(*trans*) | >50 | ND | ND | >50 | ND | ND |

<sup>a</sup>LysRS 50% inhibitory concentration as assessed using the reported methodology<sup>14</sup>; <sup>b</sup>KARS 50% inhibitory concentration<sup>14</sup>. <sup>c</sup>MIC is the minimum concentration required to inhibit the growth of *M. tuberculosis* (H37Rv) in liquid culture by 90% compared to untreated control. MIC was assessed using an optical density assay, except for 7 compounds (\*) where the MIC was determined using a resazurin readout; <sup>d</sup>HepG2 inhibitory concentration (IC<sub>50</sub>) is the concentration required to inhibit growth of HepG2 cells by 50%; <sup>e</sup>Intrinsic clearance (C<sub>li</sub>) using CD1 mouse liver microsomes; <sup>f</sup>Intrinsic clearance (C<sub>li</sub>) using mouse hepatocytes. ND is not done

#### Early structure activity relationship (SAR) modifications

Guided by the preliminary structural information, the SAR investigation initially concentrated on the cyclopentyl group at R_1_, which occupied a sizable hydrophobic pocket containing several water molecules. Potentially, there was an opportunity to displace these water molecules and enhance potency by ideally filling the pocket. Increasing the ring size to cyclohexyl (**2**) and cycloheptyl (**3**) improved both LysRS potency (7- & 20-fold respectively) and selectivity with **3** displaying ∼40-fold selectivity window vs. KARS1 (**Table1**). The existence of polar residues, like Glu422, within the R_1_ pocket presented an opportunity to introduce polar substituents, capable of binding to this residue. Consequently, straightforward SAR changes were explored on the cyclohexyl ring. The *cis* 2-hydroxy (**4**) analogue was tolerated, although it did result in a modest reduction in potency against LysRS. Interestingly the *trans* 2-hydroxy (**5**) was found to be inactive (**Table 1**). Encouragingly for future development, as with the initial hit, there was an excellent correlation between enzyme and MIC potency observed in these compounds.

### Hit Expansion

#### Development of R_2_ pocket SAR

To investigate further opportunities for enhanced potency and selectivity, substitution at R_2_ was explored (**Table 2**). The initial focus of this work simply employed the incorporation of halogens to explore the chemical space. Chloro (**6**) and bromo (**7**) substituents were more potent than their unsubstituted equivalents (**2** & **3**) resulting in sub micromolar activity against both the enzyme and bacterial growth. Unfortunately, they also significantly increased activity against KARS1 and correspondingly HepG2 cytotoxicity.

**Table 2.**
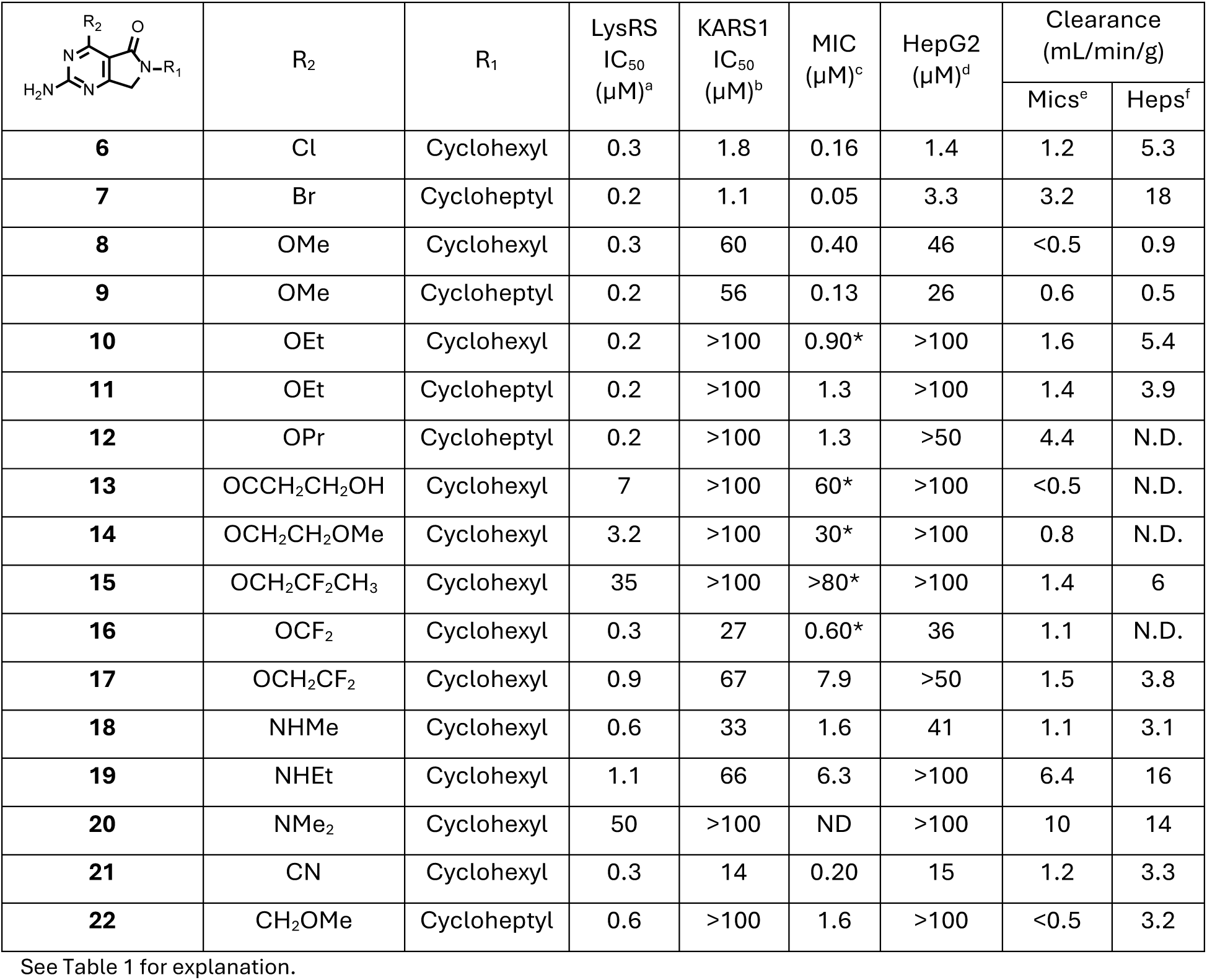
SAR of R_2_ substituent.

|  | R<sub>2</sub> | R<sub>1</sub> | LysRS<br>IC<sub>50</sub><br>(μM)<sup>a</sup> | KARS1<br>IC<sub>50</sub><br>(μM)<sup>b</sup> | MIC<br>(μM)<sup>c</sup> | HepG2<br>(μM)<sup>d</sup> | Clearance<br>(mL/min/g) |  |
| --- | --- | --- | --- | --- | --- | --- | --- | --- |
|  |  |  |  |  |  |  | Mics<sup>e</sup> | Heps<sup>f</sup> |
| **6** | Cl | Cyclohexyl | 0.3 | 1.8 | 0.16 | 1.4 | 1.2 | 5.3 |
| **7** | Br | Cycloheptyl | 0.2 | 1.1 | 0.05 | 3.3 | 3.2 | 18 |
| **8** | OMe | Cyclohexyl | 0.3 | 60 | 0.40 | 46 | <0.5 | 0.9 |
| **9** | OMe | Cycloheptyl | 0.2 | 56 | 0.13 | 26 | 0.6 | 0.5 |
| **10** | OEt | Cyclohexyl | 0.2 | >100 | 0.90* | >100 | 1.6 | 5.4 |
| **11** | OEt | Cycloheptyl | 0.2 | >100 | 1.3 | >100 | 1.4 | 3.9 |
| **12** | OPr | Cycloheptyl | 0.2 | >100 | 1.3 | >50 | 4.4 | N.D. |
| **13** | OCCH<sub>2</sub>CH<sub>2</sub>OH | Cyclohexyl | 7 | >100 | 60* | >100 | <0.5 | N.D. |
| **14** | OCH<sub>2</sub>CH<sub>2</sub>OMe | Cyclohexyl | 3.2 | >100 | 30* | >100 | 0.8 | N.D. |
| **15** | OCH<sub>2</sub>CF<sub>2</sub>CH<sub>3</sub> | Cyclohexyl | 35 | >100 | >80* | >100 | 1.4 | 6 |
| **16** | OCF<sub>2</sub> | Cyclohexyl | 0.3 | 27 | 0.60* | 36 | 1.1 | N.D. |
| **17** | OCH<sub>2</sub>CF<sub>2</sub> | Cyclohexyl | 0.9 | 67 | 7.9 | >50 | 1.5 | 3.8 |
| **18** | NHMe | Cyclohexyl | 0.6 | 33 | 1.6 | 41 | 1.1 | 3.1 |
| **19** | NHEt | Cyclohexyl | 1.1 | 66 | 6.3 | >100 | 6.4 | 16 |
| **20** | NMe<sub>2</sub> | Cyclohexyl | 50 | >100 | ND | >100 | 10 | 14 |
| **21** | CN | Cyclohexyl | 0.3 | 14 | 0.20 | 15 | 1.2 | 3.3 |
| **22** | CH<sub>2</sub>OMe | Cycloheptyl | 0.6 | >100 | 1.6 | >100 | <0.5 | 3.2 |

See Table 1 for explanation.

Methoxy at R_2_ (**8** & **9**) afforded sub micromolar LysRS potency. As with the halogen substitutions, the increase in enzymatic potency was mirrored with increased growth inhibition against the bacteria. In contrast to the halogens the methoxy substituent displayed improved selectivity over KARS1 but retained some HepG2 cytotoxicity. Extending the size of R_2_ to an ethoxy (**10** & **11**) retained good potency against LysRS and significantly, was found to be highly selective over KARS1 and HepG2 (IC_50_ >100 µM). To understand the gain in selectivity with the ethoxy group, a crystal structure of **10** in complex with LysRS was obtained (PDB 9qea). Key interactions formed by the molecule are shown in **Figure 1A**; hydrogen bonds are formed to the main chain carbonyl and amide of Ser266, mimicking the interactions of the adenine group of ATP. The ethoxy oxygen and the carbonyl of the lactam group form a water mediated hydrogen bonded network to Asp480. The crystal structure also indicated that the selectivity observed over KARS1 was primarily driven by the ethoxy group due to increased flexibility in this region of the protein compared to KARS1 (**Figure 1B**). In LysRS residues Asn258 – Pro267 form a loop close to the adenosine binding pocket that adopts differing conformations depending on the presence or absence of a ligand in this pocket (**Figure 1B**). In the closed conformation, the loop is close in space to the C-terminus of the protein, but no interactions are observed. By contrast in KARS1 this loop is always observed in the closed conformation, even in the absence of any ligand in the adenosine pocket. A hydrogen bond is formed between the carbonyl of Leu329 and the amide of Ile564 (**Figure 1B**), reducing the flexibility of this region in KARS1. This limits the ability of the pocket to accommodate larger substituents and explains the selectivity gain caused by the addition of the ethoxy group

**Figure 1:**
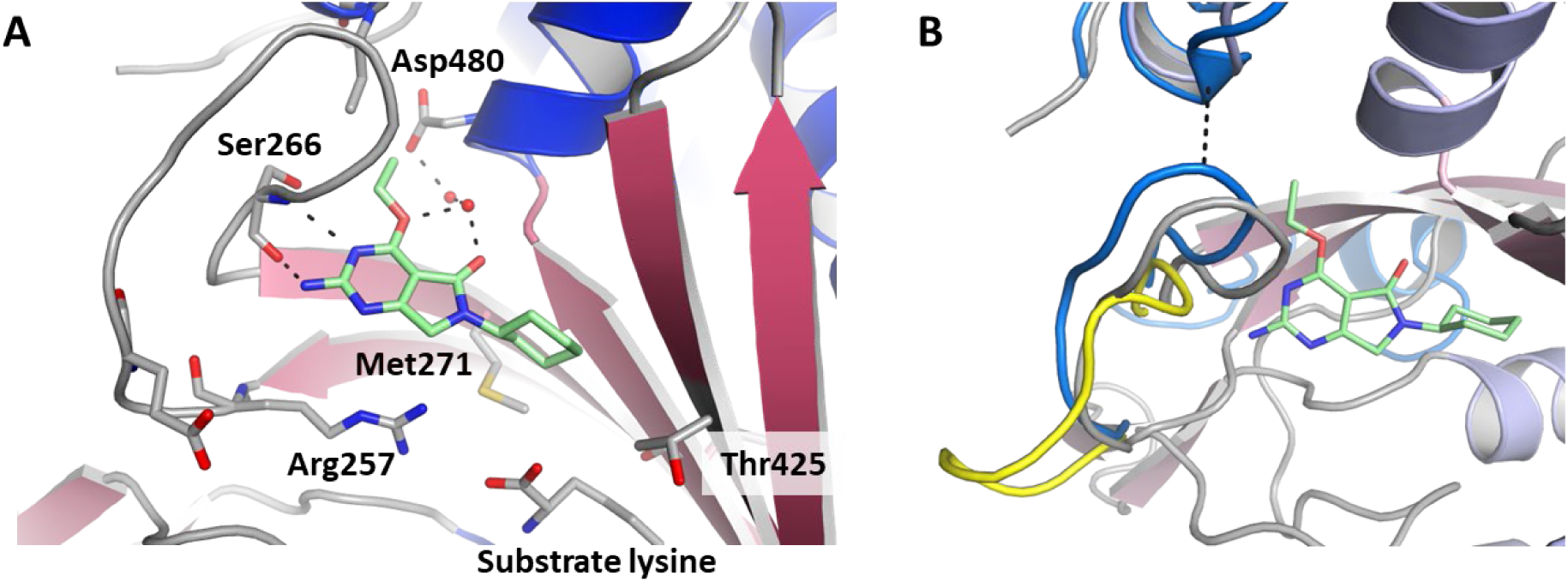
Binding of 10 in the ATP pocket of LysRS (PDB 9qea). **A.** The ligand is shown with mint green carbon atoms, blue nitrogen atoms and red oxygen atoms. Key water molecules are shown as red spheres and hydrogen bonds by dashed lines. Key residues involved with binding are shown. **B.** View of the binding site highlighting the different loop conformations. The ligand bound structure is shown with the same colour scheme as Fig1A, overlayed on this is the loop position of the lysine bound “apo” protein in yellow (PDB 7qh8) and human KARS bound to L-lysylsulfamoyl adenosine in blue (PDB 6chd). Black dashes indicate a hydrogen bond formed in KARS1 that has not been observed in LysRS. Figures prepared using PyMol: The PyMOL Molecular Graphics System, Version 2.5.5 Schrödinger, LLC.

To explore the potential of exploiting the R_2_ pocket to enhance the selectivity window further, a range of alternative groups were examined. Extending the alkyl substituent by one carbon to *n*-propyl, (**12**), continued to improve selectivity (> 400-fold) against KARS1, however the modification led to an increase in microsomal turnover. In a bid to block metabolism on the extended alkyl chain, polarity was introduced at suspected metabolic soft spots by the addition of a hydroxy and methoxy substituent (**13** & **14**). Although this strategy was successful in reducing the *in vitro* microsomal turnover, it resulted in a drop-off in LysRS inhibition. Further design rounds focused on an alternative approach of introducing fluoro substituents to block metabolic soft spots. 2,2-difluoropropoxy (**15**) displayed an improved intrinsic microsomal clearance profile but with reduced LysRS potency. Di-fluoromethoxy at R_2_, (**16**) maintained potency at the primary target LysRS, that was comparable to the methoxy (**8**) but like **8** it still exhibited KARS1 inhibition and HepG2 toxicity. 2,2-difluoroethoxy (**17**), exhibited sub micromolar activity against LysRS but unexpectedly, also exhibited activity against KARS1, unlike the unsubstituted ethoxy analogue (**10**).

After exploring a variety of O-linked substituents at the R_2_ position, the requirement for an oxygen linker was examined by synthesising the relevant N- and C-linked analogues. The N-linked version of **8** (**18**), slightly reduced LysRS potency and afforded a smaller KARS1 selectivity window. Increasing the size of the substituent to ethyl (**19**), did not improve the selectivity as had been observed in the O-linked equivalent analogues. The decrease in activity between the enzymatic and cellular assays was slightly higher than observed with the O-linked, one hypothesis was that the addition of an extra hydrogen bond donor (HBD) had a negative impact on the cellular potency. An added benefit of the N-linked analogues was that they allowed the opportunity to add an additional substitution vector and consequently cap the additional HBD that had been introduced. To explore this possibility, di-methyl amine was introduced at R_2_ and found to be inactive (**20**). In the exploration of C-linked analogues in the R_2_ pocket, a nitrile substituent (**21**) was tolerated, however it did not display a large selectivity window over KARS1 suggesting the shape of the substituent was an important factor to induce the R_2_ loop movement. The C-linked equivalent of **11,** where the O atom was moved along the alkyl chain was prepared (**22)** but was found to result in an unfavourable drop in MIC potency. Thus, the R_2_ SAR exploration demonstrated several significant improvements compared to the original hit, including identifying analogues with sub micromolar activity against both the enzymatic and cellular assays with excellent selectivity over KARS1 and reasonable metabolic stability profile (**Table 2**).

Three compounds (**11**, **10** & **8**) were selected for progression to *in vivo* pharmacokinetic (PK) studies, based on their *in vitro* potency and clearance data (**Table S2**). All had acceptable PK profiles with reasonable exposure and bioavailability. Slight variations in the R_1_ substituent had little impact on the PK profile (where R_2_ was ethoxy and R_1_ = cyclohexyl (**10**) or cycloheptyl (**11**)). A greater difference was observed varying the R_2_ group, with methoxy (**8**) having a significantly lower C_max_ and AUC compared to the ethoxy (**10**) equivalent. Following the PK studies two compounds (**8** & **11**) were selected to be progressed to a proof-of-concept efficacy study in an acute murine model of TB infection^15^ (**Figure 2**). Both compounds, when evaluated at 200 mg/kg, displayed promising results for early molecules in a new series: with statistically significant reductions in the bacterial load being observed (2.6 and 1.9 Log_10_ reductions respectively). This result gave an early validation that inhibiting LysRS would produce an efficacious effect *in vivo* and supported further investigation of the SAR. The *in vitro* and *in vivo* profiling of the hit to lead phase had identified new regions to focus the optimisation upon, including metabolic stability as well as improving the volume of distribution (V_d_) and consequently the *in vivo* half-life.

**Figure 2:**
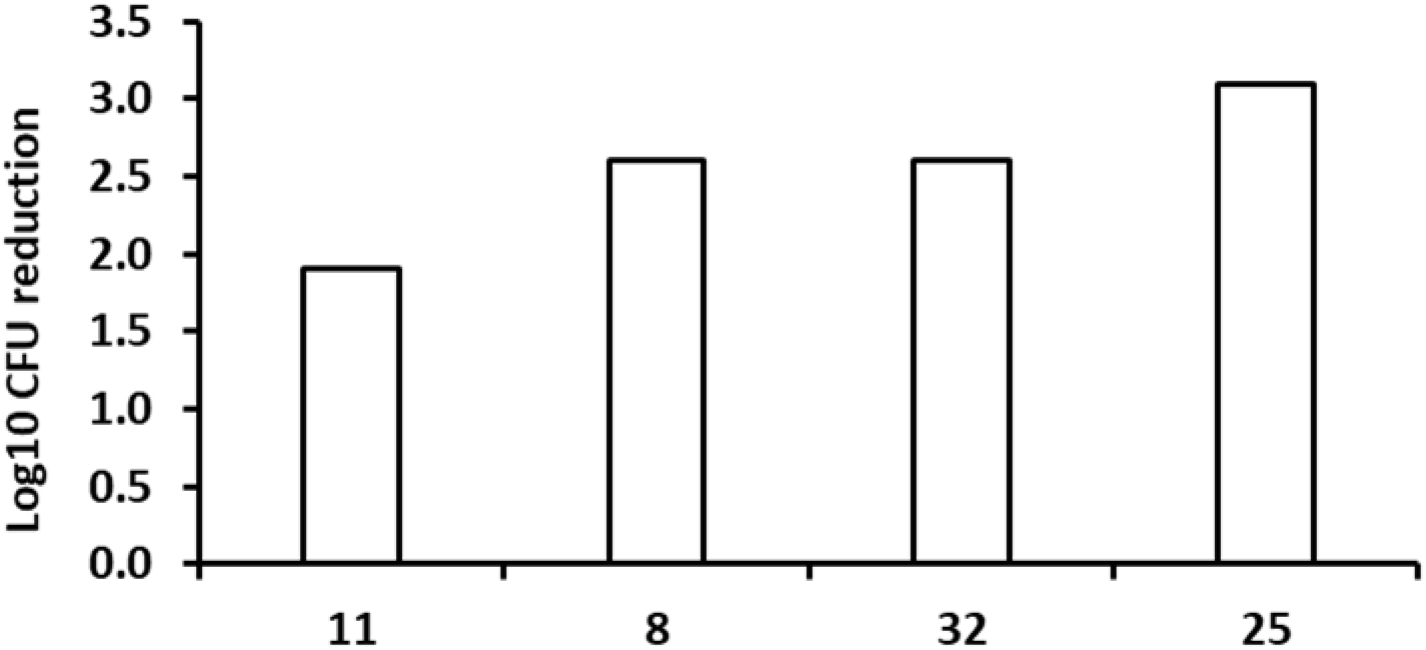
*In vivo* efficacy of early LysRS inhibitors in murine models of infection. Compounds **11**, **8**, **32** and **25** were tested in an acute model of infection in C57BL/6mice. Dosing started 1 day after infection and lasted for 8 days; dosing at 200 mg/kg, each group consisted of two mouse/dose. The effect on the number of colony-forming units (CFU) in mouse lungs is shown.

#### Development of R_1_ pocket SAR

An *in vivo* metabolite ID study was carried out on blood samples taken from the PK study of **11**. This identified oxidation on the R_1_ cycloalkyl ring as the main route of metabolism *in vivo* (**Figure S1**). Utilising the potency gained from addition of a substituent at R_2_, SAR around R_1_ was further explored, with an emphasis on attempting to block the metabolic liability (**Table 3**).

**Table 3:**
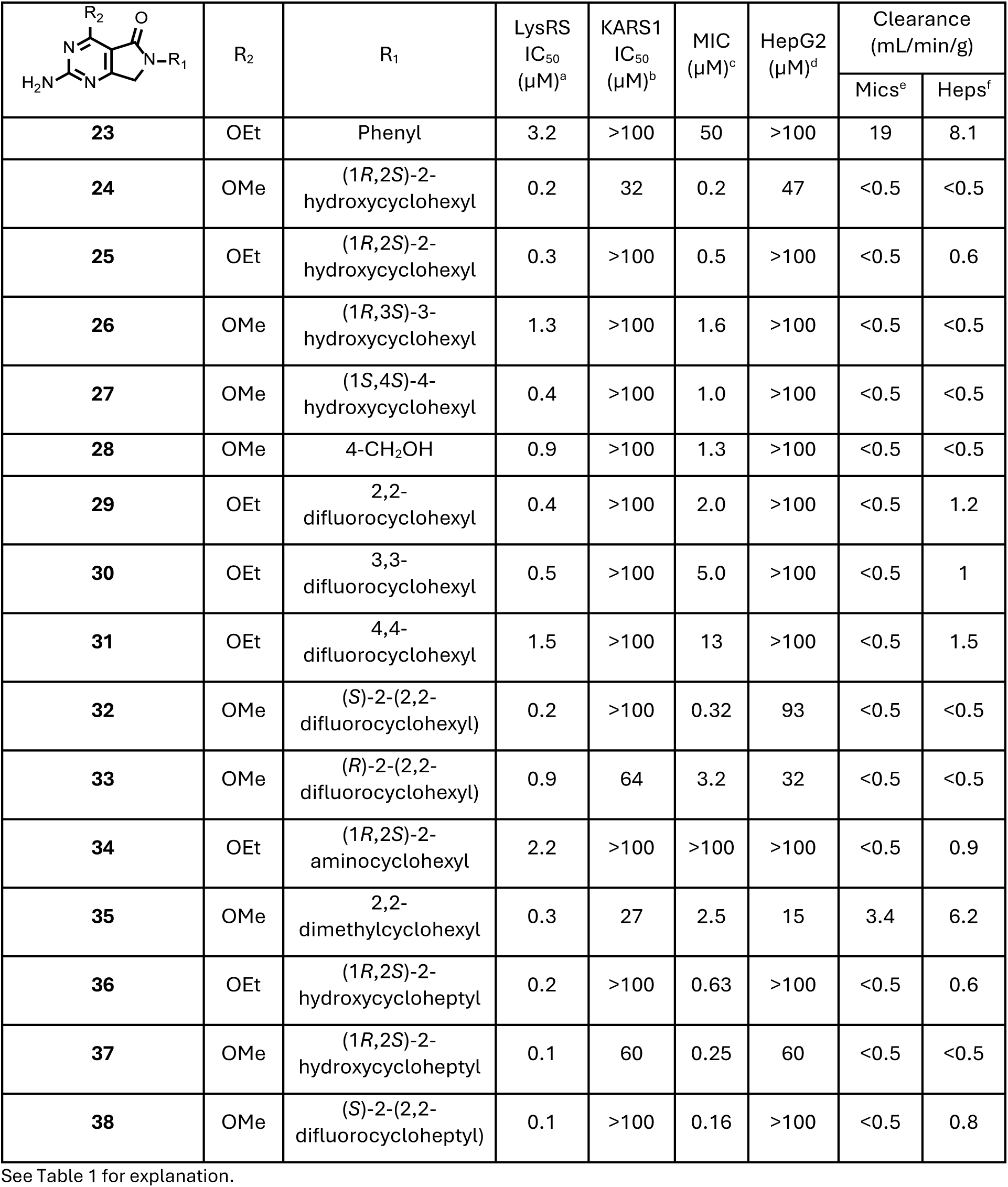
Improving potency and selectivity through the R_1_ pocket.

| | R<sub>2</sub> | R<sub>1</sub> | LysRS<br>IC<sub>50</sub><br>(μM)<sup>a</sup> | KARS1<br>IC<sub>50</sub><br>(μM)<sup>b</sup> | MIC<br>(μM)<sup>c</sup> | HepG2<br>(μM)<sup>d</sup> | Clearance<br>(mL/min/g) | |
| --- | --- | --- | --- | --- | --- | --- | --- | --- |
|  |  |  |  |  |  |  | Mics<sup>e</sup> | Heps<sup>f</sup> |
| **23** | OEt | Phenyl | 3.2 | >100 | 50 | >100 | 19 | 8.1 |
| **24** | OMe | (1*R*,2*S*)-2-hydroxycyclohexyl | 0.2 | 32 | 0.2 | 47 | <0.5 | <0.5 |
| **25** | OEt | (1*R*,2*S*)-2-hydroxycyclohexyl | 0.3 | >100 | 0.5 | >100 | <0.5 | 0.6 |
| **26** | OMe | (1*R*,3*S*)-3-hydroxycyclohexyl | 1.3 | >100 | 1.6 | >100 | <0.5 | <0.5 |
| **27** | OMe | (1*S*,4*S*)-4-hydroxycyclohexyl | 0.4 | >100 | 1.0 | >100 | <0.5 | <0.5 |
| **28** | OMe | 4-CH<sub>2</sub>OH | 0.9 | >100 | 1.3 | >100 | <0.5 | <0.5 |
| **29** | OEt | 2,2-difluorocyclohexyl | 0.4 | >100 | 2.0 | >100 | <0.5 | 1.2 |
| **30** | OEt | 3,3-difluorocyclohexyl | 0.5 | >100 | 5.0 | >100 | <0.5 | 1 |
| **31** | OEt | 4,4-difluorocyclohexyl | 1.5 | >100 | 13 | >100 | <0.5 | 1.5 |
| **32** | OMe | (*S*)-2-(2,2-difluorocyclohexyl) | 0.2 | >100 | 0.32 | 93 | <0.5 | <0.5 |
| **33** | OMe | (*R*)-2-(2,2-difluorocyclohexyl) | 0.9 | 64 | 3.2 | 32 | <0.5 | <0.5 |
| **34** | OEt | (1*R*,2*S*)-2-aminocyclohexyl | 2.2 | >100 | >100 | >100 | <0.5 | 0.9 |
| **35** | OMe | 2,2-dimethylcyclohexyl | 0.3 | 27 | 2.5 | 15 | 3.4 | 6.2 |
| **36** | OEt | (1*R*,2*S*)-2-hydroxycycloheptyl | 0.2 | >100 | 0.63 | >100 | <0.5 | 0.6 |
| **37** | OMe | (1*R*,2*S*)-2-hydroxycycloheptyl | 0.1 | 60 | 0.25 | 60 | <0.5 | <0.5 |
| **38** | OMe | (*S*)-2-(2,2-difluorocycloheptyl) | 0.1 | >100 | 0.16 | >100 | <0.5 | 0.8 |

See Table 1 for explanation.

Initially, the importance of the alkyl ring at R_1_ was assessed by replacing it with a phenyl group (**23**), although the replacement was tolerated there was a reduction in potency against both LysRS and bacterial growth. The combination of a methoxy at R_2_ and the *cis* 2-hydroxy cyclohexyl at R_1_ (**24**) had good potency against LysRS and bacterial growth but only modest selectivity against KARS1 and HepG2 cytotoxicity (analogous to the unsubstituted cyclohexyl (**8**)). To gain selectivity, the ethoxy equivalent (**25**), was synthesised, this retained a promising potency profile and was found to be >100-fold selective over KARS1 and HepG2. A crystal structure of **25** bound in the LysRS active site (PDB 9qei), showed that, as anticipated the polar hydroxyl group was able to establish interactions with Glu422 within the ATP binding pocket (**Figure 3A**). Although this extra interaction did not increase potency, the addition of the 2-hydroxy improved the metabolic stability vs. **10** (**Table 3**). Following the successful addition of a hydroxyl group at the two-position of the cyclohexyl ring, it was subsequently examined at the 3- and 4-positions with a methoxy at R_2_. The 3-hydroxyl (**26**) was slightly less active against both LysRS and KARS1. At the 4-position, the hydroxyl (**27**) retained potency against LysRS and gained KARS1 selectivity. The crystal structure of **27** (PDB 9qbr) confirmed that the 4-hydroxyl group was making a direct hydrogen bond with Thr425 in LysRS (**Figure S2A**). In KARS1, the equivalent residue is Asn497, which forms hydrogen bonds with the bound substrate lysine and Arg485, thereby blocking access to this part of the pocket. In LysRS, Thr425 is too distant from the bound substrate lysine and Lys413 (Arg485 in KARS1) to establish this hydrogen bond network, thereby allowing access of **27** and explaining the gain in selectivity. The crystal structure of **27** indicated that there was space to extend the hydroxy group deeper into the pocket. In a bid to develop this further, to explore the potential for interaction at the 4-position, a carbon spacer was added (**28**); again, this displayed good LysRS potency, but the slight decrease in MIC potency, for both **27** & **28**, prevented further scope for optimisation (**Table 1**).

**Figure 3:**
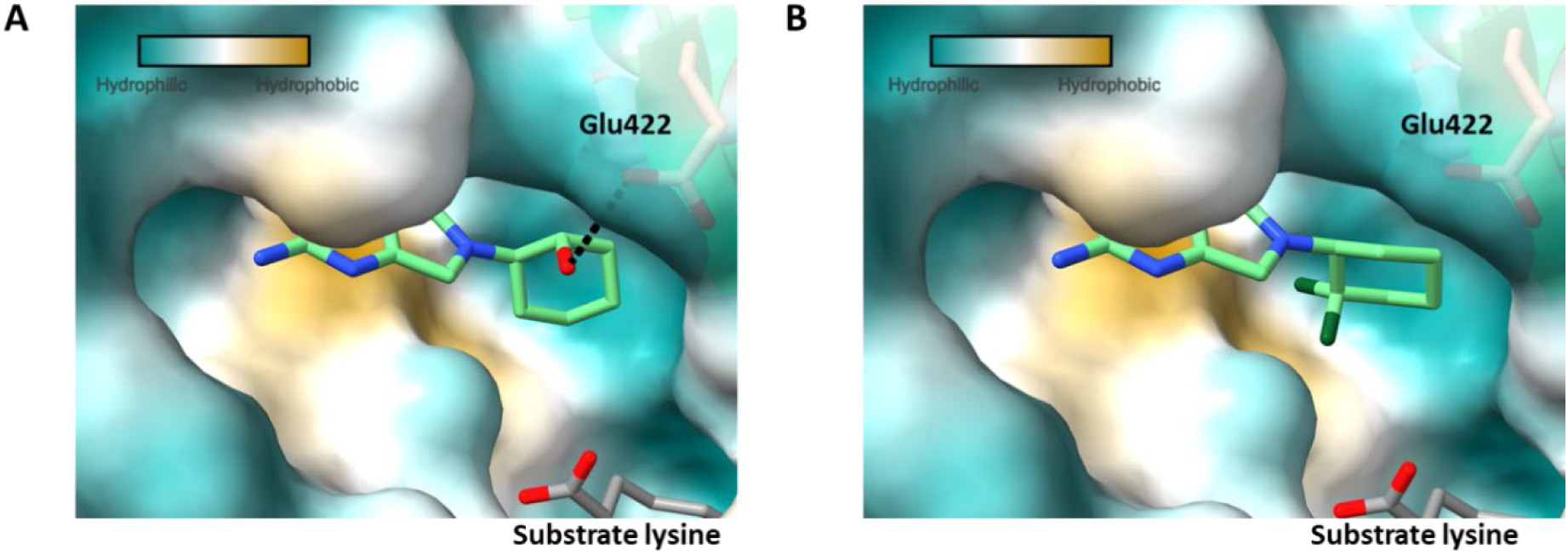
Surface views of inhibitors in the ATP binding pocket of LysRS. Figure prepared using UCSF ChimeraX^16^ and coloured using the Kyte-Doolittle scale for hydrophobicity^17^. **A.** The polar hydroxyl group of **25** sits in the more hydrophilic part of the pocket making a hydrogen bond with Glu422 (PDB 9qei). **B.** The hydrophobic di-fluoro group of **32** sits in the hydrophobic area formed by the side chain of Met271 (yellow surface) (PDB 9qdj).

Adopting a similar strategy to block metabolic vulnerabilities, the addition of fluorine at each position on the cyclohexyl ring was explored. To prevent the formation of complex diastereoisomers each position was disubstituted with fluorine. At first, ethoxy at R_2_ was combined with 2,2-, 3,3- and 4,4-difluorocyclohexyl at R_1_ (**29** - **31**); for these initial explorations, racemic 2,2- and 3,3-difluorocyclohexyl were employed. Substitution was tolerated, at all positions and a good *in vitro* metabolic profile was observed. However, they all displayed a decrease in activity in the MIC assay, compared to unsubstituted **10** (**Table 3**). Next, the individual *R* and *S* enantiomers of the 2,2-difluorocyclohexyl substituted cyclohexyl with a methoxy at R_2_ were prepared. The *R* enantiomer (**33**) had similar properties to the bare cyclohexyl derivative (**8**), with potent activity but limited selectivity over KARS1 and HepG2. Interestingly, the *S* enantiomer (**32**) was not only potent but also the first methoxy R_2_ substituted compound to show >150-fold selectivity at both the enzyme and cellular level. The bound crystal structure of **32** (PDB 9qdj) showed that the difluoro group was oriented towards Met271 at the rear of the ATP pocket (**Figure 3B**); this residue forms a hydrophobic region in the pocket, favourable for interacting with the hydrophobic difluoro group. The corresponding residue in KARS1 is a smaller Thr, significantly altering the proximity and electrostatic potential of this part of the pocket, thereby explaining the gain in selectivity.

Given the improved properties of **32** & **25**, they were selected for profiling in the acute efficacy model. Clear reductions in bacterial load were achieved for both compounds (**Figure 2**). **32** displayed a similar reduction to **8** (2.6 Log_10_ reduction), while **25** generated a 3.1 Log_10_ reduction. The later representing a clear cidal effect on the model, reducing the bacterial load to below that at the start of treatment. The PK profile of **25** (**Table S2**) showed that the bioavailability and exposure (AUC) was significantly improved compared to the unsubstituted **10**, and the V_d_ was modestly increased. To explore the efficacy of **25** in more depth an *in vivo* dose response study was performed, the ED_99_ (dose that causes a 2 Log_10_ reduction in CFU with respect to untreated mice) was 49 mg/kg^14^. The improved PK profile and efficacy of **25** were sufficient to mark the transition from hit to lead to lead optimisation.

### Lead Optimisation

After demonstrating that the enhanced PK profile of **25** led to improved efficacy, further SAR were conducted with a particular focus on improving the V_d_ and ultimately the compounds half-life. Several strategies were employed, the first was to evaluate if a basic group would be tolerated in the R_1_ pocket. Utilising the same stereochemical configuration as **25** (1*R*,2*S*)-2-aminocyclohexyl (**34**) was examined in the R_1_ position and found to display a 7-fold reduction in enzymatic potency compared to **25**, surprisingly this was accompanied by a >45-fold differential in potency between the enzymatic and MIC assays, an anomaly for the series. This was hypothesised to be due to the presence of four HBDs, resulting in poor permeability into the bacteria (**Table 3**). Although not progressible further, evaluation of **34** in a PK study demonstrated that the added basicity resulted in a significant increase in both the V_d_ and half-life (**Table S2**). A second approach to improve V_d_ focussed on the addition of discrete lipophilicity by the introduction of 2,2-dimethylcyclohexyl at R_1_ (**35**). Although tolerated, it lost the selectivity observed with 2,2-difluoro (**32**) (**Table 3**). A third strategy, again aimed at subtly increasing lipophilicity, involved taking the most effective R_1_ substituted cyclohexyls and increasing the ring size to cycloheptyl (**Table 3**); the larger ring, not only increased lipophilicity but was also thought to optimally fill the R_1_ pocket, displacing multiple waters that existed in the apo structure. This strategy afforded highly potent molecules against LysRS, each approaching the tight binding limit for the assay (IC_50_ = 0.1 µM).

When ethoxy was at R_2_, the cycloheptyl (**36**) displayed a good overall profile but offered little advantage over the cyclohexyl equivalent (**25**). Methoxy at R_1_ in combination with (1R,2S)-2-hydroxycycloheptyl at R_2_ (**37**) had a modest (4-fold) improvement in selectivity over KARS1 compared to the cyclohexyl equivalent (**24**). Encouragingly, as anticipated from its cyclohexyl equivalent (**32**) when methoxy at R_2_ was combined with (*S*)-2-(2,2-difluorocycloheptyl) (**38**), this retained potency and improved its selectivity profile (>1000-fold over KARS1 and >500-fold at a cellular level).

Having demonstrated significant improvements, **37** & **38** were evaluated in the acute efficacy model. Both compounds were evaluated at 50 mg/kg, the “benchmark” ED_99_ activity for **25**^14^. Despite being more potent at the enzyme and cellular levels, **38** was less efficacious than **25** resulting in only a 1.3 Log_10_ reduction in bacterial load (**Figure 4**). But, encouragingly, **37** was significantly more potent than **25** with a 3 Log_10_ reduction in bacterial load (**Figure 4**). The PK profile for **37** was assessed, compared to **25**, it had improved half-life and V_d_, while bioavailability and AUC were comparable if not marginally better (**Table S2**). The improved PK profile led to an improved efficacious effect for **37** but, because of its limited specificity over KARS1, **37** was deemed not suitable for further progression. In summary the extended assessment of the R_1_ SAR had produced notably more potent molecules, with a greatly enhanced selectivity profile, that demonstrated a cidal *in vivo* profile. To examine additional opportunities for improving the observed efficacious effect, the SAR evaluation was expanded to explore R_3_ and R_4_ (**TOC Figure**).

**Figure 4:**
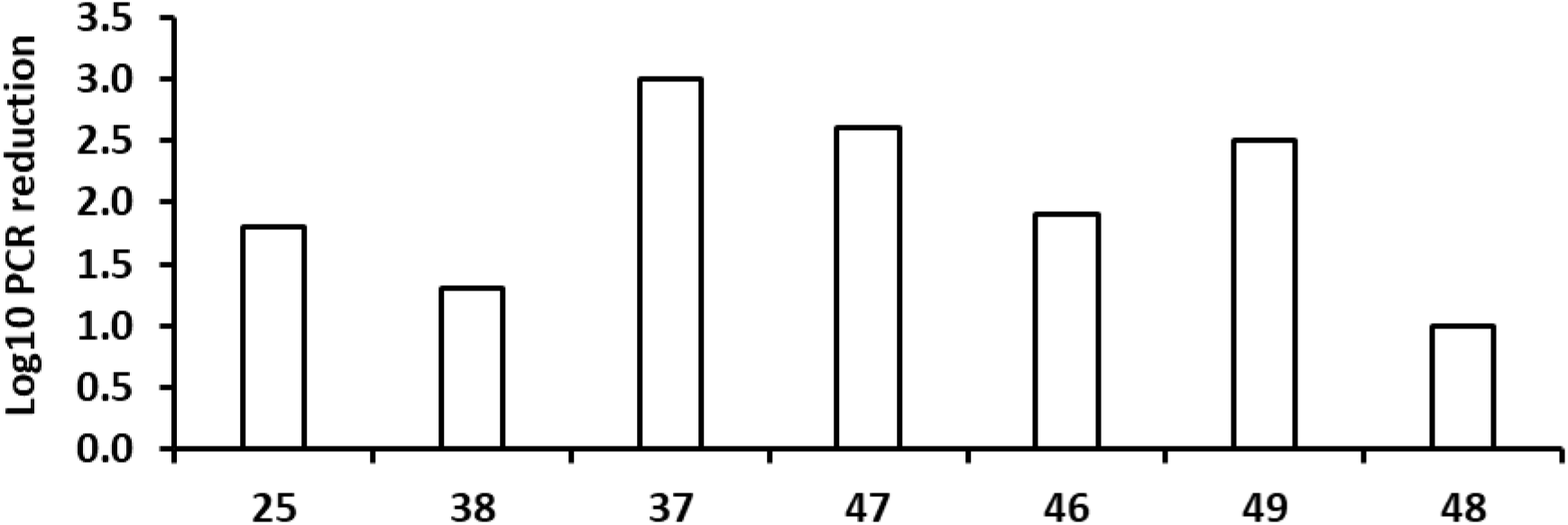
*In vivo* efficacy of advanced LysRS inhibitors. Compounds **25**, **38**, **37**, **47**, **46**, **49** and **48** were tested in an acute model of infection in C57BL/6mice. Dosing (@50 mg/kg the ED99 for **774**) started 1 day after infection and lasted for 8 days; each group consisted of two mouse/dose. The effect on bacterial load was monitored by PCR detection of bacterial DNA.

#### Development of R_3_ pocket SAR

Evaluation of crystal structures generated from the series, such as **37** (PDB 9qc3) highlighted a water network adjacent to the benzylic carbon on the R_3_ core (**Figure S2B**). This provided an opportunity to explore a different substitution vector with the potential to modulate physicochemical properties (**Table 4**). In a bid to further introduce discreet lipophilicity, the benzylic hydrogens were replaced with methyl (**39**); this was not well tolerated, with a dramatic loss in potency in the LysRS assay and correspondingly against *M. tuberculosis* growth. Replacing the benzylic carbon with NH or NMe, (**40** & **41** respectively), also reduced activity against LysRS. Analysis of the crystal structure indicated that it might be possible for a substituent to interact with Glu422, (as observed with the 2-hydroxy on **25**). An extended azetidine (**42**) was synthesised, and an interaction was confirmed by crystallography (**Figure S2C** PDB 9qc4). **42** possessed reasonable LysRS activity, but this did not translate into cellular potency, potentially again due to an impact on bacterial permeability from the additional HBD.

**Table 4:** Core changes.

| Compound ID | Structure | LysRS<br>IC<sub>50</sub><br>(μM)<sup>a</sup> | KARS1<br>IC<sub>50</sub><br>(μM)<sup>b</sup> | MIC<br>(μM)<sup>c</sup> | HepG2<br>(μM)<sup>d</sup> | Clearance<br>(mL/min/g) | |
| --- | --- | --- | --- | --- | --- | --- | --- |
|  |  |  |  |  |  | Mics<sup>e</sup> | Heps<sup>f</sup> |
| **39** |  | 21 | >100 | >100 | >100 | <0.5 | ND |
| **40** |  | 12 | >100 | ND | >100 | <0.5 | ND |
| **41** |  | 8 | >100 | ND | >100 | 39 | ND |
| **42** |  | 1.5 | 56 | 50 | >100 | <0.5 | <0.5 |

See Table 1 for explanation.

**Table 5:** R_4_ changes.

| | R<sub>4</sub> | R<sub>2</sub> | R<sub>1</sub> | LysRS<br>IC<sub>50</sub><br>(μM)<sup>a</sup> | KARS1<br>IC<sub>50</sub><br>(μM)<sup>b</sup> | MIC<br>(μM)<sup>c</sup> | HepG2<br>(μM)<sup>d</sup> | Clearance<br>(mL/min/g) | |
| --- | --- | --- | --- | --- | --- | --- | --- | --- | --- |
|  |  |  |  |  |  |  |  | Mics<sup>e</sup> | Heps<sup>f</sup> |
| **43** | methyl | OEt | cyclohexyl | 1.3 | >100 | 7.9 | >100 | 12 | ND |
| **44** | cyclopropyl | OMe | cyclohexyl | 0.6 | >100 | 0.40 | >100 | 9.3 | ND |
| **45** | cyclopropyl | OMe | (1*R*,2*S*)-2-hydroxycyclohexyl | 0.4 | >100 | 1.6 | >100 | 1.1 | 3.6 |

See Table 1 for explanation.

#### Evaluation of R_4_ pocket SAR

The final area of SAR explored was to reduce the number of HBDs by capping the R_4_ nitrogen. The addition of methyl (**43)**, resulted in a reduction in both LysRS and MIC potency. In contrast, cyclopropyl (**44**) retained good LysRS inhibitor but had a detrimental effect on MIC potency and microsomal turnover. To mitigate this metabolic instability, (1*R*,2*S*)-2-hydroxycyclohexyl was examined as the R_1_ substituent, to decrease the lipophilicity (**45**); this had a good overall profile, including an acceptable clearance profile, but ultimately the potency was not in the desired range. In summary modifying R_3_/R_4_ did not improve the overall profile of the compound, so attention focussed on combining the key R_1_/R_2_ SAR changes.

#### Combination of key SAR features

Analysis of the SAR generated highlighted clear trends within the series: 1) Methoxy at R_2_ consistently resulted in an improved MIC profile compared to ethoxy; 2) Ethoxy at R_2_ provided better selectivity over KARS1 than the methoxy; 3) Cycloheptyl at R_1_ boosted both enzymatic and cellular potency compared to cyclohexyl; 4) (*S*)-2-(2,2-difluorocycloalkyls at R_1_ improved selectivity over KARS1; 5) (1*R*,2*S*)-2-hydroxycycloalkyls at R_1_ significantly improved bioavailability and exposure, highlighting the importance of the hydroxyl substituent for the overall pharmacokinetic profile. Comparing the crystal structure of **25** with **32**, it was evident that the 2-hydroxy and 2,2-difluoro substituents on the R_1_ hexyl group were orientated towards different parts of the pocket (**Figure 3**). The 2,2-difluoro in **32** pointed towards residue Met271 while the 2-hydroxy of **25** aligned in the opposite direction towards residue Glu422. Based on the trends seen and this structural information, combination molecules were designed and prepared, with methoxy/ethoxy at R_2_ and 2-((1*S*,6*S*)-2,2-difluoro-6-hydroxycyclohexyl) or 2-((1*S*,7*S*)-2,2-difluoro-7-hydroxycycloheptyl) at R_1_ (**Table 6**). All the combination molecules (**46** - **49**) were very potent against LysRS, reaching the tight binding limit for the assay. Subsequently, to decrease the tight binding limit, the assay was reconfigured, so *in vitro* potencies could be differentiated (increased ATP concentration, decreased amount of LysRS and extended incubation time). In the reconfigured assay, all four molecules were very potent against LysRS with the methoxy/cycloheptyl (**49**) having an *in vitro* IC_50_ potency of 50 nM (**Table 6**). The crystal structure of **49** was solved^14^ and the surface view in the ATP binding pocket of LysRS is shown in the supplementary information (**Figure S3**). As hypothesised, in this combined molecule, the 2,2-difluoro pointed towards the hydrophobic region around Met271, while the 2-hydroxy aligned in the opposite direction towards residue Glu422. All four molecules displayed an excellent selectivity profile against KARS1, with none of them showing any significant inhibition up to the maximum 100 µM tested. As the program advanced, **49** was tested further against KARS1 to determine the true IC_50_, which equated to 1.2 mM, reflecting an enzyme selectivity of ∼25,000-fold. The excellent LysRS potency was mirrored in the antibacterial activity, with all four molecules achieving <1 µM in both extra and intracellular MIC assays. Again, **49** was found to be the most potent with an MIC of 40 nM and a cellular selectivity over HepG2 cytotoxicity of >2,500-fold (**Table 6**).

**Table 6:**
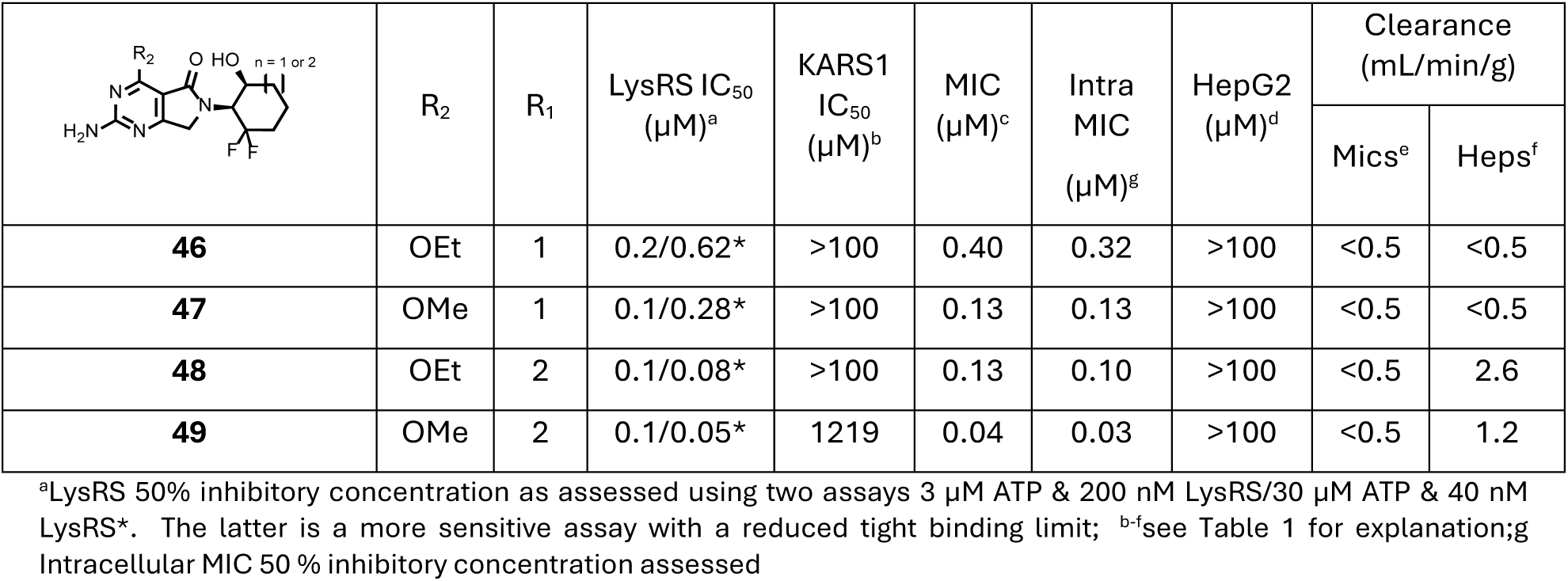
Combination molecules (low ATP/high ATP)

|  | R<sub>2</sub> | R<sub>1</sub> | LysRS IC<sub>50</sub><br>(µM)<sup>a</sup> | KARS1<br>IC<sub>50</sub><br>(µM)<sup>b</sup> | MIC<br>(µM)<sup>c</sup> | Intra<br>MIC<br>(µM)<sup>g</sup> | HepG2<br>(µM)<sup>d</sup> | Clearance<br>(mL/min/g) |  |
| --- | --- | --- | --- | --- | --- | --- | --- | --- | --- |
|  |  |  |  |  |  |  |  | Mics<sup>e</sup> | Heps<sup>f</sup> |
| **46** | OEt | 1 | 0.2/0.62* | >100 | 0.40 | 0.32 | >100 | <0.5 | <0.5 |
| **47** | OMe | 1 | 0.1/0.28* | >100 | 0.13 | 0.13 | >100 | <0.5 | <0.5 |
| **48** | OEt | 2 | 0.1/0.08* | >100 | 0.13 | 0.10 | >100 | <0.5 | 2.6 |
| **49** | OMe | 2 | 0.1/0.05* | 1219 | 0.04 | 0.03 | >100 | <0.5 | 1.2 |

<sup>a</sup>LysRS 50% inhibitory concentration as assessed using two assays 3 µM ATP & 200 nM LysRS/30 µM ATP & 40 nM LysRS\*. The latter is a more sensitive assay with a reduced tight binding limit; <sup>b-f</sup>see Table 1 for explanation; <sup>g</sup>Intracellular MIC 50 % inhibitory concentration assessed

The four molecules were progressed to an acute murine efficacy model at 50 mg/kg, except for **48** which was dosed at 39 mg/kg due to compound limitation (**Figure 4**). Both compounds with methoxy at R_2_ were very efficacious at this dose, with a 2.6 & 2.5 Log_10_ reduction for **47** & **49** respectively. The analogues with an ethoxy at R_2_ were less efficacious, although both still reduced bacterial load significantly. Over the course of the efficacy experiment, limited PK samples were taken, to obtain a preliminary estimate of compound exposure (**Table S3**). Only **49** had detectable levels in mouse blood at 24 h post dosing. Given its *in vitro/in vivo* potency and that it had the best *in vivo* exposure, **49** was selected to go into a dose response acute efficacy model. The result was reported previously^14^ with **49** achieving an ED_99_ @12 mg/kg. Since in the initial 50 mg/kg model **47** had a similar impact on bacterial load to **49**; once the ED_99_ was established at 12 mg/kg, **47** was further tested at that dose. In that experiment, when compared to the bacterial load in the untreated mice, **47** only achieved a 0.8 Log_10_ CFU reduction. Consequently, **49** was selected as the preferred molecule to move into additional preclinical studies.

### Biology

#### Mode of Inhibition

The crystal structures had confirmed that the compounds from this series were occupying the ATP pocket of LysRS. In the reconfigured biochemical assay, shifts in IC_50_ were observed when employing different ATP concentrations; weaker potencies in assays at high ATP concentrations, suggesting that the series was acting as a competitive inhibitor towards ATP. Next, the mode of inhibition with respect to lysine was explored. To address this, LysRS reactions were performed at a range of concentrations of lysine with **11** & **49**. As the lysine concentration increased, the apparent potency of the compounds increased (**Figure 5**); comparing assays at 12 µM vs. 0.5 µM lysine, the observed IC_50_ shifts were ∼7-fold & ∼6-fold for **11** & **49**, respectively. The subsequent IC_50_ against the substrate concentration/Km ratio ([S]/Km) plots were characteristic for this series being uncompetitive with lysine^18^ (**Figure 5**). Uncompetitive inhibition occurs when an enzyme with two substrates (lysine and ATP) binds them in a specific sequence (in this case lysine binds before ATP). In this scenario, when lysine is already bound to LysRS, inhibitors from this series exhibit stronger binding in the ATP pocket; consequently, higher lysine concentrations increase the apparent potency of the inhibitors.

**Figure 5:**
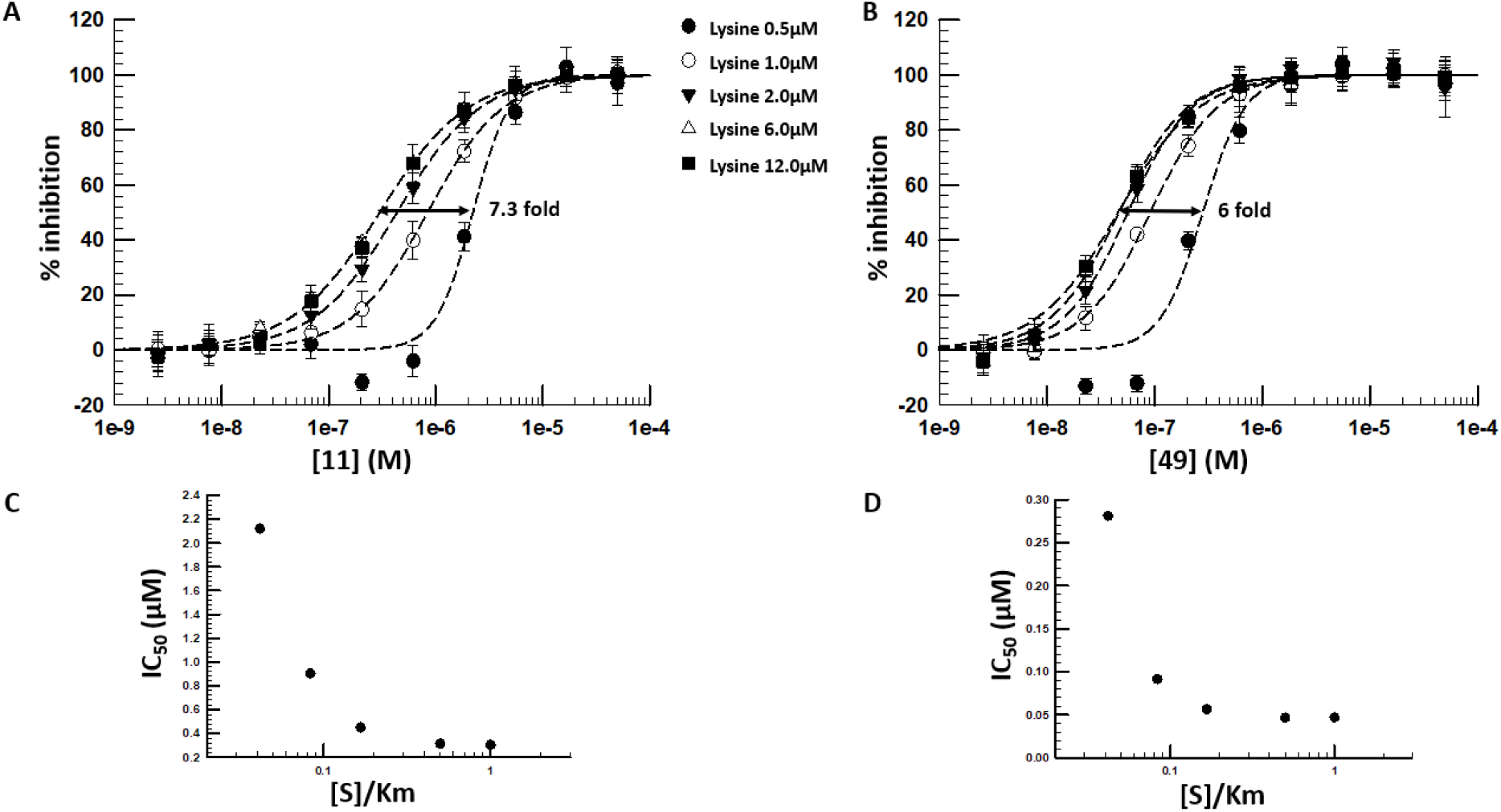
Mode of inhibition studies. Determination of shifts in IC50 for **11** (**A**) and **49** (**B**) at different lysine concentrations. The effect of [S]/Km on Inhibition (IC50) showing tighter inhibition (lower IC50 values) at higher lysine concentrations (**C** & **D**) indicate that the series acts as uncompetitive inhibitors with respect to lysine binding^18^. The data is the average ±standard deviation of four technical repeats and is a representative of four different experiments.

#### Cellular Mode of Action (MoA)

The initial MoA studies for this series, including resistant mutant generation, overexpression studies and metabolomics were presented previously ^14^. Here this analysis was extended by constructing an H37Rv strain, Mtb *lysS*-TetON, in which transcription of *lysS* is repressed by a tetracycline repressor, TetR. In Mtb *lysS*-TetON, transcription of *lysS* is induced with anhydrotetracycline (ATc) and repressed when grown without ATc. **11** was chosen as a representative for the series to demonstrate that induction of LysRS expression (with Atc) decreased growth inhibition and repression of LysRS expression (achieved by cultivation of Mtb *lysS*-TetON without Atc) increased growth inhibition by this compound (**Figure 6**). Susceptibility of Mtb *lysS*-TetON to growth inhibition by isoniazid was similar to that of WT H37Rv with and without ATc, demonstrating that silencing of *lysS* transcription caused specific susceptibility to growth inhibition by LysRS inhibitors.

**Figure 6:**
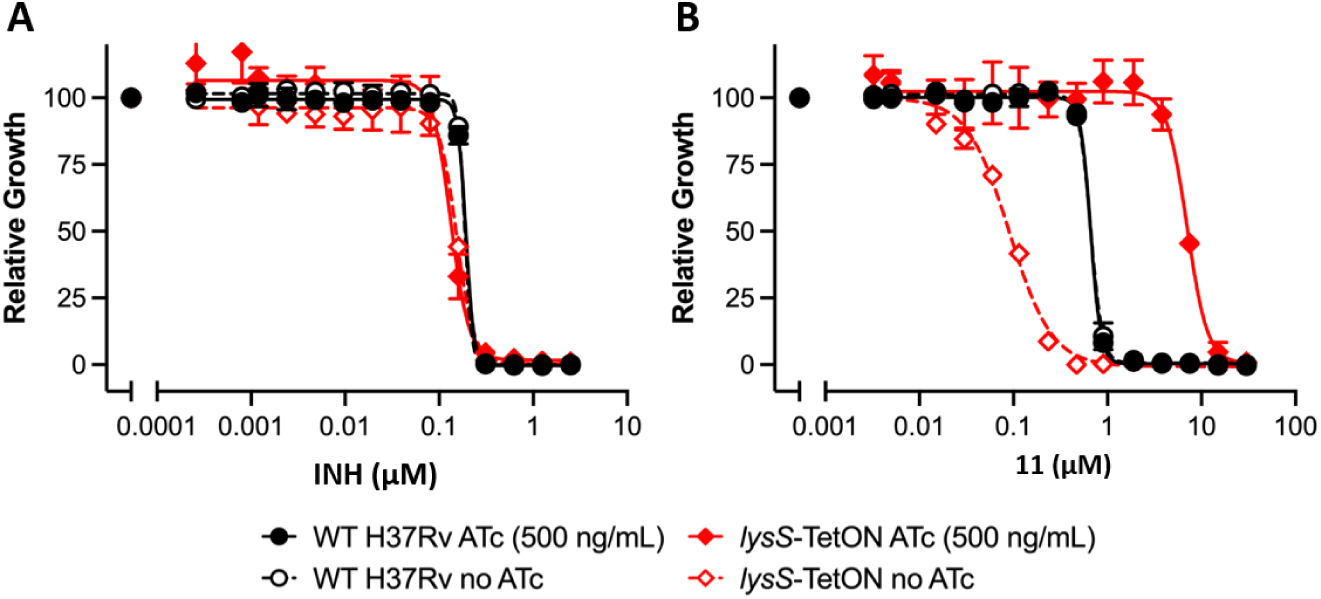
Cellular mode of action is dependent on LysRS. Dose response curves for isoniazid and **11** tested against WT Mtb H37Rv and Mtb *lysS*-tetON grown with and without ATc. Data are averages of two or three cultures and representative of two independent experiments.

### Chemistry

The syntheses of target compounds **2**-**5** are summarised in **Scheme 1**. Starting material **S1**^14^ was reacted with the relevant R_1_ amine, under basic conditions to deliver target compounds (**2**-**5**).

**Scheme 1:**
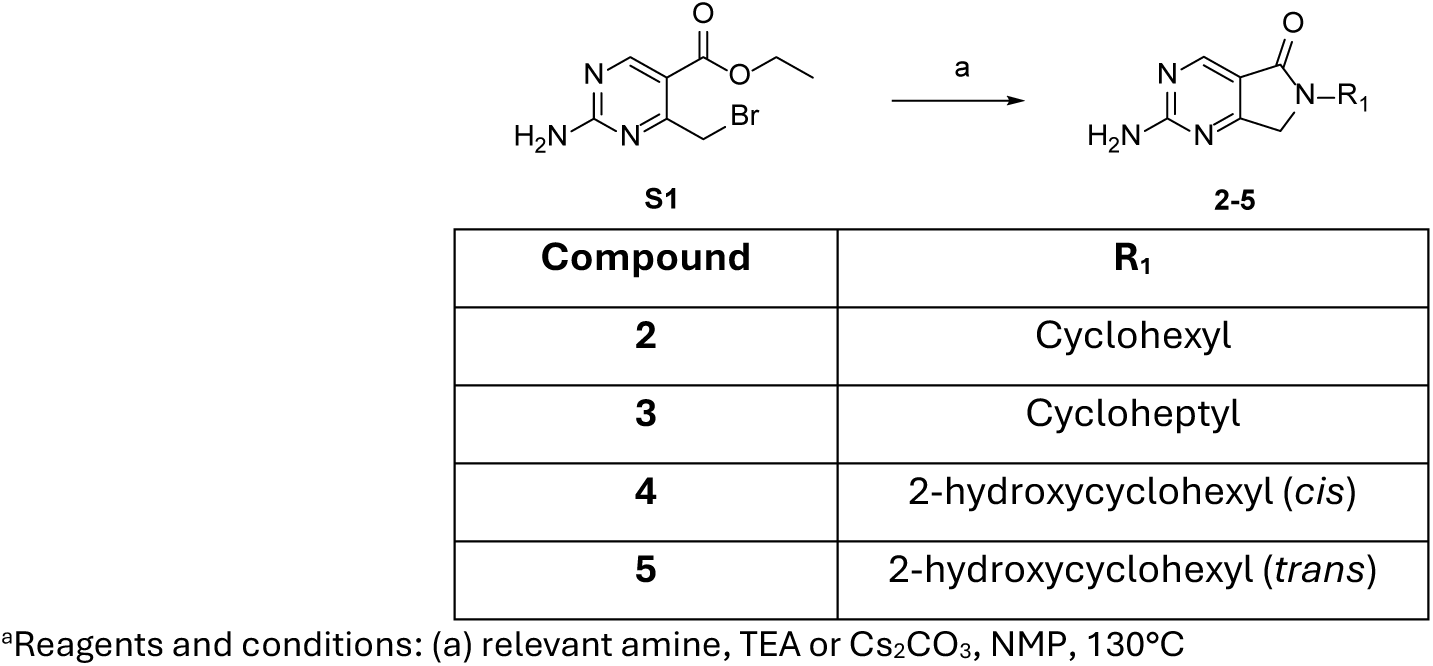
Synthesis of target compounds 2 – 5^a^.

**Scheme 2** summarises the syntheses of common ethyl 4-chloro-6-(difluoromethoxy)-2-(methylthio)pyrimidine-5-carboxylate intermediates **S6 – S10** that afforded aldehydes **S11-S15.** Multiple routes were utilised to deliver intermediates **S6 – S10** over the course of investigation into the series. Firstly, **S16** was reacted with vinylmagnesium bromide in the presence of (diacetoxyiodo)benzene to afford **S14**. Secondly, ethyl 4,6-dichloro-2-(methylthio)pyrimidine-5-carboxylate (**S2**) was reacted with the relevant R_2_ alcohol, deprotonated by sodium hydride to afford **S3 – S5**, subsequent cross coupling with potassiumtrifluoro(vinyl)boranuide yielded (**S11 – S13**). Finally, where the R_2_ substituent was difluoromethoxy, 6-chloro-2-(methylthio)pyrimidin-4(3*H*)-one (**S17**) was reacted with sodium chlorodifluoroacetic acid in the presence of sodium carbonate to afford **S18**, the ester functionality was introduced by reaction with *n*-butyllithium, 2,2,6,6-tetramethylpiperidine and carbonochloridate to afford (**S19**). Universally intermediates **S6 – S10** were reacted with osmium tetroxide (4 wt.% in water), 2,6-lutidine and sodium periodate to deliver the relevant aldehyde (**S11 – S15**) varying R_2_ at the 6-position as shown in **Scheme 2**.

**Scheme 2:**
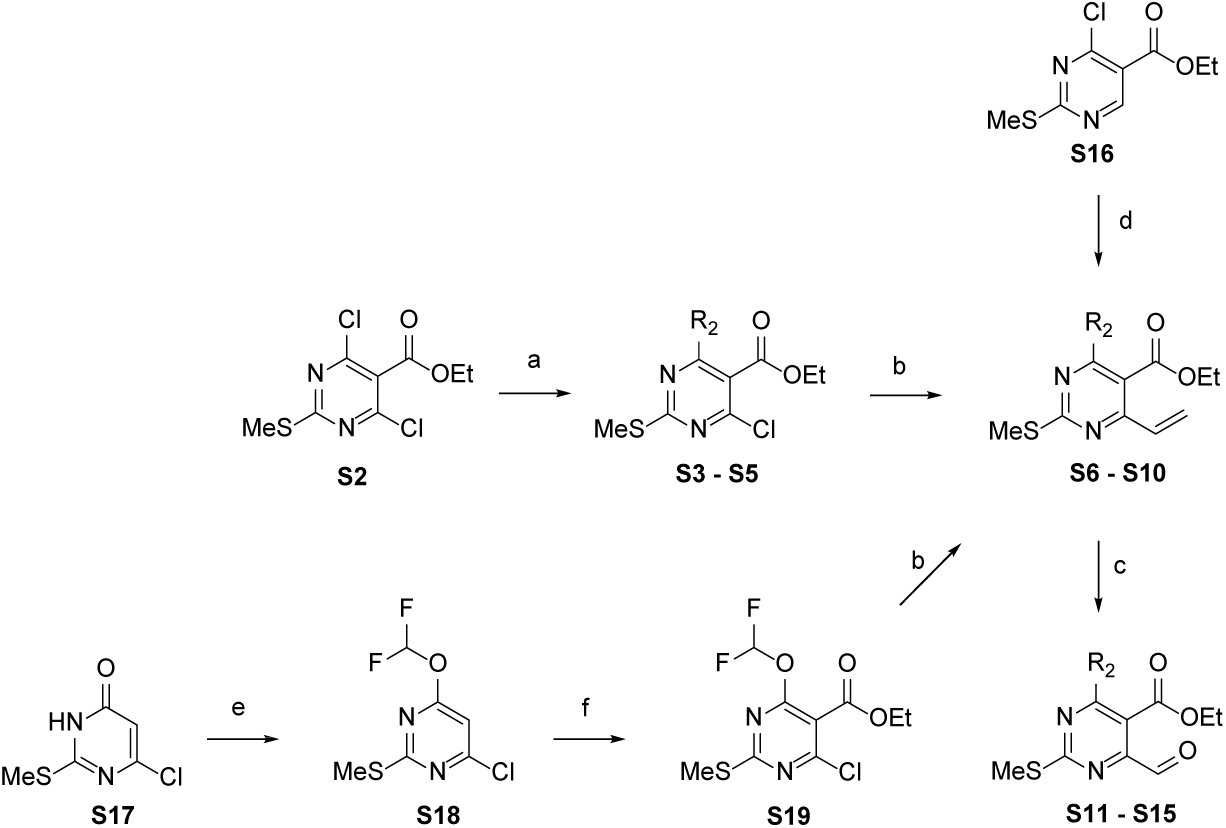
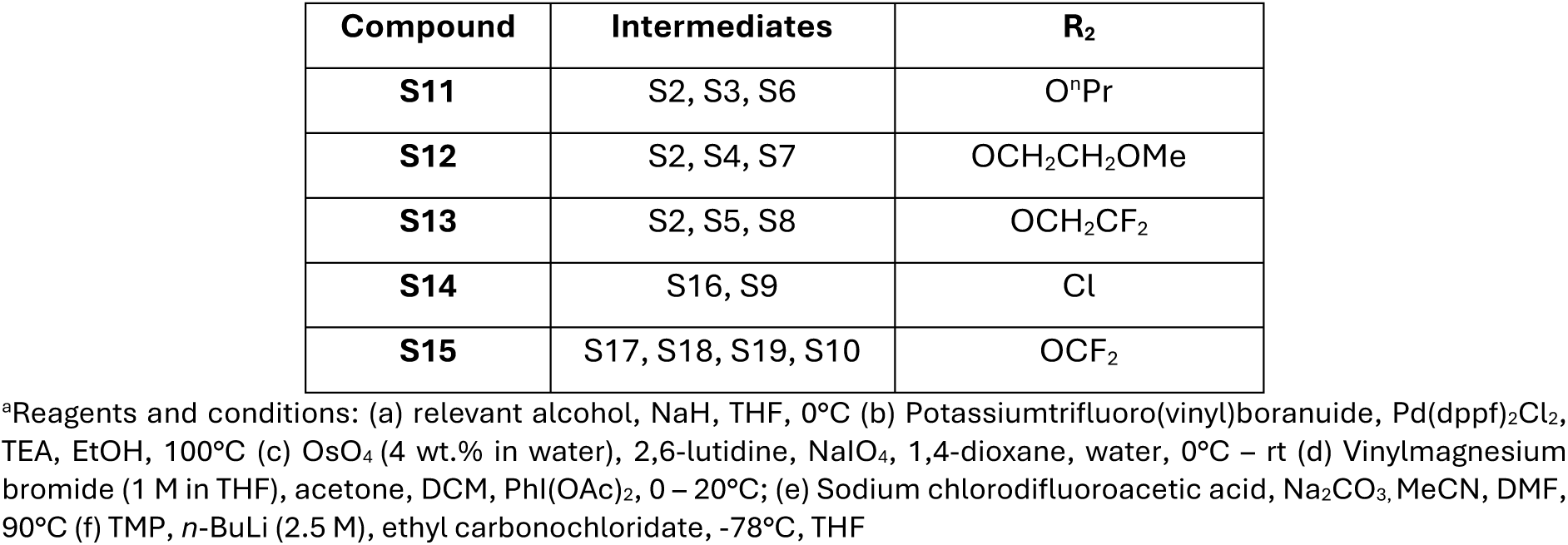
Synthesis of intermediate S11 - S15.

**Scheme 3** summarises the synthesis of target compounds **12 - 14, 16, 17, 21, 23, 26 –31, 34, 35, 38, 39, 43 – 45 and 48**. To afford most target compounds, a reductive amination was carried out on the relevant R_1_ amine and corresponding intermediate **S11-13, S15** and **S20 – S21** utilising sodium triacetoxyborohydride to provide intermediate 2-(methylthio)-6,7-dihydro-5*H*-pyrrolo[3,4-*d*]pyrimidin-5-one substituted at the 6-position with R_1_ and the 4-positon with R_2_ **S22-36**. Oxidation of the methylthiol was achieved with either oxone or *m*-CPBA, subsequent displacement with ammonia (0.5 M in 1,4-dioxane) or the R_4_ relevant amine yielded target compounds (**12**,**14**,**16**,**17**,**21**, **23, 29**-**31**,**36**,**43**-**45** and **48**). Alternatively, a reductive amination was carried out methyl 2-((*tert*-butoxycarbonyl)amino)-4-formyl-6-methoxypyrimidine-5-carboxylate (**S37**) in an analogous manner to form **S38-S42** with the appropriate R_1_ substitution at the 6-position. Removal of the Boc group afforded desired compounds (**26**-**28, 35** and **38**).

Disubstitution on the benzylic position of the lactam ring with R_3_ substituents was achieved by deprotonation of 2-(methylthio)-6,7-dihydro-5*H*-pyrrolo[3,4-*d*]pyrimidin-5-one substituted where R_1_ is (1*S*,4*S*)-4-hydroxycyclohexyl and R_2_ is methoxy with sodium hydride and addition of methyl iodide to yield **S43**, the target compound (**39**) was obtained as described previously by oxidation and subsequent displacement with ammonia. Ether cleavage of the methyl of **14** with boron tribromide afforded target compound **13**. Boc deprotection of **S31** provided target compound **34**.

**Scheme 3:**
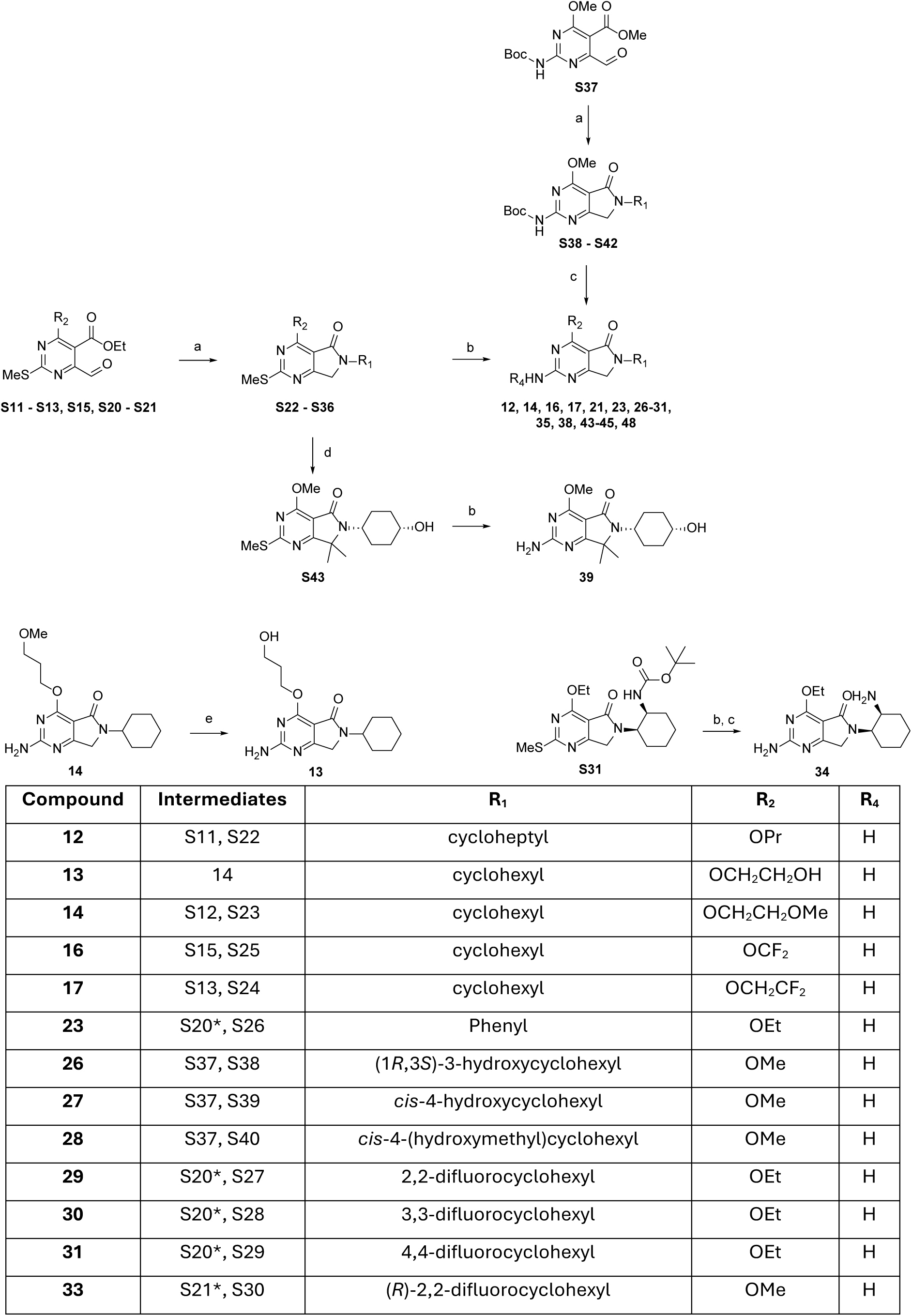
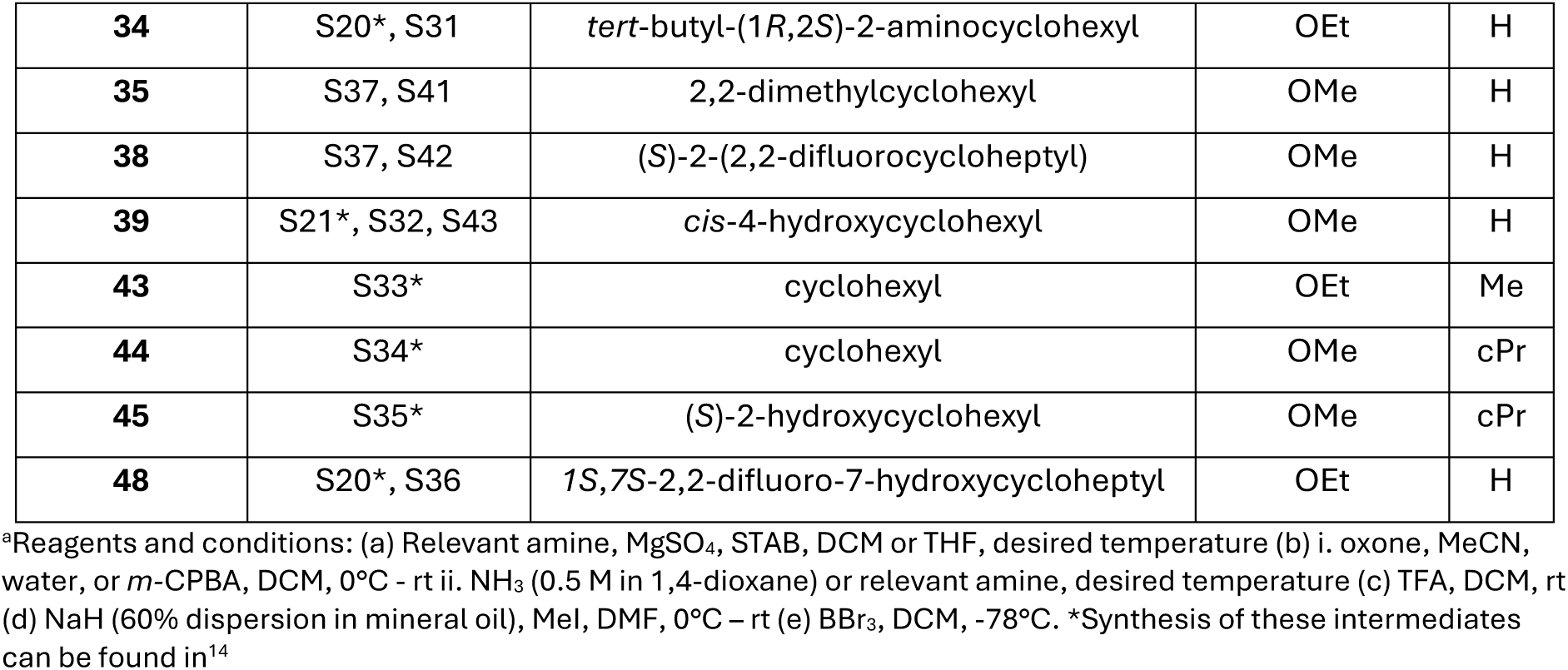
Synthesis of target compounds 12 - 14, 16, 17, 21, 23, 26 – 31, 34, 35, 38, 39, 43 – 45 and 48.

**Scheme 4:**
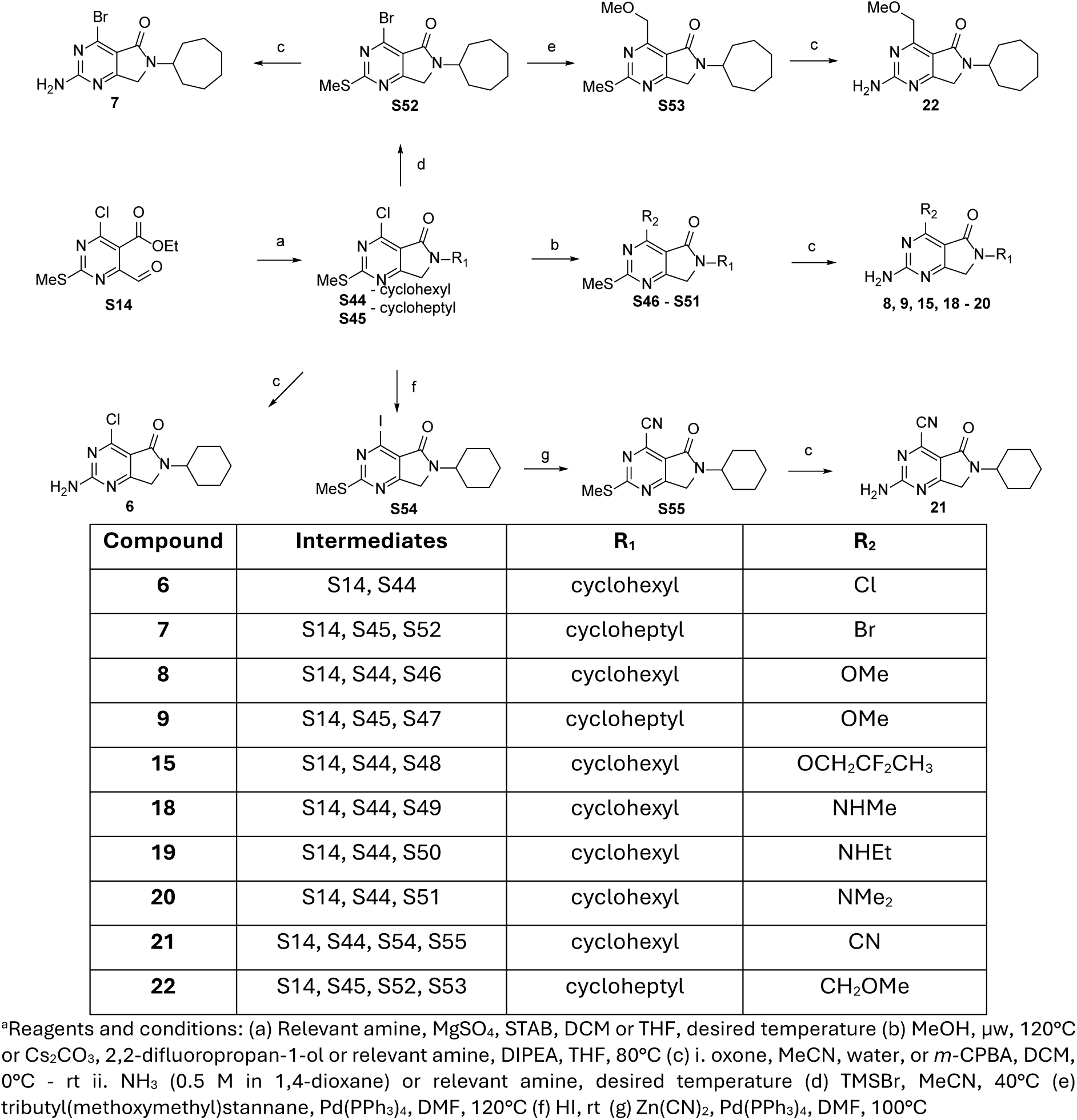
Synthesis of target compounds 6 – 9, 15, 18 - 22.

The synthesis of target compounds **6** – **9**, **15**, **18** – **22** is summarised in **Scheme 4**. Following a similar procedure to that described in scheme 3, a reductive amination with **S14** with cyclohexylamine and cycloheptyl amine yielded **S44** and **S45** respectively. The chlorine at R_1_ was displaced with the relevant amine or 2,2-difluoropropan-1-ol to yield **S46**-**51** that were converted into the corresponding target products (**8**,**9**,**15**,**18**-**20**) by oxidation of the methylthiol and displacement with ammonia. Reaction of **S44** with bromotrimethylsilane in acetonitrile exchanged the chloro at R_1_ for a bromo **S52**. Reaction of **S52** with tributyl(methoxymethyl)stannane in a palladium catalysed reaction introduced a C-linked methoxymethyl at R_1_ (**S53**). Both **S52** and **53** were converted to the corresponding target compounds **7** and **22** by oxidation of the methylthiol and displacement with ammonia. In an analogous manner **S44** yielded target compound **6**. Reaction of **S44** with hydriodic acid formed **S54**, reaction of this intermediate with zinc cyanide formed **S55**, the amino group was introduced to yield target compound **21**.

The synthesis of target compounds **40**-**42** is outlined in **Scheme 5**, ethyl 4-chloro-6-ethoxy-2-(methylthio)pyrimidine-5-carboxylate (**S56**) underwent aromatic nucleophilic substitution (SNAr) with *tert*-butyl *N*-amino-*N*-cycloheptyl-carbamate under basic conditions to yield **S57**, removal of the Boc group under acidic conditions afforded **S58**. Aza ring formation was achieved by reaction of S58 in the presence of potassium hydroxide to yield **S59** as a common intermediate. Oxidation of **S59** in a similar to manner as described in Scheme 3 afforded target compound **40**. Reaction of **S59** with the appropriate amine and potassium carbonate delivered intermediates **S60** and **S61**. **S60** was oxidised as previously described to afford 6-amino-2-cycloheptyl-1-(cyclopropylmethyl)-4-ethoxy-1,2-dihydro-3*H*-pyrazolo[3,4-*d*]pyrimidin-3-one (**41**). **S61** was oxidised and the Boc group removed under acidic conditions to yield target compound **42**.

**Scheme 5:**
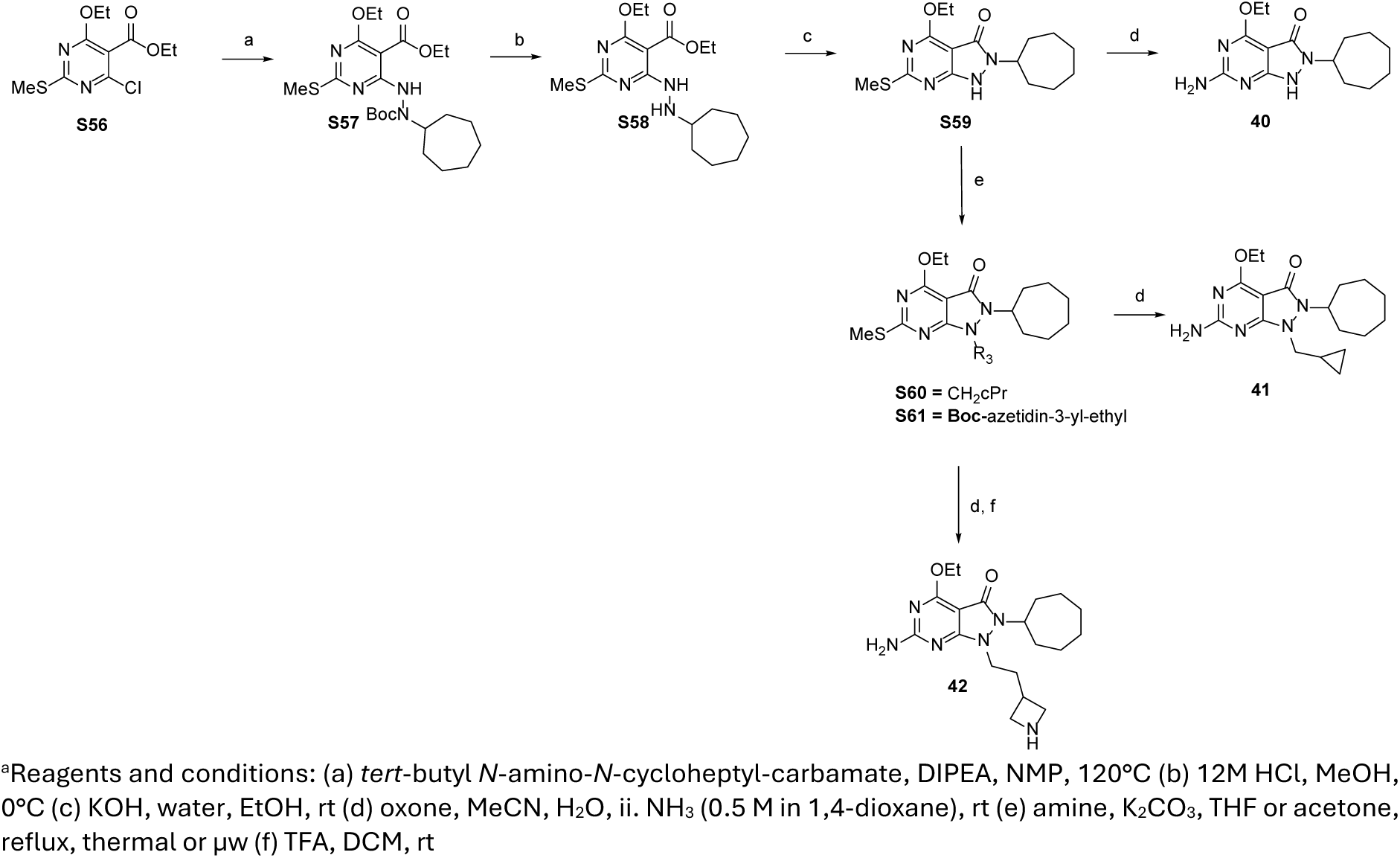
Synthesis of Compounds 40 - 42.

## CONCLUSION

Previously, we have presented a novel pre-clinical candidate (**49**) for treatment of TB through targeting *M. tuberculosis* LysRS^14^. This manuscript reports the full medicinal chemistry programme that successfully optimised a fragment-like hit into a potent nanomolar inhibitor, with minimal addition of heavy atoms. The SAR programme was guided by structure-based design as a *M. tuberculosis* LysRS crystallography platform was developed to support the design, make, test, analyse cycles. The series has been demonstrated to be active in murine *in vivo* efficacy models of TB infection, and crucially, validates a novel mode of action. Screening against panels of clinical isolates has demonstrated no apparent pre-existing resistance in the field. This work demonstrates the possibilities of evaluating a target, hypothesised to be highly vulnerable and susceptible to inhibit *M. tuberculosis* growth, and translating this approach into a drug discovery program. Inhibiting LysRS, has potential for new classes of inhibitors needed to expand the arsenal of TB drugs; to help combat a disease that profoundly impacts the lives of millions around the globe.

## EXPERIMENTAL

Synthesis of compounds **1, 10, 11, 24, 25, 29, 36**, **37, 46, 47**, **49** and **S1, S2, S20, S21, S33** and **S35** has been described previously^14, 19^.

### General Methods

Chemicals and solvents were obtained from commercial vendors and used as provided, unless specified otherwise. Dry solvents were purchased in Sure Seal bottles stored over molecular sieves. Analytical TLCs were carried out on pre-coated silica plates (Kieselgel 60 F254, BDH) with visualisation via U.V. light (UV254/365 nm) and/or potassium permanganate solution. Flash chromatography was performed using Combiflash Companion Rf (commercially available from Teledyne ISCO) and prepacked silica gel columns purchased from Teledyne ISCO. Mass-directed preparative HPLC separations were performed using a Waters HPLC (2545 binary gradient pumps, 515 HPLC make up pump, 2767 sample manager) connected to a Waters 2998 photodiode array and a Waters 3100 mass detector. Preparative HPLC separations were performed with a Gilson HPLC (321 pumps, 819 injection module, 215 liquid handler/injector) connected to a Gilson 155 UV/vis detector. On both instruments, HPLC chromatographic separations were conducted using Waters XBridge C18 columns, 19 x 100 mm, 5 um particle size; using 0.1% ammonia in water (solvent A) and acetonitrile (solvent B) or 0.1% formic acid in water (solvent A) and acetonitrile (solvent B) as mobile phase. Chiral separation was performed using Mingram Semi-prep SFC, 10 x 250 mm, 5um column. NMR spectra were recorded on a Bruker Avance DPX 500 spectrometer (1 H at 500.1 MHz), DPX 400 (1H at 400 MHz) or a 500MHz Cryo NMR (Bruker, Avance 3). Chemical shifts (δ) are expressed in ppm recorded using the residual solvent as the internal reference in all cases. Splitting patterns are defined as singlet (s), doublet (d), triplet (t), quartet (q), multiplet (m), broad (br), or a combination thereof. Coupling constants (*J*) are quoted to the nearest 0.1 Hz. High-resolution electrospray measurements were performed on a Bruker Daltonics MicroTOF mass spectrometer or on a Orbitrap Exploris 120 Mass spectrometer. Low resolution electrospray (ES) mass spectra were recorded on an Advion Compact Mass Spectrometer (CMS; model ExpressIon CMS) connected to Dionex Ultimate 3000 UPLC system with diode array detector. UPLC chromatographic separations were conducted using a Waters XBridge C18 column, 2.1 x 50mm, 3.5 μm particle size or Waters XSelect 2.1 x 30mm, 2.5 μm particle size. The compounds were eluted with a gradient of 5 to 95% acetonitrile/water +0.1% Ammonia or +0.1% formic acid. Unless otherwise stated, all compounds were >95% pure by HPLC. Microwave-assisted chemistry was performed using a CEM or Biotage microwave synthesizer.

### 2-Amino-6-cyclohexyl-6,7-dihydro-5*H*-pyrrolo[3,4-d]pyrimidin-5-one (2)

To a suspension of cyclohexanamine (58 mg, 0.58 mmol) and Cs_2_CO_3_ (113 mg, 0.87 mmol) in MeCN (3 mL) was added ethyl 2-amino-4-(bromomethyl)pyrimidine-5-carboxylate hydrobromide **(S1)** (100 mg, 0.29 mmol) dropwise as a solution in NMP (1 mL). The reaction was stirred for 30 min at rt then heated under µW conditions at 130 °C for 90 min, filtered and purified using prep-HPLC to afford compound **2** (20 mg, 28%) as a white solid. ^1^H NMR (400 MHz, CD_3_OD) *δ* 8.55 (d, *J* = 1.3 Hz, 1H), 4.31 (s, 2H), 4.08 (tt, *J* = 11.8, 3.9 Hz, 1H), 1.94 – 1.76 (m, 4H), 1.71 (d, *J* = 13.3 Hz, 1H), 1.62 – 1.36 (m, 4H), 1.31 – 1.12 (m, 1H). HRMS (ESI) calcd. for [M + H]^+^ C_12_H_17_ON_4_ 233.1402 found 233.1399

### 2-Amino-6-cycloheptyl-6,7-dihydro-5*H*-pyrrolo[3,4-d]pyrimidin-5-one (3)

Reaction of cycloheptanamine (79 mg, 0.70 mmol) and ethyl 2-amino-4-(bromomethyl)pyrimidine-5-carboxylate hydrobromide (**S1)** (120 mg, 0.35 mmol), Cs_2_CO_3_ (136 mg, 1.06 mmol), MeCN (3 mL) and NMP (1 mL) by an analogous method to compound **2** afforded **3** a white solid (23 mg, 25%). ^1^H NMR (400 MHz, CD_3_OD) *δ* 8.54 (s, 1H), 4.36 – 4.23 (m, 3H), 1.93 – 1.54 (m, 12H). HRMS (ESI) calcd. for [M + H]^+^ C_13_H_19_ON_4_ 247.1558 found 247.1556

### 2-Amino-6-((1*S*,2*R*)-2-hydroxycyclohexyl)-6,7-dihydro-5*H*-pyrrolo[3,4-*d*]pyrimidin-5-one (4)

To a suspension of ethyl 2-amino-4-(bromomethyl)pyrimidine-5-carboxylate (S1) (100 mg, 0.38 mmol) and (1*R*,2*S*)-2-aminocyclohexanol hydrochloride (116 mg, 0.77 mmol) in NMP (0.64 mL) was added TEA (0.41 mL, 2.31 mmol). The reaction was stirred at rt for 1 h then heated under µW conditions at 130 °C for 2 h and purified by prep-HPLC to afford compound 4 (18 mg, 18%), as an off white solid. ^1^H NMR (400 MHz, DMSO-*d_6_*) *δ* 8.51 (s, 1H), 7.32 (s, 2H), 4.75 (d, *J* = 4.4 Hz, 1H), 4.53 (d, *J* = 18.7 Hz, 1H), 4.30 (d, *J* = 18.7 Hz, 1H), 4.02 – 3.87 (m, 2H), 2.01 – 1.90 (m, 1H), 1.73 (dd, *J* = 27.9, 12.5 Hz, 2H), 1.61 – 1.44 (m, 3H), 1.42 – 1.29 (m, 2H). HRMS (ESI) calcd. for [M + H]^+^ C_12_H_17_O_2_N_4_ 249.1351 found 249.1344

### 2-Amino-6-((1*S*,2*S*)-2-hydroxycyclohexyl)-6,7-dihydro-5*H*-pyrrolo[3,4-*d*]pyrimidin-5-one (5)

Reaction of ethyl 2-amino-4-(bromomethyl)pyrimidine-5-carboxylate (S1) (100 mg, 0.38 mmol), (1*S*,2*S*)-2-aminocyclohexanol hydrochloride (116 mg, 0.76 mmol), NMP (0.7 mL) and TEA (0.41 mL, 2.31 mmol) in an analogous method to compound 4 to afforded 5 as an off white solid (32 mg, 30%). ^1^H NMR (400 MHz, DMSO*-d_6_*) *δ* 8.49 (s, 1H), 7.32 (s, 2H), 4.74 (d, *J* = 5.2 Hz, 1H), 4.35 – 4.16 (m, 2H), 3.83 – 3.70 (m, 1H), 3.59 – 3.43 (m, 1H), 1.99 – 1.89 (m, 1H), 1.72 – 1.58 (m, 3H), 1.57 – 1.44 (m, 1H), 1.34 – 1.17 (m, 3H). HRMS (ESI) calcd. for [M + H]^+^ C_12_H_17_O_2_N_4_ 249.1351 found 249.1345

### 2-Amino-4-chloro-6-cyclohexyl-6,7-dihydro-5*H*-pyrrolo[3,4-d]pyrimidin-5-one (6)

To a solution of 4-chloro-6-cyclohexyl-2-(methylthio)-6,7-dihydro-5*H*-pyrrolo[3,4-*d*]pyrimidin-5-one **(S41)** (130 mg, 0.44 mmol) in MeCN (3 mL) and water (0.5 mL) was added oxone (322 mg, 0.52 mmol). The reaction was stirred at rt for 16 h, partitioned between DCM and water, passed through a hydrophobic frit and concentrated *in vacuo*. The crude material was dissolved in NH_3_ (0.5 M in 1,4-dioxane, 9 mL) the reaction sealed and stirred at rt for 16 h, concentrated *in vacuo* and purified by prep-HPLC to afford compound **6** (35 mg, 28%) as a white solid. ^1^H NMR (400 MHz, DMSO*-d_6_*) *δ* 7.70 (bs, 2H), 4.26 (s, 2H), 3.93 (tt, 1H, *J* = 11.9, 3.5 Hz), 1.81 - 1.60 (m, 5H), 1.48 (qd, 2H, *J* = 12.4, 3.2 Hz), 1.40 - 1.27 (m, 2H), 1.19 - 1.07 (m, 1H). HRMS (ESI) calcd. for [M + H]^+^ C_12_H_16_ON_4_Cl 267.1013 found 267.1008

### 2-Amino-4-bromo-6-cycloheptyl-7H-pyrrolo[3,4-d]pyrimidin-5-one (7)

To a solution of ethyl 4-chloro-6-formyl-2-(methylthio)pyrimidine-5-carboxylate **(S14)** (2000 mg, 7.67 mmol) in THF (110 mL) was added aminocycloheptane (955 mg, 8.43 mmol), sodium triacetoxyborohydride (4877 mg, 23.01 mmol) and MgSO_4_. The reaction stirred at rt for 16 h, concentrated *in vacuo* and the residue partitioned between EtOAc and water. The combined organic layers were dried over Na_2_SO_4_, concentrated *in vacuo* and purified by column chromatography (0 – 50 % EtOAc in heptane) to afford 4-chloro-6-cycloheptyl-2-(methylthio)-6,7-dihydro-5*H*-pyrrolo[3,4-*d*]pyrimidin-5-one **(S45)**. MS (ES+) *m/z* 312.2 [M+ H]^+^. To a solution of compound **S45** (100 mg, 0.32 mmol) in MeCN (5 mL) was added TMSBr (0.55 mL, 4.17 mmol) and the reaction was stirred at 40 °C for 3 h. A second portion of TMSBr (0.550 mL, 4.17 mmol) was added and the reaction heated at 40 °C for a further 48 h, the reaction mixture partitioned between DCM and water and the combined organic layers dried over Na_2_SO_4_, concentrated *in vacuo* and purified by flash chromatography (0 - 60% EtOAc in heptane) to afford 4-bromo-6-cycloheptyl-2-(methylthio)-6,7-dihydro-5*H*-pyrrolo[3,4-*d*]pyrimidin-5-one **(S52)**. MS (ES+) *m/z* 356.3 [M + H]^+^. Reaction of **S52** (74 mg, 0.21 mmol), oxone (153 mg, 0.25 mmol), NH_3_ (0.5 M in 1,4-dioxane, 4 mL), water (0.5 mL) and MeCN (3 mL) in an analogous method to compound **6** afforded **7** as a tan solid (10 mg, 5% over three steps). ^1^H NMR (400 MHz, DMSO*-d_6_*) *δ* 7.74 (br s, 2H), 4.26 (s, 2H), 4.15 – 4.11 (m, 1H), 1.74 – 1.45 (m, 12H). HRMS (ESI) calcd. for [M + H]^+^ C_13_H_18_BrN_4_O, 325.0663 found 325.0688

### 2-Amino-6-cyclohexyl-4-methoxy-6,7-dihydro-5*H*-pyrrolo[3,4-d]pyrimidin-5-one (8)

4-Chloro-6-cyclohexyl-2-(methylthio)-6,7-dihydro-5*H*-pyrrolo[3,4-*d*]pyrimidin-5-one **(S44)** was synthesised from ethyl 4-chloro-6-formyl-2-(methylthio)pyrimidine-5-carboxylate **(S14)** (200 mg, 0.76 mmol), cyclohexanamine (84 mg, 0.84 mmol), sodium triacetoxyborohydride (163 mg, 0.76 mmol) and THF (10 mL) in an analogous method to compound **S45**. The residue of compound **S44** was dissolved in MeOH (2 mL), the reaction sealed and heated under µW conditions at 120 ⁰C for 30 min and concentrated *in vacuo* to afford 6-cyclohexyl-4-methoxy-2-(methylthio)-6,7-dihydro-5*H*-pyrrolo[3,4-*d*]pyrimidin-5-one (**S46)**. Reaction of **S46** and NH_3_ (0.5 M in 1,4-dioxane, 4mL) in an analogous method to compound **6** at 80 °C for 4 h afforded **8** as a white solid (50 mg, 22% yield over 3 steps). ^1^H NMR (500 MHz, DMSO*-d_6_*) *δ* 7.19 (s, 2H), 4.14 (s, 2H), 3.89 – 3.84 (m, 4H), 1.77 – 1.75 (m, 2H), 1.66 – 1.61 (m, 3H), 1.44 (qd, *J* = 12.4, 3.4 Hz, 2H), 1.33 (qt, *J* = 13.2, 3.4 Hz, 2H), 1.11 (dddd, *J* = 16.2, 12.6, 8.2, 3.6 Hz, 1H). HRMS (ESI) calcd. for [M + H]^+^ C_13_H_19_O_2_N_4_ 263.1508 found 263.1505

### 2-Amino-6-cycloheptyl-4-methoxy-7H-pyrrolo[3,4-d]pyrimidin-5-one (9)

Reaction of 4-chloro-6-cycloheptyl-2-(methylthio)-6,7-dihydro-5*H*-pyrrolo[3,4-*d*]pyrimidin-5-one **(S47)** (72 mg, 0.23 mmol), oxone (144 mg, 0.23 mmol), MeCN (8 mL), water (4 mL) and NH_3_ (0.5 M in 1,4-dioxane, 4mL) in an analogous method to compound **6** at 90 °C for 2 h afforded **9** as a white solid (36 mg, 53%). ¹H NMR (400 MHz, CDCl_3_) δ 5.26 (2H, s), 4.37 - 4.32 (1H, m), 4.12 (2H, s), 4.04 (3H, s), 1.90 - 1.85 (2H, m), 1.74 - 1.52 (10H, m). HRMS (ESI) calcd. for [M + H]^+^ C_14_H_21_N_4_O_2_ 277.1664, found 277.1671

### 2-Amino-6-cycloheptyl-4-propoxy-6,7-dihydro-5*H*-pyrrolo[3,4-*d*]pyrimidin-5-one (12)

Reaction of 6-cycloheptyl-2-(methylthio)-4-propoxy-6,7-dihydro-5*H*-pyrrolo[3,4-*d*]pyrimidin-5-one **(S22)** (74 mg, 0.23 mmol), oxone (283 mg, 0.46 mmol), MeCN (5mL), water (2.5 mL) and NH_3_ (0.5 M in 1,4-dioxane, 2 mL) in an analogous method to compound **6** at 105 °C for 16 h and purification by prep-HPLC to afforded **12** as a white solid (68 mg, 95%). ¹H NMR (500 MHz, DMSO*-d_6_*) *δ* 7.14 (s, 2H), 4.28 (t, *J* = 6.8 Hz, 2H), 4.14 (s, 2H), 4.09 - 4.04 (m, 1H), 1.74 - 1.42 (m, 14H), 0.96 (t, *J* = 7.4 Hz, 3H). HRMS (ESI) calcd. for [M + H]^+^ C_16_H_25_N_4_O_2_ 305.1277, found 105.1998

### 2-Amino-6-cyclohexyl-4-(2-hydroxyethoxy)-6,7-dihydro-5*H*-pyrrolo[3,4-*d*]pyrimidin-5-one (13)

To a solution of 2-amino-6-cyclohexyl-4-(2-methoxyethoxy)-6,7-dihydro-5*H*-pyrrolo[3,4-*d*]pyrimidin-5-one **(14)** (123 mg, 0.40 mmol) in DCM (5 mL) at -78 °C under N_2_ was added tribromoborane (1M in DCM, 1.20 mL, 1.20 mmol) dropwise. The reaction stirred at rt for 3.5 h, quenched with dropwise addition of water at 0 °C and partitioned between DCM and water, passed through a hydrophobic frit, concentrated *in vacuo* and purified by prep-HPLC to afford compound **13** (6 mg, 5%) as a white solid. ^1^H NMR (500 MHz, DMSO*-d_6_*) *δ* 7.15 (s, 2H), 4.80 (t, *J* = 5.4 Hz, 1H), 4.37 (t, *J* = 5.3 Hz, 2H), 4.14 (s, 2H), 3 - 87 (tt, *J* = 11.9, 3.8 Hz, 1H), 3.70 (q, *J* = 5.3 Hz, 2H 1.76 (d, J = 13.0 Hz, 2H), 1.64 (t, *J* = 13.1 Hz, 3H), 1.45 (qd, *J* = 12.3, 3.0 Hz, 2H), 1.33 (q, *J* = 12.9 Hz, 2H), 1.12 (dd, *J* = 17.6, 8.2 Hz, 1H). HRMS (ESI) calcd. for [M + H]^+^ C_14_H_21_N_4_O_3_ 293.1613, found 293.1617

### 2-Amino-6-cyclohexyl-4-(2-methoxyethoxy)-6,7-dihydro-5*H*-pyrrolo[3,4-*d*]pyrimidin-5-one (14)

To a solution of 6-cyclohexyl-4-(2-methoxyethoxy)-2-(methylthio)-6,7-dihydro-5*H*-pyrrolo[3,4-*d*]pyrimidin-5-one **(S23)** (53 mg, 0.16 mmol) in DCM (4 mL) at 0 °C was added *m*-CPBA (54 mg, 0.24 mmol). The reaction stirred at rt for 1.5 h, concentrated *in vacuo* and the crude material suspended in NH_3_ (0.5 M in 1,4-dioxane, 1.25 mL), the reaction sealed and stirred at 105 °C for 16 h, concentrated *in vacuo* and purified by prep-HPLC to afford compound **14** (15 mg, 29 %) as an off-white solid. ^1^H NMR (400 MHz, DMSO*-d_6_*) *δ* 7.21 (s, 2H), 4.52 – 4.37 (m, 2H), 4.15 (s, 2H), 3.86 (ddt, *J* = 11.7, 7.6, 3.8 Hz, 1H), 3.69 – 3.60 (m, 2H), 3.30 (s, 3H), 1.76 (d, *J* = 12.7 Hz, 2H), 1.66 – 1.60 (m, 3H), 1.45 (qd, *J* = 12.1, 3.1 Hz, 2H), 1.30 (dt, *J* = 27.2, 13.4 Hz, 2H), 1.16 – 1.06 (m, 1H). HRMS (ESI) calcd. for [M + H]^+^ C_15_H_23_N_4_O_3_ 307.1770, found 307.1771

### 2-Amino-6-cyclohexyl-4-(2,2-difluoropropoxy)-6,7-dihydro-5*H*-pyrrolo[3,4-*d*]pyrimidin-5-one (15)

To a solution of 4-chloro-6-cyclohexyl-2-(methylthio)-6,7-dihydro-5*H*-pyrrolo[3,4-*d*]pyrimidin-5-one **(S44)** (68 mg, 0.23 mmol) in THF (2 mL) was added Cs_2_CO_3_ (223 mg, 0.68 mmol) and 2,2-difluoropropan-1-ol (0.05 mL, 0.57 mmol). The reaction stirred at 80 °C for 40 mins, diluted with EtOAc and brine, the combined organic layers dried over MgSO_4_ and concentrated *in vacuo* to afford 6-cyclohexyl-4-(2,2-difluoropropoxy)-2-(methylthio)-6,7-dihydro-5*H*-pyrrolo[3,4-*d*]pyrimidin-5-one **(S48)** (81 mg, 0.27 mmol). Reaction of **S48** (81 mg, 0.27 mmol), *m*-CPBA (73 mg, 0.34 mmol), DCM (4 mL) and NH_3_ (0.5 M in 1,4-dioxane, 1.8 mL) in an analogous manner to compound **14** afforded **15** as a white solid (10 mg, 13% over two steps). ^1^H NMR (400 MHz, DMSO) *δ* 7.34 (s, 8H), 4.63 (t, *J* = 12.7 Hz, 8H), 4.18 (s, 8H), 3.93 – 3.79 (m, 4H), 1.81 – 1.57 (m, 34H), 1.46 (dt, *J* = 12.0, 9.2 Hz, 9H), 1.30 (dt, *J* = 26.9, 13.5 Hz, 10H), 1.12 (t, *J* = 13.0 Hz, 5H). HRMS (ESI) calcd. for [M + H]^+^ C_15_H_21_F_2_N_4_O_2_ 327.1633, found 327.1645

### 2-Amino-6-cyclohexyl-4-(difluoromethoxy)-6,7-dihydro-5*H*-pyrrolo[3,4-d]pyrimidin-5-one (16)

Reaction of 6-cyclohexyl-4-(difluoromethoxy)-2-(methylthio)-6,7-dihydro-5*H*-pyrrolo[3,4-*d*]pyrimidin-5-one **(S25)** (90 mg, 0.27 mmol), oxone (252 mg, 0.41 mmol), MeCN (3 mL), water (0.5 mL) and NH_3_ (0.5 M in 1,4-dioxane, 0.5 mL) in an analogous manner to compound **6,** heating the displacement at 100 °C for 2 h afforded **16** as a white solid (30 mg, 36%). ^1^H NMR (500 MHz, DMSO*-d_6_*) *δ* 7.73 (t, 1H, *J* = 71.6 Hz), 7.57 (bs, 2H), 4.26 (s, 2H), 3.91 (tt, 1H, *J* = 11.6, 3.6 Hz), 1.79 - 1.62 (m, 5H), 1.48 (qd, 2H, *J* = 12.4, 3.2 Hz), 1.39 - 1.29 (m, 2H), 1.14 - 1.11 (m, 1H). HRMS (ESI) calcd. for [M + H]^+^ C_13_H_17_F_2_N_4_O_2_ 299.1319, found 299.1326

### 2-Amino-6-cyclohexyl-4-(2,2-difluoroethoxy)-6,7-dihydro-5*H*-pyrrolo[3,4-*d*]pyrimidin-5-one (17)

Reaction of 6-cyclohexyl-4-(2,2-difluoroethoxy)-2-(methylthio)-6,7-dihydro-5*H*-pyrrolo[3,4-*d*]pyrimidin-5-one **(S24)** (134 mg, 0.39 mmol), *m*-CPBA (135 mg, 0.59 mmol), DCM (4 mL) and NH_3_ (0.5 M in 1,4-dioxane, 3 mL) in an analogous manner to compound **14** afforded **17** as a white solid (39 mg, 30%). ^1^H NMR (400 MHz, DMSO*-d_6_*) *δ* 7.36 (s, 2H), 6.41 (tt, *J* = 54.9, 3.8 Hz, 1H), 4.65 (td, *J* = 14.6, 3.7 Hz, 2H), 4.19 (s, 2H), 3.95 – 3.80 (m, 1H), 1.80 – 1.76 (m, 2H), 1.67 – 1.65 (m, 3H), 1.51 – 1.32 (m, 4H), 1.13 – 1. (m, 1H). HRMS (ESI) calcd. for [M + H]^+^ C_14_H_19_F_2_N_4_O_2_ 313.1476 found 313.1485

### 2-Amino-6-cyclohexyl-4-(methylamino)-6,7-dihydro-5*H*-pyrrolo[3,4-d]pyrimidin-5-one (18)

To a solution of 4-chloro-6-cyclohexyl-2-(methylthio)-6,7-dihydro-5*H*-pyrrolo[3,4-*d*]pyrimidin-5-one **(S44)** (50 mg, 0.17 mmol) in THF (1 mL) was added methanamine (2 M in THF, 0.840 mL). The rection was stirred at rt for 1 h and concentrated *in vacuo* to afford 6-cyclohexyl-4-(methylamino)-2-(methylthio)-6,7-dihydro-5*H*-pyrrolo[3,4-*d*]pyrimidin-5-one **(S49)** (50 mg, 0.16 mmol). MS (ES+) m/z 293.1 [M + H]^+^ Reaction of **S49** (50 mg, 0.16 mmol), *m*-CPBA (89 mg, 0.43 mmol) in DCM (10 mL) and NH_3_ (0.5 M in 1,4-dioxane, 1.25 mL) in an analogous manner to compound **14** afforded **18** as a white solid (14 mg, 47% over two steps). ^1^H NMR (400 MHz, DMSO-*d_6_*) *δ* 8.24 (s, 1H), 6.80 - 6.58 (m, 3H), 4.07 (s, 2H), 3.91 - 3.78 (t, *J* = 11.6 Hz, 1H), 2.92-2.83 (m, 3H), 1.82 - 1.72 (m, 2H), 1.71 - 1.58 (m, 3H), 1.52 -1.40 (m, 2H), 1.38 -1.27 (m, 2H), 1.18 -1.05 (m, 1H). HRMS (ESI) calcd. for [M + H]^+^ C_13_H_20_ON_5_ 262.1668 found 262.1664

### 2-Amino-6-cyclohexyl-4-(ethylamino)-6,7-dihydro-5*H*-pyrrolo[3,4-d]pyrimidin-5-one (19)

Reaction of 6-cyclohexyl-4-(ethylamino)-2-(methylthio)-6,7-dihydro-5*H*-pyrrolo[3,4-*d*]pyrimidin-5-one **(S50)** (100 mg, 0.33 mmol), oxone (300 mg, 0.49 mmol), MeCN (4 mL), water (0.25 mL) and NH_3_ (0.5 M in 1,4-dioxane, 1 mL) in an analogous manner to compound **6,** heating the displacement at 100 °C, afforded **19** as a white solid (30 mg, 30%). ^1^H NMR (400 MHz, DMSO) δ 6.83 – 6.45 (m, 3H), 4.06 (s, 2H), 3.89 – 3.76 (m, 1H), 3.41 (pent, *J* = 6.1 Hz, 2H), 1.76 (d, *J* = 12.7 Hz, 2H), 1.70 – 1.57 (m, *J* = 12.7 Hz, 3H), 1.45 (dt, *J* = 12.5, 11.1 Hz, 2H), 1.31 (dd, *J* = 25.8, 12.7 Hz, 2H), 1.18 – 1.03 (m, *J* = 14.3, 7.1 Hz, 4H). HRMS (ESI) calcd. for [M + H]^+^ C_14_H_22_N_5_O 276.1824 found 276.1835

### 2-Amino-6-cyclohexyl-4-(dimethylamino)-6,7-dihydro-5*H*-pyrrolo[3,4-d]pyrimidin-5-one (20)

To a solution of 4-chloro-6-cyclohexyl-2-(methylthio)-6,7-dihydro-5*H*-pyrrolo[3,4-*d*]pyrimidin-5-one **(S44)** (100 mg, 0.35 mmol) in THF (2 mL) was added dimethyl amine hydrochloride (42 mg, 0.52 mmol) and DIPEA (0.18 mL, 1.04 mmol). The reaction was stirred at 80 °C for 80 min, diluted with DCM and water, passed through a hydrophobic frit and concentrated *in vacuo* to afford 4-chloro-6-cyclohexyl-2-(methylthio)-6,7-dihydro-5*H*-pyrrolo[3,4-*d*]pyrimidin-5-one **(S51)** (109 mg, 0.36 mmol). MS (ES+) m/z 307.4 [M + H]^+^ Reaction of **S51** (109 mg, 0.36 mmol), oxone (437 mg, 0.71 mmol), MeCN (2 mL), water (1 mL) and NH_3_ (0.5 M in 1,4-dioxane, 3 mL) in an analogous manner to compound **6,** heating the displacement at 105 °C, afforded **20** as a white solid (40 mg, 39% over two steps). ^1^H NMR (500 MHz, DMSO*-d_6_*) *δ* 6.56 (s, 2H), 4.04 (s, 2H), 3.88 (tt, *J* = 11.9, 3.7 Hz, 1H), 3.26 (s, 6H), 1.78 – 1.75 (m, 2H), 1.66 – 1.61 (m, 3H), 1.43 (qd, *J* = 12.3, 3.2 Hz, 2H), 1.37 – 1.26 (m, 2H), 1.11 (qt, *J* = 12.8, 3.6 Hz, 1H). HRMS (ESI) calcd. for [M + H]^+^ C_14_H_22_ON_5_ 276.1824 found 276.1823

### 2-Amino-6-cyclohexyl-5-oxo-6,7-dihydro-5*H*-pyrrolo[3,4-d]pyrimidine-4-carbonitrile (21)

Reaction of 6-cyclohexyl-2-(methylthio)-5-oxo-6,7-dihydro-5*H*-pyrrolo[3,4-*d*]pyrimidine-4-carbonitrile **(S55)** (29 mg, 3.46 mmol), oxone (160 mg, 0.26 mmol), MeCN (2 mL), water (0.2mL) and NH_3_ (0.5 M in 1,4-dioxane, 3.5 mL) in an analogous manner to compound **6** heating the displacement at 100 °C for 2 h, afforded **21** as a white solid (16 mg, 34%). ^1^H NMR (400 MHz, DMSO*-d_6_*) *δ* 7.88 (bs, 2H), 4.35 (s, 2H), 3.95 (tt, 1H, *J* = 11.8, 3.8 Hz), 1.86 - 1.58 (m, 5H), 1.51 (qd, 2H, *J* = 12.3, 3.1 Hz), 1.39 - 1.30 (m, 1H), 1.17 – 1.11 (m, 1H). HRMS (ESI) calcd. for [M + H]^+^ C_13_H_16_N_5_O 258.1354 found 258.1373

### 2-Amino-6-cycloheptyl-4-(methoxymethyl)-6,7-dihydro-5*H*-pyrrolo[3,4-d]pyrimidin-5-one (22)

Reaction of 6-cycloheptyl-4-(methoxymethyl)-2-(methylthio)-6,7-dihydro-5*H*-pyrrolo[3,4-*d*]pyrimidin-5-one **(S53)** (16 mg, 0.05 mmol), oxone (40 mg, 0.06 mmol), MeCN (2 mL), water (1 mL) and NH_3_ (0.5 M in 1,4-dioxane, 1 mL) in an analogous manner to compound **6**, running the oxidation step for 90 min, afforded **22** as a white solid (5 mg, 33%). ^1^H NMR (400 MHz, DMSO*-d_6_*) *δ* 7.35 (s, 2H), 4.63 (s, 2H), 4.24 (s, 2H), 4.14 – 4.09 (m, 1H), 3.32 (s, 3H), 1.74 – 1.44 (m, 12H). HRMS (ESI) calcd. for [M + H]+ C_15_H_23_N_4_O_2_ 291.1821 found 291.1816

### 2-Amino-4-ethoxy-6-phenyl-6,7-dihydro-5*H*-pyrrolo[3,4-d]pyrimidin-5-one (23)

To a solution of ethyl 4-ethoxy-6-formyl-2-(methylthio)pyrimidine-5-carboxylate **(S20)** (100 mg, 0.37 mmol) in THF (3.7 mL) was added aniline (0.04mL, 0.37 mmol), MgSO_4_ and the reaction left for 1 h before the addition of sodium triacetoxyborohydride (235 mg, 1.1 mmol) and a drop of AcOH. The reaction was stirred at rt for 16 h, diluted with DCM and water, passed through a hydrophobic frit and concentrated *in vacuo*. The residue was dissolved in EtOH (3.7 mL) and Water (1 mL) and NaOH (62 mg, 1.48 mmol) added and stirred at rt for 1 h, diluted with water and DCM, passed through a hydrophobic frit, loaded onto a SCX/PEAK cartridge, eluted with DCM and concentrated *in vacuo* to afford 4-ethoxy-2-(methylthio)-6-phenyl-6,7-dihydro-5*H*-pyrrolo[3,4-*d*]pyrimidin-5-one **(S26)**. MS (ES+) m/z 302.3 [M + H]^+^ Reaction of **S26** (111 mg, 0.37 mmol), oxone (454 mg, 0.74 mmol), MeCN (6 mL), H_2_O (3 mL) and NH_3_ (3 M in 1,4-dioxane, 1 mL) in an analogous manner to compound **6**, heating the displacement at 105°C, afforded **23** as a white solid (10 mg, 10% over two steps). ^1^H NMR (400 MHz, DMSO*-d_6_*) *δ* 7.83 – 7.69 (m, 2H), 7.48 – 7.26 (m, 4H), 7.13 – 7.01 (m, 1H), 4.72 (s, 2H), 4.44 (q, *J* = 7.1 Hz, 2H), 1.34 (t, *J* = 7.1 Hz, 3H). HRMS (ESI) calcd. for [M + H]^+^ C_14_H_15_O_2_N_4_ 271.1195 found 271.1194

### 2-Amino-6-((1R,3S)-3-hydroxycyclohexyl)-4-methoxy-6,7-dihydro-5*H*-pyrrolo[3,4-*d*]pyrimidin-5-one (26)

Reaction of ethyl 2-((*tert*-butoxycarbonyl)amino)-4-formyl-6-methoxypyrimidine-5-carboxylate **(S37)** (200 mg, 0.64 mmol), (1*S*,3*R*)-3-aminocyclohexan-1-ol hydrochloride (97 mg, 0.64 mmol), sodium triacetoxyborohydride (408 mg, 1.92 mmol) and DCM (4 mL) following an analogous method to compound **S45**, with the addition of DIPEA (0.11 mL, 0.64 mmol) afforded *tert*-butyl (6-((1*R*,3*S*)-3-hydroxycyclohexyl)-4-methoxy-5-oxo-6,7-dihydro-5*H*-pyrrolo[3,4-*d*]pyrimidin-2-yl)carbamate **(S38)** MS (ES+) m/z 379.5 [M + H]^+^ To a solution of compound **S38** (147 mg, 0.39mmol) in DCM (3 mL) was added TFA (0.12 mL, 1.55 mmol). The reaction was stirred at rt for 16 h, concentrated *in vacuo* and the residue dissolved in MeCN (4 mL) and 2M NaOH (1 mL) stirred at rt for 4 h, concentrated *in vacuo* and purified by prep-HPLC to afford compound **26** (50 mg, 27% over two steps) as a white solid. ¹H NMR (400 MHz, DMSO*-d_6_*) *δ* 7.21 (2H, s), 4.67 (1H, s), 4.15 (2H, s), 3.95 – 3.90 (4H, m), 3.51 - 3.49 (1H, m), 1.87 - 1.70 (3H, m), 1.59 - 1.55 (1H, m), 1.42 - 1.27 (3H, m), 1.10 - 1.01 (1H, m) HRMS (ESI) calcd. for [M + H]^+^ C_13_H_19_N_4_O_3_ 279.1457 found 279.1447

### 2-Amino-6-(cis-4-hydroxycyclohexyl)-4-methoxy-6,7-dihydro-5*H*-pyrrolo[3,4-*d*]pyrimidin-5-one (27)

Reaction of ethyl 2-((*tert*-butoxycarbonyl)amino)-4-formyl-6-methoxypyrimidine-5-carboxylate **(S37)** (250 mg, 0.80 mmol), *cis*-4-amino-cyclohexanol (93 mg, 0.80 mmol), sodium triacetoxyborohydride (511 mg, 3.41 mmol) in DCM (4 mL) in an analogous manner to compound **S45** afforded *tert*-butyl (6-((1*S*,4*S*)-4-hydroxycyclohexyl)-4-methoxy-5-oxo-6,7-dihydro-5*H*-pyrrolo[3,4-*d*]pyrimidin-2-yl)carbamate (**S39)**. MS (ES+) m/z 379.4 [M + H]^+^ To a solution of **S39** (50 mg, 0.13 mmol) in DCM (2 mL) was added HCl (4 M in 1,4-dioxane, 0.17 mL, 0.66 mmol). The reaction was stirred at rt for 16 h concentrated *in vacuo*, partitioned between a 3:1 solution of DCM:IPA and sat. NaHCO_3_, the combined organic layers were concentrated *in vacuo* and purified by prep-HPLC to afford **27** (11 mg, 13% over two steps) as a white solid. ^1^H NMR (400 MHz, DMSO*-d_6_*) *δ* 7.24 (s, 2H), 4.38 (s, 1H), 4.15 (s, 2H), 3.95 – 3.84 (m, 4H), 3.82 (s, 1H), 1.95 – 1.77 (m, 2H), 1.72 (d, *J* = 13.5 Hz, 2H), 1.50 (t, *J* = 13.5 Hz, 2H), 1.36 (d, *J* = 11.9 Hz, 2H). HRMS (ESI) calcd. for [M + H]^+^ C_13_H_19_N_4_O_3_ 279.1457 found 279.1481

### 2-Amino-6-(cis-4-(hydroxymethyl)cyclohexyl)-4-methoxy-6,7-dihydro-5*H*-pyrrolo[3,4-*d*]pyrimidin-5-one (28)

Reaction of ethyl 2-((*tert*-butoxycarbonyl)amino)-4-formyl-6-methoxypyrimidine-5-carboxylate **(S37)** (100 mg, 0.32 mmol), *cis*-(4-aminocyclohexyl)methanol (41.5 mg, 0.32 mmol) and sodium triacetoxyborohydride (204 mg, 0.96 mmol) in DCM (5 mL) following an analogous method to compound **S45** afforded *tert*-butyl (6-((1s,4s)-4-(hydroxymethyl)cyclohexyl)-4-methoxy-5-oxo-6,7-dihydro-5*H*-pyrrolo[3,4-*d*]pyrimidin-2-yl)carbamate **(S40)**. Reaction of **S40,** TFA (1 mL), MeCN (3 mL) and NaOH (2M, 2 mL) by an analogous method to compound **26** afforded **28** as a white solid (23 mg, 22% over two steps). ^1^H NMR (400 MHz, CDCl_3_) *δ* 5.31 (s, 2H), 4.21 – 4.07 (m, 3H), 4.01 (s, 3H), 3.67 (d, *J* = 7.4 Hz, 2H), 1.92 – 1.76 (m, 5H), 1.74 – 1.53 (m, 7H). HRMS (ESI) calcd. for [M + H]^+^ C_14_H_21_N_4_O_3_ 293.1613 found 293.1610

### 2-Amino-6-(2,2-difluorocyclohexyl)-4-methoxy-6,7-dihydro-5*H*-pyrrolo[3,4-*d*]pyrimidin-5-one (29)

Reaction of ethyl 4-ethoxy-6-formyl-2-(methylthio)pyrimidine-5-carboxylate **(S16)** (100 mg, 0.37 mmol), 2,2-difluorocyclohexanamine (60 mg, 0.44 mmol), sodium triacetoxyborohydride (235mg, 1.11 mmol) in (10.5 mL) in an analogous manner to compound **S45** afforded 6-(2,2-difluorocyclohexyl)-4-methoxy-2-(methylthio)-6,7-dihydro-5*H*-pyrrolo[3,4-*d*]pyrimidin-5-one **(S27).** Reaction of **S27**, oxone (341 mg, 0.55 mmol), MeCN (2 mL), water (0.5 mL) and NH_3_ (3 M in 1,4-dioxane, 2 mL), in an analogous manner to compound **6**, stirring the displacement at 90 °C, afforded **29** as a white solid (39 mg, 32% over two steps). ^1^H NMR (400 MHz, DMSO-d_6_) *δ* 7.26 (s, 2H), 4.49 – 4.29 (m, 4H), 4.17 (d, *J* = 16.9 Hz, 1H), 2.17 – 2.02 (m, 1H), 2.02 – 1.67 (m, 5H), 1.58 – 1.36 (m, *J* = 31.1, 12.9 Hz, 2H), 1.32 (t, *J* = 7.1 Hz, 3H). HRMS (ESI) calcd. for [M + H]^+^ C_14_H_19_F_2_N_4_O_2_ 313.1476 found 313.1506

### 2-Amino-6-(3,3-difluorocyclohexyl)-4-methoxy-6,7-dihydro-5*H*-pyrrolo[3,4-*d*]pyrimidin-5-one (30)

Reaction of 6-(3,3-difluorocyclohexyl)-4-ethoxy-2-(methylthio)-6,7-dihydro-5*H*-pyrrolo[3,4-*d*]pyrimidin-5-one **(S28)** (860 mg, 2.5 mmol), oxone (2309 mg, 3.76 mmol), MeCN (30 mL), water (2 mL) and NH_3_ (3 M in 1,4-dioxane, 50 mL) in an analogous manner to compound **6**, stirring the oxidation for 2 h followed by heating the displacement at 100 °C for 2 h afforded **30** as a white solid (480 mg, 60%). ^1^H NMR (500 MHz, DMSO*-d_6_*) *δ* 7.19 (bs, 2H), 4.41 (q, 2H, *J* = 7.2 Hz), 4.21 (d, 1H, *J* = 18.3 Hz), 4.16 (d, 1H, *J* = 18.3 Hz), 4.12 (m, 1H), 2.20 - 1.98 (m, 3H), 1.88 - 1.66 (m, 3H), 1.60 (qd, 1H, *J* = 12.6, 3.2 Hz), 1.52 - 1.42 (m, 1H), 1.31 (t, 3H, *J* = 7.2 Hz). HRMS (ESI) calcd. for [M + H]^+^ C_14_H_19_F_2_N_4_O_2_ 313.1476 found 313.1468

### 2-Amino-6-(4,4-difluorocyclohexyl)-4-ethoxy-6,7-dihydro-5*H*-pyrrolo[3,4-*d*]pyrimidin-5-one (31)

Reaction of 6-(4,4-difluorocyclohexyl)-4-ethoxy-2-(methylthio)-6,7-dihydro-5*H*-pyrrolo[3,4-*d*]pyrimidin-5-one **(S29)** (101 mg, 0.29 mmol), oxone (361 mg, 0.59 mmol), MeCN (5 mL), water (2.5 mL) and NH_3_ (3 M in 1,4-dioxane, 2 mL) in an analogous manner to compound **6**, stirring the oxidation for 40 min and heating the displacement to 100 °C afforded **31** as a white solid (57 mg, 59%). ^1^H NMR (400 MHz, DMSO*-d_6_*) δ 7.21 (s, 2H), 4.40 (q, *J* = 7.0 Hz, 2H), 4.17 (s, 2H), 4.14 – 4.04 (m, 1H), 2.13 – 1.91 (m, 5H), 1.77 – 1.68 (m, 3H), 1.30 (t, *J* = 7.0 Hz, 3H). HRMS (ESI) calcd. for [M + H]^+^ C_14_H_19_O_2_N_4_F_2_ 313.1476 found 313.1479

### (*R*)-2-Amino-6-(2,2-difluorocyclohexyl)-4-methoxy-6,7-dihydro-5*H*-pyrrolo[3,4-*d*]pyrimidin-5-one (33)

Reaction of ethyl 4-formyl-6-methoxy-2-(methylthio)pyrimidine-5-carboxylate **(S21)** (373 mg, 1.45 mmol), (*R*)-2,2-difluorocyclohexan-1-amine hydrochloride (250 mg, 1.45 mmol), DIPEA (0.38 mL, 2.18 mmol) and sodium triacetoxyborohydride (926 mg, 4.37 mmol) in THF (14.5 mL) in an analogous manner to compound **S45,** with the addition of DIPEA (0.38 mL, 2.18 mmol) afforded (*R*)-6-(2,2-difluorocyclohexyl)-4-methoxy-2-(methylthio)-6,7-dihydro-5*H*-pyrrolo[3,4-*d*]pyrimidin-5-one **(S30)**. MS (ES+) m/z 259.1 [M + H]^+^ Reaction of **S30** (220 mg, 0.67 mmol), oxone (657 mg, 1.07 mmol), MeCN (4 mL), water (3 mL) and NH_3_ (3 M in 1,4-dioxane, 8 mL) in an analogous manner to compound **6** afforded **33** as a white solid (113 mg, 24% over two steps). ^1^H NMR (400 MHz, DMSO*-d_6_*) *δ* 7.31 (s, 2H), 4.48 – 4.11 (m, 3H), 3.91 (s, 3H), 2.02 – 1.63 (m, 6H), 1.61 – 1.26 (m, 2H). HRMS (ESI) calcd. for [M + H]^+^ C_13_H_17_F_2_N_4_O_2_ 299.1319 found 299.1314

### 2-Amino-6-((1R,2S)-2-aminocyclohexyl)-4-ethoxy-6,7-dihydro-5*H*-pyrrolo[3,4-*d*]pyrimidin-5-one (34)

*Tert*-butyl ((1*S*,2*R*)-2-(4-ethoxy-2-(methylthio)-5-oxo-5,7-dihydro-6*H*-pyrrolo[3,4-*d*]pyrimidin-6-yl)cyclohexyl)carbamate **(S31)** was synthesised from ethyl 4-ethoxy-6-formyl-2-(methylthio)pyrimidine-5-carboxylate **(S20)** (200 mg, 0.74 mmol), *tert*-butyl ((1*S*,2*R*)-2-aminocyclohexyl)carbamate (158 mg, 0.74 mmol) and sodium triacetoxyborohydride (470 mg, 2.22 mmol) in THF (4 mL) in an analogous manner to compound **S45** where the purification was performed using an SCX cartridge. MS (ES+) m/z 423.3 [M + H]^+^ Reaction of **S31** (240 mg, 0.57 mmol), oxone (698 mg, 1.14 mmol), MeCN (6 mL), water (3 mL) and NH_3_ (3 M in 1,4-dioxane, 3 mL) in an analogous manner to compound **6**, stirring the oxidation for 1 h and heating the displacement to 95 °C. The resulting product was dissolved in DCM (1 mL) added 4M HCl in dioxane (0.285 mL, 1.14 mmol) was added. The reaction was stirred at rt for 5 h, diluted with MeOH and loaded onto a SCX cartridge, eluting with 1M NH_3_ in MeOH, concentrated *in vacuo* and purified by prep-HPLC to afford **34** as a white solid (76 mg, 44% over three steps). ^1^H NMR (400 MHz, DMSO*-d_6_*) *δ* 7.09 (s, 2H), 4.60 (d, *J* = 18.6 Hz, 1H), 4.40 (qq, *J* = 6.9, 3.6 Hz, 2H), 4.16 (d, *J* = 18.6 Hz, 1H), 3.89 (dt, *J* = 12.7, 3.6 Hz, 1H), 3.16 (s, 1H), 1.99 – 1.84 (m, 1H), 1.78 – 1.59 (m, 2H), 1.57 – 1.41 (m, 4H), 1.38 – 1.24 (m, 5H). HRMS (ESI) calcd. for [M + H]^+^ C_14_H_22_O_2_N_5_ 292.1773 found 292.1772

### 2-Amino-6-(2,2-dimethylcyclohexyl)-4-methoxy-6,7-dihydro-5*H*-pyrrolo[3,4-*d*]pyrimidin-5-one (35)

Reaction of ethyl 2-((*tert*-butoxycarbonyl)amino)-4-formyl-6-methoxypyrimidine-5-carboxylate **(S37)** (210 mg, 0.67 mmol), 2,2-dimethylcyclohexan-1-amine (85 mg, 0.10 mmol) and sodium triacetoxyborohydride (286 mg, 1.35 mmol) in DCM (8 mL) in an analogous manner to compound **S45** afforded *tert*-butyl (6-(2,2-dimethylcyclohexyl)-4-methoxy-5-oxo-6,7-dihydro-5*H*-pyrrolo[3,4-*d*]pyrimidin-2-yl)carbamate **(S41).** MS (ES+) m/z 391.3 [M + H]^+^ To a solution of compound **S41** in DCM (5 mL) was added TFA (0.5 mL). The reaction was stirred at rt for 16 h, concentrated *in vacuo* and purified by prep-HPLC to afford compound **35** (133 mg, 64%, over two steps) as a white solid. ^1^H NMR (500 MHz, DMSO*-d_6_*) *δ* 7.19 (s, 2H), 4.28 – 4.10 (m, 2H), 3.89 (s, 4H), 1.87 – 1.73 (m, 2H), 1.37 (ddd, *J* = 38.1, 19.0, 7.2 Hz, 6H), 0.94 (s, 3H), 0.85 (s, 3H). HRMS (ESI) calcd. for [M + H]^+^ C_15_H_23_O_2_N_4_ 291.1821 found 291.1820

### 2-Amino-6-(2,2-difluorocycloheptyl)-4-methoxy-6,7-dihydro-5*H*-pyrrolo[3,4-*d*]pyrimidin-5-one (38)

To a solution of *tert*-butyl (6-(2,2-difluorocycloheptyl)-4-methoxy-5-oxo-6,7-dihydro-5*H*-pyrrolo[3,4-*d*]pyrimidin-2-yl)carbamate **(S42)** (430 mg, 1.04 mmol) in DCM (3 mL) was added TFA (0.32 mL, 4.17 mmol). The reaction stirred at rt for 16 h, concentrated *in vacuo* and loaded onto a SCX cartridge, eluting with 1M NH_3_ in MeOH solution and charily purified by SFC, utilising an isocratic gradient of 45% ACN+EtOH 1:1, 10 mL/min to afford compound **38** as a white solid.^1^H NMR (400 MHz, DMSO*-d_6_*) *δ* 7.34 (s, 2H), 4.61 – 4.41 (m, 1H), 4.33 (d, *J* = 18.2 Hz, 1H), 4.14 (d, *J* = 18.3 Hz, 1H), 3.91 (s, 3H), 2.27 – 1.94 (m, 2H), 1.94 – 1.41 (m, 8H). HRMS (ESI) calcd. for [M + H]^+^ C_14_H_19_F_2_N_4_O_2_ 313.1476 found 313.1469

### 2-Amino-6-((cis-4-hydroxycyclohexyl)-4-methoxy-7,7-dimethyl-6,7-dihydro-5*H*-pyrrolo[3,4-*d*]pyrimidin-5-one (39)

Reaction of ethyl 4-formyl-6-methoxy-2-(methylthio)pyrimidine-5-carboxylate **(S21)** (110 mg, 0.43 mmol), *cis*-4-aminocyclohexanol (59 mg, 0.52 mmol) and sodium triacetoxyborohydride (273 mg, 1.29 mmol) in DCM (8 mL) in an analogous manner compound **S45** where purification was achieved by chromatography eluting with 0 – 100% EtOAc in heptane afforded *cis*-4-hydroxycyclohexyl)-4-methoxy-2-(methylthio)-6,7-dihydro-5*H*-pyrrolo[3,4-*d*]pyrimidin-5-one **(S32)**. MS (ES+) m/z 310.4 [M + H]^+^ Compound **S32** (94 mg, 0.30 mmol) was suspended in DMF (3 mL) and cooled to 0°C. NaH (60% dispersion in mineral oil, 24 mg, 0.61 mmol) was added and the reaction stirred at rt for 30 min before the addition of MeI (107 mg, 0.76 mmol), the reaction allowed to warm to rt and stirred for 16 h. The reaction diluted with EtOAc and brine, the combined organics layers dried over Na_2_SO_4_, concentrated *in vacuo* and purified by flash chromatography (0 to 100% EtOAc in Heptane) to afford *cis*-4-hydroxycyclohexyl)-4-methoxy-7,7-dimethyl-2-(methylthio)-6,7-dihydro-5*H*-pyrrolo[3,4-*d*]pyrimidin-5-one **(S43).** MS (ES+) m/z 338.4 [M + H]^+^ Reaction of **S43** (76 mg, 0.23 mmol), oxone (166 mg, 0.27 mmol), MeCN (2.5 mL), water (1 mL) and NH_3_ (3 M in 1,4-dioxane, 2 mL) in an analogous manner to compound **6**, heating the displacement at 80 °C afforded **39** as a yellow solid (9 mg, 7% over three steps). ^1^H NMR (400 MHz, DMSO*-d_6_*) *δ* 7.25 (s, 2H), 4.24 (s, 1H), 3.87 (s, 3H), 3.81 (s, 1H), 3.23 – 3.09 (m, 1H), 2.66 (q, *J* = 11.5 Hz, 2H), 1.71 (d, *J* = 13.3 Hz, 2H), 1.46 (t, *J* = 13.2 Hz, 2H), 1.32 (s, 6H), 1.23 (d, *J* = 12.0 Hz, 2H). HRMS (ESI) calcd. for [M + H]^+^ C_15_H_23_N_4_O_3_ 307.1770 found 307.1775

### 6-Amino-2-cycloheptyl-4-ethoxy-1,2-dihydro-3H-pyrazolo[3,4-*d*]pyrimidin-3-one (40)

Reaction of 2-cycloheptyl-4-ethoxy-6-(methylthio)-1,2-dihydro-3*H*-pyrazolo[3,4-*d*]pyrimidin-3-one **(S59)** (85 mg, 0.26 mmol), oxone (163 mg, 0.26 mmol), MeCN (0.8 mL), H_2_O (0.5 mL) and NH_3_ (3 M in 1,4-dioxane, 2 mL) in an analogous manner to compound **6**, stirring the oxidation step for 50 min and the heating the displacement to 95 °C afforded **40** (4 mg, 4%) as a white solid. ^1^H NMR (500 MHz, DMSO*-d_6_*) *δ* 10.55 (s, 1H), 7.19 – 6.99 (m, 2H), 6.87 (s, 2H), 4.39 (q, *J* = 7.0 Hz, 3H), 4.18 (s, 2H), 1.82 – 1.63 (m, 9H), 1.63 – 1.54 (m, 3H), 1.54 – 1.37 (m, 6H), 1.30 (t, *J* = 7.1 Hz, 4H). HRMS (ESI) calcd for [M + H]^+^ C_14_H_22_O_2_N_5_ 292.1774 found 292.1774

### 6-Amino-2-cycloheptyl-1-(cyclopropylmethyl)-4-ethoxy-1,2-dihydro-3H-pyrazolo[3,4-*d*]pyrimidin-3-one (41)

To a solution of 2-cycloheptyl-4-ethoxy-6-(methylthio)-1,2-dihydro-3*H*-pyrazolo[3,4-*d*]pyrimidin-3-one **(S59)** (50 mg, 0.16 mmol) in THF (1 mL) was added K_2_CO_3_ (21 mg, 0.16 mmol). The reaction sealed, heated at 90 °C for 1 h, cooled to rt, bromomethylcyclopropane (21 mg, 0.16 mmol) added, the reaction re-sealed and heated at 90 °C for a further 6 h. The residue was purified by prep-HPLC to afford 2-cycloheptyl-1-(cyclopropylmethyl)-4-ethoxy-6-(methylthio)-1,2-dihydro-3*H*-pyrazolo[3,4-*d*]pyrimidin-3-one **(S60)**. MS (ES+) m/z 377.2 [M + H]^+^ Reaction of **S60** (50 mg, 0.13 mmol), oxone (82 mg, 0.13 mmol), MeCN (0.8 mL), water (0.5 mL) and NH_3_ (3 M in 1,4-dioxane, 2 mL) in an analogous manner to compound **6**, heating the displacement at 95 °C for 90 min afforded **41** as a white solid (6 mg, 12% over two steps). ^1^H NMR (500 MHz, DMSO*-d_6_*) *δ* 7.04 (s, 2H), 4.38 (q, *J* = 7.1 Hz, 2H), 3.89 – 3.80 (m, 1H), 3.67 (d, *J* = 6.8 Hz, 2H), 2.11 – 1.99 (m, 2H), 1.78 – 1.66 (m, 4H), 1.63 – 1.55 (m, 2H), 1.54 – 1.47 (m, 2H), 1.45 – 1.36 (m, 2H), 1.31 (t, *J* = 7.1 Hz, 3H), 0.94 – 0.82 (m, 1H), 0.41 – 0.32 (m, 2H), 0.25 –0.16 (m, 2H). HRMS (ESI) calcd. for [M + H]+ C_18_H_28_N_5_O_2_ 346.2243 found 346.2242

### 6-Amino-1-(2-(azetidin-3-yl)ethyl)-2-cycloheptyl-4-ethoxy-1,2-dihydro-3H-pyrazolo[3,4-*d*]pyrimidin-3-one (42)

To a solution of 2-cycloheptyl-4-ethoxy-6-(methylthio)-1,2-dihydro-3*H*-pyrazolo[3,4-*d*]pyrimidin-3-one **(S59)** (180 mg, 0.56 mmol) in acetone (3 mL) was added *tert-*butyl 3-(2-iodoethyl)azetidine-1-carboxylate (174 mg, 0.56 mmol) and K_2_CO_3_ (77 mg, 0.56 mmol). The reaction heated under µW conditions at 100 °C for 20 min followed by 120 °C for 45 min, filtered, concentrated *in vacuo* and purified by flash chromatography (0-100 % EtOAc in heptane) to afford *tert*-butyl 3-(2-(2-cycloheptyl-4-ethoxy-6-(methylthio)-3-oxo-2,3-dihydro-1*H*-pyrazolo[3,4-*d*]pyrimidin-1-yl)ethyl)azetidine-1-carboxylate **(S61)**. Reaction of **S61**, oxone (343 mg, 0.56 mmol), MeCN (2 mL), H_2_O (1mL) and NH_3_ (3 M in 1,4-dioxane, 1 mL) in an analogous manner to compound **6**, running the oxidation for 3 h and the heating the displacement at 95 °C for 3 h where the resultant residue was dissolved in DCM and TFA (0.5 mL) added, the reaction stirred at rt for 2 h, concentrated *in vacuo* and purified by prep-HPLC afforded **42** as a white solid (25 mg, 12% yield over 2 steps). ^1^H NMR (400 MHz, DMSO) *δ* 7.08 (s, 4H), 4.38 (q, *J* = 7.1 Hz, 5H), 3.97 – 3.80 (m, 3H), 3.80 – 3.60 (m, *J* = 6.8 Hz, 6H), 3.46 – 3.30 (m, 8H), 3.08 (s, 4H), 2.08 – 1.89 (m, *J* = 20.7, 10.3 Hz, 5H), 1.78 – 1.35 (m, 28H), 1.30 (t, *J* = 7.1 Hz, 7H). HRMS (ESI) calcd. for [M + H]+ C_19_H_31_O_2_N_6_ 375.2508 found 375.2507

### 6-Cyclohexyl-4-ethoxy-2-(methylamino)-6,7-dihydro-5*H*-pyrrolo[3,4-*d*]pyrimidin-5-one (43)

To a solution of 6-cyclohexyl-4-ethoxy-2-(methylthio)-6,7-dihydro-5*H*-pyrrolo[3,4-*d*]pyrimidin-5-one **(S33)** (200 mg, 650 µmol) in MeCN (4 mL) was added Oxone (799 mg, 1.30 mmol). The reaction was stirred at 25 °C for 0.5 h, methanamine (2 M in THF, 650 µL) added and the reaction stirred at 25 °C for 12 h, poured into aq.HCl (1M 30 mL), extracted with EtOAc, the combined organics layers washed with brine, dried over Na_2_SO_4_, concentrated *in vacuo* and purified by prep-HPLC to afford compound **43** (35.43 mg, 19% as the formic acid salt) as a white solid. ^1^H NMR (400 MHz, DMSO*-d_6_*) *δ* 7.60 (br d, *J* = 19.2 Hz, 1H), 4.54 - 4.43 (m, 1H), 4.41 - 4.35 (m, 1H), 4.23 -4.10 (m, 2H), 3.93 - 3.82 (m, 1H), 2.85 (s, 3H), 1.77 (br d, *J* = 12.4 Hz, 2H), 1.71 -1.58 (m, 3H), 1.51 - 1.40 (m, 2H), 1.39-1.26 (m, 5H), 1.21 - 1.03 (m, 1H). HRMS (ESI) calcd for [M + H]+ HRMS (ESI) calcd. for [M + H]+ C_15_H_23_O_2_N_4_ 291.1821 found 291.1820

### 6-Cyclohexyl-2-(cyclopropylamino)-4-methoxy-6,7-dihydro-5*H*-pyrrolo[3,4-*d*]pyrimidin-5-one (44)

To a solution of 6-cyclohexyl-4-methoxy-2-(methylthio)-6,7-dihydro-5*H*-pyrrolo[3,4-*d*]pyrimidin-5-one **(S34)** (310 mg, 1.06 mmol) in MeCN (10 mL) and water (1 mL) was added Oxone (780 mg, 1.27 mmol). The reaction stirred at rt for 16 h, diluted with water and DCM, passed through hydrophobic frit and concentrated *in vacuo*. The residue was suspended in 1,4-dioxane (3 mL) and aminocyclopropane (34 mg, 0.59 mmol) and DIPEA (0.11 mL, 0.65 mmol) were added. The reaction was stirred at 80 °C for 3 h, concentrated *in vacuo* and purified by prep-HPLC to afford compound **44** (43 mg, 40%) as a white solid. ^1^H NMR (400 MHz, DMSO*-d_6_*) *δ* 7.85 (br s, 1H), 4.20 (br s, 2H), 4.04 – 3.80 (m, 4H), 2.88 – 2.76 (m, 1H), 1.77 (d, *J* = 12.7 Hz, 2H), 1.71 – 1.55 (m, 3H), 1.53 – 1.24 (m, 4H), 1.21 – 1.03 (m, 1H), 0.69 (d, *J* = 6.7 Hz, 2H), 0.56 – 0.46 (m, 2H). HRMS (ESI) calcd. for [M + H]^+^ C_16_H_23_O_2_N_4_ 303.1821 found 303.1821

### 2-(Cyclopropylamino)-6-((2S)-2-hydroxycyclohexyl)-4-methoxy-6,7-dihydro-5*H*-pyrrolo[3,4-*d*]pyrimidin-5-one (45)

Reaction of 6-((2*S*)-2-hydroxycyclohexyl)-4-methoxy-2-(methylthio)-6,7-dihydro-5*H*-pyrrolo[3,4-*d*]pyrimidin-5-one **(S35)** (74 mg, 0.23 mmol), aminocyclopropane (19 mg, 0.34 mmol), DIPEA (0.08 mL) and 1,4-dioxane (2.5 mL) in an analogous manner to compound **44** afforded **45** a white solid (38 mg, 50%). ^1^H NMR (500 MHz, DMSO*-d_6_*) *δ* 7.86 (d, *J* = 43.1 Hz, 1H), 4.66 (s, 1H), 4.58 – 4.37 (m, 1H), 4.37 – 4.09 (m, 1H), 4.05 – 3.80 (m, 5H), 2.82 (dq, *J* = 7.3, 3.6 Hz, 1H), 2.00 – 1.81 (m, 1H), 1.81 – 1.63 (m, 2H), 1.63 – 1.39 (m, 3H), 1.39 – 1.27 (m, 2H), 0.76 – 0.62 (m, 2H), 0.55 – 0.43 (m, 2H). HRMS (ESI) calcd. for [M + H]^+^ C_16_H_22_N_4_O_3_ 319.1770 found 319.1776

### 2-Amino-6-((1S,7S)-2,2-difluoro-7-hydroxycycloheptyl)-4-ethoxy-6,7-dihydro-5*H*-pyrrolo[3,4-*d*]pyrimidin-5-one (48)

Reaction of 6-((1*S*,7*S*)-2,2-difluoro-7-hydroxycycloheptyl)-4-ethoxy-2-(methylthio)-6,7-dihydro-5*H*-pyrrolo[3,4-*d*]pyrimidin-5-one **(S36)** (84 mg, 0.22 mmol), oxone (166 mg, 0.27 mmol), MeCN (3 mL), water (1 mL) and NH_3_ (0.5 M in 1,4-dioxane, 3 mL) in an analogous manner to compound **6**, afforded **48** as a white solid (44 mg, 55%). ^1^H NMR (500 MHz, DMSO*-d_6_*) *δ* 7.26 (s, 2H), 5.20 (d, *J* = 4.2 Hz, 1H), 4.62 – 4.50 (m, 2H), 4.43 (q, *J* = 7.0 Hz, 2H), 4.34 (d, *J* = 18.7 Hz, 1H), 4.07 – 3.99 (m, 1H), 2.30 – 2.10 (m, 2H), 1.98 – 1.85 (m, 1H), 1.83 – 1.56 (m, 4H), 1.52 – 1.41 (m, 1H), 1.33 (t, *J* = 7.0 Hz, 3H). HRMS (ESI) calcd. for [M + H]^+^ C_15_H_21_F_2_N_4_O_3_ 343.1581 found 343.1585

### Ethyl 4-chloro-6-(2-methoxyethoxy)-2-(methylthio)pyrimidine-5-carboxylate (S4)

A solution of 2-methoxyethanol (0.21 mL, 2.62 mmol) and NaH (60% dispersion in mineral oil, 110 mg, 2.76 mmol) in THF (12 mL), cooled to 0 °C was added dropwise to solution of ethyl 4,6-dichloro-2-(methylthio)pyrimidine-5-carboxylate **(S2)** (702 mg, 2.63 mmol), in THF (60 mL). The reaction was stirred at 0 °C for 5 min, the volatiles concentrated *in vacuo* and the residue partitioned between EtOAc and water. The combined organic layers were dried over Na_2_SO_4_, concentrated *in vacuo* and purified by column chromatography (0 – 20% EtOAc in heptane) to afford compound **S4** as a colourless oil. The material was combined with previous reaction that was performed on a smaller scale to afford compound **S4** (569 mg, 28%). ^1^H NMR (500 MHz, CDCl_3_) *δ* 4.63 – 4.50 (m, 2H), 4.39 (q, *J* = 7.1 Hz, 2H), 3.78 – 3.63 (m, 2H), 3.39 (s, 3H), 2.54 (s, 3H), 1.37 (t, *J* = 7.1 Hz, 3H). MS (ES+) m/z 307.2 [M + H]+

### Ethyl 4-chloro-6-(2,2-difluoroethoxy)-2-(methylthio)pyrimidine-5-carboxylate (S5)

Reaction of ethyl 4,6-dichloro-2-(methylthio)pyrimidine-5-carboxylate **(S2)** (1014 mg, 3.79 mmol), 2,2-difluoroethanol (0.24 mL, 3.79 mmol), NaH (60% dispersion in mineral oil, 159 mg, 3.98 mmol) and THF (82 mL) in an analogous manner to compound **S4**, afforded **S5** as a colourless oil (837 mg, 71%). ^1^H NMR (500 MHz, CDCl_3_) *δ* 6.06 (tt, *J* = 55.0, 4.1 Hz, 1H), 4.62 (td, *J* = 13.1, 4.1 Hz, 2H), 4.40 (q, *J* = 7.1 Hz, 2H), 2.56 (s, 3H), 1.37 (t, *J* = 7.1 Hz, 3H). MS (ES+) m/z 313.3 [M + H]^+^

### Ethyl 4-(2-methoxyethoxy)-2-(methylthio)-6-vinylpyrimidine-5-carboxylate (S7)

To a solution of ethyl 4-chloro-6-(2-methoxyethoxy)-2-(methylthio)pyrimidine-5-carboxylate **(S4)** (569 mg, 1.85 mmol) in EtOH (20 mL) was added potassium trifluoro(vinyl)boranuide (469 mg, 3.7 mmol), Pd(dppf)_2_Cl_2_ (65 mg, 0.09 mmol), TEA (0.42 mL, 2.41 mmol), the reaction purged with N_2_ and sealed. The reaction was stirred at reflux for 16 h, concentrated *in vacuo*, partitioned between EtOAc and water, the combined organic layers dried over Na_2_SO_3_, concentrated *in vacuo* and purified by column chromatography (0 – 30% EtOAc in heptane) to afford **S7** as an orange oil clear, colourless oil (424 mg, 76%). ^1^H NMR (500 MHz, CDCl_3_) *δ* 6.83 (dd, *J* = 16.8, 10.4 Hz, 1H), 6.65 (dd, *J* = 16.8, 2.0 Hz, 1H), 5.65 (dd, *J* = 10.5, 2.0 Hz, 1H), 4.59 – 4.50 (m, 2H), 4.38 (q, *J* = 7.1 Hz, 2H), 3.77 – 3.66 (m, 2H), 3.40 (s, 3H), 2.57 (s, 3H), 1.37 (t, *J* = 7.1 Hz, 3H). MS (ES+) m/z 299.2 [M + H]+

### Ethyl 4-(2,2-difluoroethoxy)-2-(methylthio)-6-vinylpyrimidine-5-carboxylate (S8)

Reaction of ethyl 4-chloro-6-(2,2-difluoroethoxy)-2-(methylthio)pyrimidine-5-carboxy **(S5)** (837 mg, 2.68 mmol), potassium trifluoro(vinyl)boranuide (717 mg, 5.35 mmol), TEA (0.61 mL, 3.48 mmol) and EtOH (50 mL) in an analogous manner to compound **S7** to afford **S8** as a colourless oil (649 mg, 80%). ^1^H NMR (500 MHz, CDCl_3_) *δ* 6.87 (dd, *J* = 16.8, 10.4 Hz, 1H), 6.69 (dd, *J* = 16.8, 1.9 Hz, 1H), 6.07 (tt, *J* = 55.2, 4.2 Hz, 1H), 5.70 (dd, *J* = 10.5, 2.0 Hz, 1H), 4.60 (td, *J* = 13.2, 4.2 Hz, 2H), 4.39 (q, *J* = 7.1 Hz, 2H), 2.58 (s, 3H), 1.37 (t, *J* = 7.2 Hz, 3H). MS (ES+) m/z 305.5 [M + H]^+^

### Ethyl 4-chloro-2-(methylthio)-6-vinylpyrimidine-5-carboxylate (S9)

To a solution of ethyl 4-chloro-2-(methylthio)pyrimidine-5-carboxylate **(S16)** (150 g, 644 mmol) in THF (1 L), cooled to 0 °C was added vinylmagnesiumbromide (1 M in THF, 967 mL) dropwise. The reaction was stirred at 20 °C for 30 min, quenched by the dropwise addition of acetone (38 mL, 515 mmol) at 0 °C and concentrated *in vacuo.* The residue was dissolved in DCM (1.5 L) and MeOH (450 mL) and PhI(OAc)_2_ (311.5 g, 967 mmol) was added. The reaction was stirred at 20 °C for 12 h, poured into NaHCO_3(aq)_ (2 L) and stirred for 1 h, extracted with DCM and the combined organic layers were washed with brine, dried over Na_2_SO_4_ concentrated *in vacuo* and purified by column chromatography (0 – 2 % EtOAc in heptane) to afford compound **S9** (135 g, 40%) as a brown oil. ^1^H NMR (400 MHz, CDCl_3_) *δ* 6.79 - 6.67 (m, 2H), 5.77 (dd, *J* = 9.3, 2.9 Hz, 1H), 4.45 (q, *J* = 7.1 Hz, 2H), 2.60 (s, 3H), 1.41 (t, *J* = 7.2 Hz, 3H). MS *m/z* 259.0 [M+H]^+^.

### Ethyl 4-(difluoromethoxy)-2-(methylthio)-6-vinylpyrimidine-5-carboxylate (S10)

Reaction of ethyl 4-chloro-6-(difluoromethoxy)-2-(methylthio)pyrimidine-5-carboxylate **(S19)** (1000 mg, 3.35 mmol) in EtOH (30 mL) was added potassium trifluoro(vinyl)boranuide (896 mg, 6.69 mmol), Pd(dppf)_2_Cl_2_ (117 mg, 0.17 mmol), TEA (0.76 mmol, 4.35 mmol), in an analogous manner to compound **S7** afforded **S10** as an orange oil (683 mg, 67%). ^1^H NMR (400 MHz, CDCl_3_) *δ* 7.50 (d, *J* = 69.0 Hz, 1H), 6.93 (dd, *J* = 16.8, 10.4 Hz, 1H), 6.75 (dd, *J* = 16.7, 1.9 Hz, 1H), 5.77 (dt, *J* = 10.4, 1.2 Hz, 1H), 4.44 (q, *J* = 7.1 Hz, 2H), 2.60 (d, *J* = 0.8 Hz, 3H), 1.41 (t, *J* = 7.1 Hz, 3H). MS (ES+) m/z 291.2 [M + H]^+^

### Ethyl 4-formyl-2-(methylthio)-6-propoxypyrimidine-5-carboxylate (S11)

To a solution of propan-1-ol (0.35 mL, 1.50 mmol) in THF (10 mL) cooled to 0 °C was added NaH (60% dispersion in mineral oil, 63 mg, 1.57 mmol). The reaction was stirred at 0 °C for 15 min and ethyl 4,6-dichloro-2-(methylthio)pyrimidine-5-carboxylate **(S2)** (400 mg, 1.49 mmol) was added as a solution in THF (5 mL). The reaction was stirred at 0 °C for 10 min, diluted with DCM and water, passed through a hydrophobic frit, concentrated *in vacuo* and purified by flash chromatography (0 - 20% EtOAc in heptane) to afford ethyl 4-chloro-2-(methylthio)-6-propoxypyrimidine-5-carboxylate **(S3)**. MS (ES+) m/z 291.2 [M + H]^+^ **S3** (535 mg, 1.84 mmol) was dissolved in EtOH (3 mL) and water (3 mL), potassium trifluoro(vinyl)boranuide (246 mg, 1.84 mmol), Pd(dppf)_2_Cl_2_ (65 mg, 0.092 mmol), TEA (0.42 mL, 2.39 mmol) were added, the reaction purged with N_2_ and sealed. The reaction was stirred at 100 °C for 2 h, diluted with DCM and washed with 1 M HCl and then sat NaHCO_3_, the combined organic layers were passed through a hydrophobic frit, concentrated *in vacuo* and purified by column chromatography (0-10% EtOAc in heptane) to afford ethyl 2-(methylthio)-4-propoxy-6-vinylpyrimidine-5-carboxylate **(S6).** MS (ES+) m/z 283.2 [M + H]^+^ **S6** (500 mg, 1.77 mmol) was added to a cooled solution of OsO_4_ (2.5% solution in *tert*-BuOH, 0.0154 mL, 0.062 mmol) in 1,4-dioxane (40 mL) and water (5 mL) in an ice bath, and NaIO_4_ (1515 mg, 7.08 mmol) and 2,6-lutidine (0.297 mL, 2.56 mmol) were added. The reaction stirred at rt for 3 h, diluted with EtOAc and water, the combined organic layers were dried over Na_2_SO_4_, concentrated *in vacuo* and purified by flash chromatography (0 - 20% EtOAc in heptane) to afford compound **S11** (162 mg, 27% over three steps) as a colourless oil. ¹H NMR (500 MHz, CD_3_OD) *δ* 7.02 (s, 1H), 6.30 (s, 2H), 5.93 - 5.82 (m, 4H), 4.08 (s, 3H), 2.85 (dd, *J* = 7.1, 7.1 Hz, 3H), 2.52 (dd, *J* = 7.5, 7.5 Hz, 3H). MS (ES+) m/z 285.3 [M + H]^+^

### Ethyl 4-formyl-6-(2-methoxyethoxy)-2-(methylthio)pyrimidine-5-carboxylate (S12)

To a solution of ethyl 4-(2-methoxyethoxy)-2-(methylthio)-6-vinylpyrimidine-5-carboxylate **(S7)** (424 mg, 1.42 mmol) in 1,4-dioxane (5 mL) and water (0.5 mL) was added of NaIO_4_ (912 mg, 4.26 mmol) and dropwise addition of OsO_4_ (4% solution in water, 0.86 mL, 0.14 mmol). The reaction was stirred at rt for 16 h, diluted with EtOAc and water, the combined organics were combined washed with brine, dried over MgSO_4_, filtered, concentrated *in vacuo* and purified by flash chromatography (0 - 20% EtOAc in heptane) to afford **S12** (154 mg, 36%) as a colourless oil. ^1^H NMR (400 MHz, CDCl_3_) *δ* 9.89 (s, 1H), 4.67 – 4.55 (m, 2H), 4.42 (q, *J* = 7.1 Hz, 2H), 3.80 – 3.63 (m, 2H), 3.40 (s, 3H), 2.60 (s, 3H), 1.36 (t, *J* = 7.1 Hz, 3H). MS (ES+) m/z 301.3 [M + H]+

### Ethyl 4-(2,2-difluoroethoxy)-6-formyl-2-(methylthio)pyrimidine-5-carboxylate (S13)

Reaction of ethyl 4-(2,2-difluoroethoxy)-2-(methylthio)-6-vinylpyrimidine-5-carboxylate **(S8)** (649 mg, 2.13 mmol), OsO_4_ (4% solution in water, 0.65 mL, 0.11 mmol), NaIO_4_ (1369 mg, 6.39 mmol), 1,4-dioxane (10 mL) and water (1 mL) in an analogous manner to compound **S12** afforded **S13** as a yellow oil (161 mg, 25%). ^1^H NMR (500 MHz, CDCl_3_) *δ* 9.91 (s, 1H), 6.09 (tt, *J* = 55.0, 4.2 Hz, 1H), 4.65 (td, *J* = 13.0, 4.1 Hz, 2H), 4.43 (q, *J* = 7.1 Hz, 2H), 2.62 (s, 3H), 1.37 (t, *J* = 7.1 Hz, 3H). MS (ES+) m/z 307.3 [M + H]^+^

### Ethyl 4-chloro-6-formyl-2-(methylthio)pyrimidine-5-carboxylate (S14)

To a solution of ethyl 4-chloro-2-(methylthio)-6-vinylpyrimidine-5-carboxylate **(S9)** (20 g, 77.30 mmol) in 1,4-dioxane (500 mL) and water (75 mL) was added of OsO_4_ (4 wt. % in water, 401 µL, 7.73 mmol) and a solution of 2,6-lutidine (22.5 mL, 193.26 mmol) in water (75 mL) dropwise at 0 °C. The reaction stirred at 10 °C for 5 min, NaIO_4_ (83 g, 386.51 mmol) was added batch wise and the reaction was stirred at rt for 2 h. The reaction was poured into ice-water (w/w = 1/1) (2000 mL) and stirred for 30 min, extracted with EtOAc and the combined organics were washed with brine, dried over Na_2_SO_4_, concentrated *in vacuo* and purified by column chromatography (0 – 5% EtOAc in petroleum ether) to afford **S14**. The reaction was repeated in parallel on the same scale three additional times to afford **S14** (32.45 g, 39%) as a colourless oil. ^1^H NMR (400 MHz, CDCl_3_) *δ* 9.89 (s, 1H), 4.48 (q, *J* = 7.2 Hz, 2H), 2.65 (s, 3H), 1.40 (t, *J* = 7.2 Hz, 3H). MS m/z [M+H]^+^ 261.0.

### 4-Chloro-6-(difluoromethoxy)-2-(methylthio)pyrimidine (S18)

To a solution of 6-chloro-2-(methylthio)pyrimidin-4(3*H*)-one **(S17)** (5300 mg, 30 mmol) in MeCN (250 mL) and DMF (25 mL) was added Na_2_CO_3_ (6361 mg, 60 mmol) and sodium chlorodifluoroacetic acid (6862 mg, 45 mmol). The reaction was stirred at 90 °C for 16 h, diluted with water and EtOAc, the combined organics layers were washed with brine, concentrated *in vacuo* and purified by flash chromatography (0 to 50% EtOAc in heptane) to afford **S18** (6800 mg, 88%). ^1^H NMR (400 MHz, CDCl_3_) *δ* 7.46 (t, *J* = 71.3 Hz, 1H), 6.57 (d, *J* = 0.7 Hz, 1H), 2.55 (s, 4H).

### Ethyl 4-chloro-6-(difluoromethoxy)-2-(methylthio)pyrimidine-5-carboxylate (S19)

To a solution of 2,2,6,6 tetramethyl piperidine (7.9 mL, 46.861 mmol) in THF (150 mL), cooled to -78 °C was added *n*-BuLi (2.5 M in hexanes, 18.7 mL, 46.86 mmol). The reaction stirred at -78 °C for 1 h, a solution of 4-chloro-6-(difluoromethoxy)-2-(methylthio)pyrimidine **(S18)** (5900 mg, 26.03 mmol) in THF (50 mL) added, the reaction stirred at -78 °C for 1 h and ethyl carbonochloridate (4.48 mL, 46.86 mmol) added. The reaction was warmed to rt and stirred for 16 h, quenched by the addition of water, extracted with EtOAc and the combined organic layers were dried over Na_2_SO_4_, concentrated *in vacuo* and purified by flash chromatography (0 to 20% EtOAc in heptane) to afford compound **S19** (2600 mg, 33%) as a yellow oil. ^1^H NMR (400 MHz, CDCl_3_) *δ* 7.46 (t, *J* = 70.9 Hz, 2H), 4.45 (q, *J* = 7.1 Hz, 2H), 2.59 (s, 3H), 1.42 (t, *J* = 7.1 Hz, 3H).

### 6-cyclohexyl-4-(difluoromethoxy)-2-(methylthio)-6,7-dihydro-5*H*-pyrrolo[3,4-*d*]pyrimidin-5-one (S25)

Ethyl 4-(difluoromethoxy)-2-(methylthio)-6-vinylpyrimidine-5-carboxylate **(S10)** (683 mg, 2.35 mmol), OsO_4_ (4% solution in water, 0.5 mL, 0.08 mmol), NaIO_4_ (2013 mg, 9.41 mmol), 2,6-lutidine (0.55 mL, 4.7 mmol), 1,4-dioxane (15 mL) and water (2 mL) in an analogous manner to compound **S12** to afford ethyl 4-(difluoromethoxy)-6-formyl-2-(methylthio)pyrimidine-5-carboxylate **(S15)** as a brown oil. MS (ES+) m/z 293.2 [M + H]^+^ **S15** (230 mg, 0.79 mmol), cyclohexanamine (0.09 mL, 0.79 mmol) and sodium triacetoxyborohydride (500 mg, 2.36 mmol) in DCM (6 mL) in an analogous manner to compound **S45** afforded **S25** as a white solid (90 mg, 33% over two steps). ^1^H NMR (400 MHz, CDCl_3_) *δ* 7.61 (t, *J* = 70.9 Hz, 1H), 4.31 (s, 2H), 4.27 – 4.12 (m, 1H), 2.59 (s, 3H), 1.81 (d, *J* = 13.7 Hz, 4H), 1.73 (d, *J* = 13.4 Hz, 1H), 1.56 – 1.33 (m, 4H), 1.25 – 1.07 (m, 1H). MS (ES+) m/z 330.4 [M + H]^+^

### 6-Cycloheptyl-2-(methylthio)-4-propoxy-6,7-dihydro-5*H*-pyrrolo[3,4-*d*]pyrimidin-5-one (S22)

Reaction of ethyl 4-formyl-2-(methylthio)-6-propoxypyrimidine-5-carboxylate (**S11)** (162 mg, 0.57 mmol), cycloheptanamine (0.1 mL, 0.85 mmol), sodium triacetoxyborohydride (362 mg, 1.7 mmol) and THF (10 mL) in an analogous manner to compound **S45** afforded **S22** as a white solid (74 mg, 37%). ^1^H NMR (500 MHz, DMSO*-d_6_*) *δ* 4.46 - 4.39 (m, 4H), 4.16 - 4.11 (m, 1H), 2.57 (s, 3H), 1.79 - 1.45 (m, 14H), 0.98 (t, *J* = 7.4 Hz, 3H). MS (ES+) m/z 336.5 [M + H]+

### 6-Cyclohexyl-4-(2-methoxyethoxy)-2-(methylthio)-6,7-dihydro-5*H*-pyrrolo[3,4-*d*]pyrimidin-5-one (S23)

Reaction of ethyl 4-formyl-6-(2-methoxyethoxy)-2-(methylthio)pyrimidine-5-carboxylate **(S12)** (154 mg, 0.51 mmol), cyclohexylamine (0.09 mL, 0.77 mmol), sodium triacetoxyborohydride (326 mg, 1.54 mmol) and THF (10 mL) in an analogous manner to **S45** afforded **S23** as a white solid (136 mg, 79%). ^1^H NMR (500 MHz, CDCl_3_) *δ* 4.74 – 4.63 (m, 2H), 4.23 (s, 2H), 4.22 – 4.14 (m, 1H), 3.85 – 3.78 (m, 2H), 3.46 (s, 3H), 2.59 (s, 3H), 1.88 – 1.78 (m, 4H), 1.74 – 1.67 (m, 1H), 1.51 – 1.36 (m, 4H), 1.22 – 1.10 (m, 1H). MS (ES+) m/z 338.1 [M + H]^+^

### 6-Cyclohexyl-4-(2,2-difluoroethoxy)-2-(methylthio)-6,7-dihydro-5*H*-pyrrolo[3,4-*d*]pyrimidin-5-one (S24)

Reaction of ethyl 4-(2,2-difluoroethoxy)-6-formyl-2-(methylthio)pyrimidine-5-carboxylate **(S13)** (161 mg, 0.53 mmol), cyclohexanamine (0.09 mL, 0.79 mmol), sodium triacetoxyborohydride (334 mg, 1.58 mmol) and THF (10 mL) in analogous manner to compound **S45** afforded **S24** as a white solid (134 mg, 74%). ^1^H NMR (400 MHz, CDCl_3_) *δ* 6.21 (tt, *J* = 55.2, 4.3 Hz, 1H), 4.75 (td, *J* = 12.9, 4.4 Hz, 2H), 4.26 (s, 2H), 4.19 (s, 1H), 2.59 (s, 3H), 1.89 – 1.79 (m, 3H), 1.75 – 1.65 (m, 1H), 1.52 – 1.06 (m, 6H). MS (ES+) m/z 344.2 [M + H]^+^

### 6-(3,3-Difluorocyclohexyl)-4-methoxy-2-(methylthio)-6,7-dihydro-5*H*-pyrrolo[3,4-*d*]pyrimidin-5-one (S28)

Reaction of ethyl 4-ethoxy-6-formyl-2-(methylthio)pyrimidine-5-carboxylate **(S20)** (750 mg, 2.77 mmol), 3,3-difluorocyclohexan-1-amine (583 mg, 3.40 mmol) and sodium triacetoxyborohydride (1764 mg, 8.32 mmol) in DCM (25 mL) for 16 h in an analogous manner to compound **S45** afforded **S28** as a white solid (860 mg, 86%). ^1^H NMR (400 MHz, DMSO*-d_6_*) *δ* 4.54 (q, *J* = 7.0 Hz, 2H), 4.41 (d, *J* = 3.9 Hz, 2H), 4.21 – 4.09 (m, 1H), 2.57 (s, 3H), 2.26 – 2.09 (m, 2H), 2.09 – 1.98 (m, 1H), 1.92 – 1.81 (m, 1H), 1.81 – 1.56 (m, 3H), 1.56 – 1.40 (m, 1H), 1.35 (t, *J* = 7.1 Hz 3H). MS (ES+) m/z 344.3 [M + H]^+^

### 6-(4,4-Difluorocyclohexyl)-4-ethoxy-2-(methylthio)-6,7-dihydro-5*H*-pyrrolo[3,4-*d*]pyrimidin-5-one (S29)

Reaction of ethyl 4-ethoxy-6-formyl-2-(methylthio)pyrimidine-5-carboxylate **(S20)** (100 mg, 0.37 mmol), 4,4-difluorocyclohexanamine (75 mg, 0.55 mmol) and sodium triacetoxyborohydride (235 mg, 1.11 mmol) in THF (10 mL) in an analogous manner to compound **S45** afforded **S29** as a white solid (101 mg, 80 %). ^1^H NMR (400 MHz, CDCl_3_) *δ* 4.63 (q, *J* = 7.1 Hz, 2H), 4.41 – 4.27 (m, 1H), 4.22 (s, 2H), 2.59 (s, 3H), 2.31 – 2.12 (m, 2H), 2.04 – 1.70 (m, 6H), 1.48 (t, *J* = 7.1 Hz, 3H). MS (ES+) m/z 344.2 [M + H]^+^

### 6-((1S,7S)-2,2-Difluoro-7-hydroxycycloheptyl)-4-ethoxy-2-(methylthio)-6,7-dihydro-5*H*-pyrrolo[3,4-*d*]pyrimidin-5-one. (S36)

Reaction of ethyl 4-ethoxy-6-formyl-2-(methylthio)pyrimidine-5-carboxylate **(S20)** (178 mg, 0.66 mmol), (1*S*,2*S*)-2-amino-3,3-difluoro-cycloheptanol^1^ (109 mg, 0.66 mmol) and sodium triacetoxyborohydride (419 mg, 1.97 mmol) in THF (5 mL) in analogous manner to compound **S45,** heating the cyclisation to reflux, afforded **S36** as a white solid (85 mg, 33%). ^1^H NMR (400 MHz, DMSO*-d_6_*) *δ* 5.24 (d, *J* = 4.2 Hz, 1H), 4.74 (d, *J* = 19.6 Hz, 1H), 4.69 – 4.47 (m, 4H), 4.12 – 4.04 (m, 1H), 2.57 (s, 3H), 2.30 – 2.08 (m, 2H), 1.96 – 1.85 (m, 1H), 1.84 – 1.57 (m, 4H), 1.47 (t, *J* = 11.3 Hz, 1H), 1.36 (t, *J* = 7.1 Hz, 3H). MS (ES+) m/z 374.1 [M + H]^+^

### *Tert*-butyl (6-(2,2-difluorocycloheptyl)-4-methoxy-5-oxo-6,7-dihydro-5*H*-pyrrolo[3,4-*d*]pyrimidin-2-yl)carbamate (S42)

Reaction of methyl 2-((*tert*-butoxycarbonyl)amino)-4-formyl-6-methoxypyrimidine-5-carboxylate **(S37)** (420 mg, 1.35 mmol), 2,2-difluorocycloheptan-1-amine hydrochloride (250 mg, 1.35 mmol) and sodium triacetoxyborohydride (857 mg, 4.05 mmol) in DCM (4 mL) in an analogous manner to compound **S45,** with the addition of DIPEA (0.24 mL, 1.35 mmol) afforded **S42** as a white solid (430 mg, 77%). ^1^H NMR (500 MHz, DMSO*-d_6_*) *δ* 10.34 (s, 1H), 4.65 – 4.43 (m, 2H), 4.31 (d, *J* = 18.3 Hz, 1H), 4.02 (s, 3H), 2.28 – 2.04 (m, 2H), 1.95 – 1.85 (m, 1H), 1.84 – 1.77 (m, 1H), 1.77 – 1.67 (m, 2H), 1.67 – 1.42 (m, 14H). MS (ES+) m/z 413.5 [M + H]^+^

### 4-Chloro-6-cyclohexyl-2-methylsulfanyl-7H-pyrrolo[3,4-d]pyrimidin-5-one (S44)

Reaction of ethyl 4-chloro-6-formyl-2-(methylthio)pyrimidine-5-carboxylate (**S14)** (4460 mg, 17.10 mmol), cyclohexanamine (1.95 mL, 17.10 mmol) and sodium triacetoxyborohydride (10877 mg, 51.32 mmol) in DCM (160 mL)in an analogous manner to compound **S45** afforded **S14** as a white solid (3023 mg, 59%). ^1^H NMR (500 MHz, DMSO*-d_6_*) *δ* 4.49 (s, 2H), 3.99 (tt, *J* = 12.0, 3.8 Hz, 1H), 2.60 (d, *J* = 1.5 Hz, 3H), 1.76 (dd, *J* = 28.5, 12.7 Hz, 4H), 1.64 (d, *J* = 13.4 Hz, 1H), 1.52 (qd, *J* = 12.4, 3.4 Hz, 2H), 1.44 – 1.29 (m, 2H), 1.21 – 1.03 (m, 1H). Purity >90% by LCMS. No ionisation was obtained.

### 6-Cycloheptyl-4-methoxy-2-(methylthio)-6,7-dihydro-5*H*-pyrrolo[3,4-d]pyrimidin-5-one (S47)

Reaction of 4-chloro-6-cycloheptyl-2-(methylthio)-6,7-dihydro-5*H*-pyrrolo[3,4-*d*]pyrimidin-5-one **(S45)** (100 mg, 0.32 mmol) and MeOH (2 mL) in an analogous method to compound **S46** afforded **S47** as a white solid (72 mg, 77%). ¹H NMR (400 MHz, CDCl_3_) *δ* 4.45 - 4.37 (m, 1H), 4.27 (s, 2H), 4.15 (s, 3H), 2.62 (s, 3H), 1.77 - 1.54 (m, 12H). MS (ES+) *m/z* 308.3 [M + H]^+^

### 6-Cyclohexyl-4-(ethylamino)-2-(methylthio)-6,7-dihydro-5*H*-pyrrolo[3,4-*d*]pyrimidin-5-one (S50)

Reaction of 4-chloro-6-cyclohexyl-2-(methylthio)-6,7-dihydro-5*H*-pyrrolo[3,4-*d*]pyrimidin-5-one **(S44)** (120 mg, 0.40 mmol), THF (7 mL), ethylamine solution (30% in EtOH, 0.3 mL, 0.60 mmol) and DIPEA (0.14 mL, 0.80 mmol) in an analogous manner to compound **S51** where purification was achieved by flash chromatography (0 to 75% EtOAc in heptane) afforded **S47** as a white solid (100 mg, 77%). ^1^H NMR (400 MHz, DMSO*-d_6_*) *δ* 7.27 (t, *J* = 6.1 Hz, 1H), 4.26 (s, 2H), 3.89 (tt, *J* = 11.9, 3.9 Hz, 1H), 3.56 – 3.41 (m, 2H), 2.48 (s, 3H), 1.88 – 1.56 (m, 5H), 1.56 – 1.22 (m, 4H), 1.20 – 1.03 (m, 4H). MS (ES+) m/z 307.4 [M + H]^+^

### 6-Cycloheptyl-4-(methoxymethyl)-2-(methylthio)-6,7-dihydro-5*H*-pyrrolo[3,4-*d*]pyrimidin-5-one (S53)

To a solution of 4-bromo-6-cycloheptyl-2-(methylthio)-6,7-dihydro-5*H*-pyrrolo[3,4-*d*]pyrimidin-5-one **(S52)** (72 mg, 0.20 mmol) in DMF (5 mL), purged with nitrogen was added tributyl(methoxymethyl)stannane (101 mg, 0.30 mmol) and Pd(PPh_3_)_4_ (23 mg, 0.02 mmol). The reaction stirred at 120 ^°^C for 90 min, diluted with water and EtOAc, the combined organic layers washed with brine, dried over MgSO_4_, filtered through celite, concentrated *in vacuo* and purified by flash chromatography (0-60% EtOAc in heptane) to afford **S53** (16 mg, 25%) as a colourless oil. ^1^H NMR (500 MHz, CDCl_3_) *δ* 4.95 (s, 2H), 4.44 – 4.36 (m, 1H), 4.32 (s, 2H), 3.55 (s, 3H), 2.64 (s, 3H), 1.96 – 1.85 (m, 2H), 1.81 – 1.63 (m, 6H), 1.63 – 1.54 (m, 4H). MS (ES+) m/z 322.3 [M + H]^+^

### 6-Cyclohexyl-2-(methylthio)-5-oxo-6,7-dihydro-5*H*-pyrrolo[3,4-*d*]pyrimidine-4-carbonitrile (S55)

4-Chloro-6-cyclohexyl-2-(methylthio)-6,7-dihydro-5*H*-pyrrolo[3,4-*d*]pyrimidin-5-one **(S44)** (500 mg, 1.68 mmol) was suspended in HI (10 mL, 1.68 mmol) and all light was excluded. The reaction was stirred at rt for 16 h, neutralised with sat. NaHCO_3_ diluted with DCM, passed through a hydrophobic frit and concentrated *in vacuo* to afford 6-cyclohexyl-4-iodo-2-(methylthio)-6,7-dihydro-5*H*-pyrrolo[3,4-*d*]pyrimidin-5-one **(S54)**. MS (ES+) m/z 390.0 [M + H]^+^ To a solution of compound **S54** (530 mg, 1.36 mmol) in DMF (10 mL), purged with nitrogen, ZN(CN)_2_ (320 mg, 2.72 mmol) and Pd(PPh_3_)_4_ (157.5 mg, 0.13 mmol) were added. The reaction was stirred at 100 °C for 16 h, diluted with water and EtOAc, the combined organic layers were washed with brine, concentrated *in vacuo* and purified by flash chromatography (0 to 50% EtOAc in heptane) to afford compound **S55** (150 mg, 34% yield over two steps) as an off-white solid. ^1^H NMR (500 MHz, DMSO*-d_6_*) *δ* 4.58 (s, 2H), 4.02 (tt, *J* = 11.9, 3.8 Hz, 1H), 2.62 (s, 3H), 1.85 – 1.69 (m, 4H), 1.65 (d, *J* = 13.2 Hz, 1H), 1.55 (qd, *J* = 12.3, 3.4 Hz, 2H), 1.43 – 1.30 (m, 2H), 1.20 – 1.08 (m, 1H). MS (ES+) m/z 289.2 [M + H]^+^

### Ethyl 4-(2-(*tert*-butoxycarbonyl)-2-cycloheptylhydrazineyl)-6-ethoxy-2-(methylthio)pyrimidine-5-carboxylate (S58)

To a solution of ethyl 4-chloro-6-ethoxy-2-(methylthio)pyrimidine-5-carboxylate **(S56)** (12 g, 43.36 mmol) in NMP (100 mL) was added *tert*-butyl *N*-amino-*N*-cycloheptyl-carbamate (10.89 g, 47.70 mmol) and DIPEA (11 mL). The reaction was stirred at 120 °C for 6 h, poured into water (250 mL), extracted with EtOAc, the combined organic layers were dried over Na_2_SO_4_ and concentrated *in vacuo* to afford ethyl 4-(2-(*tert*-butoxycarbonyl)-2-cycloheptylhydrazineyl)-6-ethoxy-2-(methylthio)pyrimidine-5-carboxylate **(S57)**. MS (ES+) m/z 469.2 [M + H]^+^ **S57** (13 g, 27.74 mmol) was dissolved in MeOH (80 mL) and HCl (12 M, 21 mL) added dropwise at 0 °C. The reaction was stirred at 20 °C for 16 h, concentrated *in vacuo*, diluted with EtOAc and saturated NaHCO_3_, the combined organic layers were dried over Na_2_SO_4_ and concentrated *in vacuo* to afford compound **S58** (5.9 g, 37% yield over two steps) as a yellow oil. ^1^H NMR (400 MHz, CDCl_3_) *δ* 9.64 (br s, 1H), 4.52 - 4.35 (m, 2H), 4.28 (q, *J* = 7.2 Hz, 2H), 3.84 (s, 1H), 3.70 (s, 2H), 3.04 - 2.96 (m, 1H), 2.63 - 2.41 (m, 3H), 1.89 - 1.67 (m, 6H), 1.60 - 1.31 (m, 12H). MS (ES+) m/z 369.1 [M + H]^+^

### 2-Cycloheptyl-4-ethoxy-6-(methylthio)-1,2-dihydro-3*H*-pyrazolo[3,4-*d*]pyrimidin-3-one (S59)

To a solution of KOH (500 mg, 8.91 mmol) in water (1.5 mL) was added a solution of ethyl 4-(2-cycloheptylhydrazineyl)-6-ethoxy-2-(methylthio)pyrimidine-5-carboxylate **(S58)** (500 mg, 1.36 mol) in EtOH (5 mL). The reaction was stirred at 20 °C for 1 h, poured into water, extracted with EtOAc, the combined organic layers were dried over Na_2_SO_4_, concentrated *in vacuo* and purified by flash chromatography (10 – 50% EtOAc in petroleum ether) to afford compound **S59** (145 mg, 32%) as a light yellow solid. ^1^H NMR (400 MHz, DMSO*-d_6_*) *δ* 12.02 - 11.81 (br, 1H), 4.47 (q, *J* = 7.0 Hz, 2H), 4.40 - 4.22 (m, 1H), 2.52 - 2.47 (s, 3H), 1.91-1.64 (m, 6H), 1.64 - 1.40 (m, 6H), 1.38 - 1.28 (m, 3H). MS (ES+) m/z 323.1 [M + H]^+^

### LysRS protein expression and crystallisation

Protein expression and purification was carried out as previously described^14^. LysRS was crystallised using vapour diffusion in hanging drop plates, with reservoir solutions containing 0.2 – 0.3 M NaOAc and 14-18% w/v PEG33350. Protein, at 20 mg/ml in 0.1 M HEPES, 0.15 M NaCl, 5 % v/v glycerol pH 7.5, was incubated with 5 mM lysine for one hour prior to setting up crystallisation drops consisting of 1 µL reservoir solution and 1 µL of protein. Ligands were dissolved in DMSO to a concentration of 200 mM; these were then diluted to 10 mM using protein storage buffer. For soaking, crystals were transferred to drops consisting of 1 µL reservoir solution and 1 µL of the 10 mM ligand stock. Crystals were soaked at room temperature for one hour, harvested and cryoprotected using reservoir solution supplemented with 33% glycerol, then flash frozen.

### Data collection and refinement

Data for **32** & **42** were collected in-house using a Rigaku Micromax -007 HF diffractometer. Data for **25** & **10** were collected at beamline ID23-1 at the European Synchrotron Radiation Facility (ESRF) and data for **27** were collected at beamline I04-1 at Diamond Light Source. Ligand dictionaries were prepared using AceDRG^20^, incorporated into the CCP4^21^ suite of software. Ligands were manually placed using Coot^22^ and structures refined using Refmac^23^. Full refinement statistics are given in **Table S1**.

### LysRS *in vitro* enzyme assay

LysRS assays were run in 50 µL reactions (Final concentration: 30 mM Tris-HCl 8.0, 40 mM MgCl2, 140 mM NaCl, 30 mM KCl, 0.01% Brij-35, 1 mM DTT, 3 µM ATP, 12 µM lysine, 200 nM LysRS and 0.5 U/mL Pyrophosphatase) at room temperature for 8 h. Reactions were started by adding enzyme into wells containing substrate and compounds. Reactions were stopped by the addition of Biomol green (50 µL: Enzo Life Sciences) with the amount of free phosphate detected by measuring absorbance (650 nm) after 20 min further incubation (BMG Pherastar plate reader). For the most potent compounds (in **Table 6**) the tight binding limit of the assay was reduced by the following modification to the original assay: 40 nM LysRS; 30 µM ATP; 20 h incubation. In both assay formats 100% inhibition control reactions were performed in the absence of lysine as substrate. Samples were all run in duplicate, data was processed and analysed using ActivityBase (IDBS).

### Mode of Inhibition studies

Assays were run in the buffer previously described in the presence of 50 µM ATP and 60 nM LysS; conditions that had a good assay signal and allowed determination of IC_50_ values not restricted by the tight binding limit. IC_50_ shift experiments were performed in the presence of variable lysine concentrations (0.5, 1, 2, 6 and 12 µM). Data analysis of dose-dependent curves was performed using SigmaPlot, and the Mode of Inhibition was informed by plotting IC_50_ values versus [S]/*K_M_*^18^.

### Construction of and MIC measurement with Mtb *lysS*-TetON

Mtb *lysS*-TetON was constructed using the same strategy used to generate similar mutants for other essential genes, such as *trxB2* and *mmpL3*^24–25^. Briefly, wild type H37Rv was transformed with a constitutive *lysS* expression plasmid that integrated into the attachment site (attL5) of the bacteriophage L5 and then deleted the native copy of lysS by homologous recombination. Next, the constitutive lysS expression plasmid was replaced with a regulated expression plasmid in which transcription of lysS is repressed by a tetracycline repressor (TetR) so that expression of LysRS is induced with anhydrotetracycline (Atc) and reduced (but not repressed completely) without it. For MIC measurements, WT H37Rv and Mtb *lysS*-TetON were grown in Middlebrook 7H9 (with hygromycin at 50 μg/mL, kanamycin at 25 μg/mL, and, for *lysS*-TetON, ATc at 500 ng/mL) supplemented with 0.2% (v/v) glycerol, 0.05% (v/v) Tyloxapol, and ADNaCl (0.5% [w/v] BSA, 0.2% [w/v] dextrose, and 0.85% [w/v] NaCl). After cultivation for 7 days at 37 °C and 5% CO_2_ in a humidified incubator, cultures were washed with 7H9, diluted to an OD_580_ of ∼0.05 and grown at 37 °C with 5% CO_2_ for 5 days in 7H9, with and without Atc (500 ng/mL). Compounds were solubilized in DMSO and dispensed into black, clear-bottom 384-well tissue culture plates using an HP D300e Digital Dispenser as 16-point, 2-fold dilution series in triplicate. OD_580_ 0.01 suspension (50 μL) was pipetted to each well, and cultures were incubated for 7–14 days at 37 °C under the same conditions as above. Final OD_580_ values were normalized to no-drug (1% [v/v] DMSO) control wells.

### Additional assays

All other assays including KARS1, MIC, HepG2, *in vitro* ADME, *in vivo* PK analysis and *in vivo* efficacy were performed as described previously^14^. For biological *in vitro* assays all samples were run at least in duplicate and the average data is shown.

### Ethical Statements

#### Mouse Pharmacokinetics

All regulated procedures, at the University of Dundee, on living animals was carried out under the authority of a project licence issued by the Home Office under the Animals (Scientific Procedures) Act 1986, as amended in 2012 (and in compliance with EU Directive EU/2010/63). Licence applications will have been approved by the University’s Ethical Review Committee (ERC) before submission to the Home Office. The ERC has a general remit to develop and oversee policy on all aspects of the use of animals on university premises and is a subcommittee of the University Court, its highest governing body.

#### Acute efficacy studies

All procedures were performed in accordance with protocols approved by the GSK Institutional Animal Care and Use Committee and met or exceeded the standards of the American Association for the Accreditation of Laboratory Animal Care (AAALAC). All animal studies were ethically reviewed and carried out in accordance with European Directive 2010/63/EEC and the GSK Policy on the Care, Welfare and Treatment of Animals.

## ANCILLARY INFORMATION SUPPORTING INFORMATION

X-ray crystallographic data collection and refinement statistics; Pharmacokinetic data collected throughout the project; *In vivo* exposure for the four most advanced compounds. *In vivo* metabolite identification for **11**; Additional co-crystal structures; Surface view of **49** in the ATP binding pocket of LysRS; 1H NMR spectra for compounds in manuscript; HRMS for compounds progressed to in vivo studies; Supplementary methods Key *in vitro* data table with PAINS alert analysis and SMILES

## PDB ID CODES

PDB codes for LysRS with bound compounds are: **10** (9qea), **25** (9qei), **27** (9qbr), **32** (9qdj), **37** (9qc3) and **42** (9qc4).

## Supporting information

Data and PAINS summary

## AUTHOR CONTRIBUTIONS

SD, MM, KB, AS, MC, RB and LC contributed to medicinal chemistry design, planning, and synthesis; AD, FT, JP, CE, JK, PP, DS ELR and SG planned/executed/analysed biological/crystallographic studies; SD, CJ, MK, FZ, and LC contributed to computational chemistry design and modelling; GW scale up synthesis and route optimisation; BB and PR for initial discussions; PS, OE and KDR planned/executed/analysed DMPK/safety studies; LGL design and interpretation of *in vivo* efficacy studies; LE, RB, PW, SG and LC contributed to monthly project planning meetings; SD, SG and LC wrote and edited the manuscript. All authors reviewed and approved the final version of this manuscript for publication.

**Notes:** ELR, LGL, KR, LE, RB and PW are employees and/or shareholders in GlaxoSmithKline. The other authors declare no competing interests.

## ACKNOWLEDGMENTS

We thank Jennifer Riley Nicole Mutter, Yoko Shishikura, Maria Osuna-Cabello, Laura Frame, Erika Pinto, Laste Stojanovski, Fred Simeons, Liam Ferguson, Lorna Campbell, Alex Cookson, Kirsty Cookson, Desiree Zeller, and Kieran Cartmill, for technical assistance. Darren Edwards for comments on the MetID study and Neil Norcross for initial synthetic chemistry discussions. We thank Beatriz Rodriguez-Miquel for her support with data analysis. GSK acknowledges the University of Zaragoza for the provision of H37Rv. This work was funded in part by an award to P.G.W. from The Gates Foundation (GF) (OPP1066891 and OPP1191579) and Wellcome (100195/Z/12/Z); awards to D.S. from GF (INV-010616 and INV-004761); awards to K.Y.R. from GF (INV-004709 and OPP 1177930). The conclusions and opinions expressed in this work are those of the author(s) alone and shall not be attributed to the Foundation. Under the grant conditions of the Foundation, a Creative Commons Attribution 4.0 License has already been assigned to the Author Accepted Manuscript version that might arise from this submission. Please note works submitted as a preprint have not undergone a peer review process. The authors would like to thank Diamond Light Source (proposal mx19844; beamline I04-1) and the European Synchrotron Radiation Facility (proposal mx1982; beamline ID23-1) for beamtime, and the staff at the beamlines for assistance with crystal testing and data collection. We would also like to acknowledge the X-ray Crystallography Facility at the University of Dundee, which was supported by Wellcome (award no. 094090).

## ABBREVIATIONS

ATc: anhydrotetracycline
DIPEA: diisopropylethylamine
HEPES: 4-(2-hydroxyethyl)piperazine-1-ethanesulfonic acid
KARS1: *H. sapiens* lysyl
tRNA: synthetase
LysRS: *M. tuberculosis* lysyl tRNA synthetase
TEA: triethylamine

## For Table of Contents Use Only

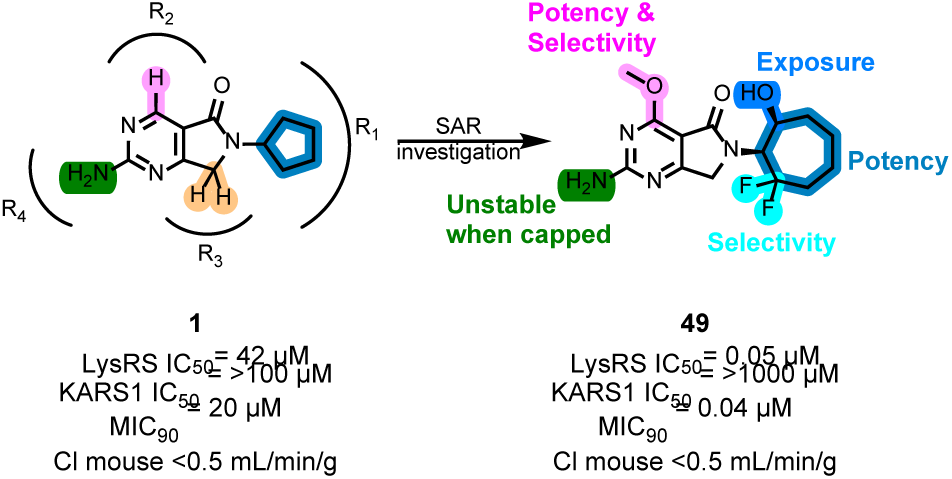

