## Supplemental info for "Design and development of lysyl tRNA synthetase inhibitors, for the treatment of tuberculosis"

**ORCHID iD** 0000-0001-6218-0092


| **Table of Contents** | Page |
|---|---|
| X-ray crystallographic data collection and refinement statistics | S2 |
| Pharmacokinetic data collected throughout the project | S4 |
| *In vivo* exposure for the four most advanced compounds | S4 |
| *In vivo* metabolite identification for **11** | S5 |
| Additional co-crystal structures | S6 |
| Surface view of **49** in the ATP binding pocket of LysRS | S6 |
| 1H NMR spectra for compounds in manuscript | S7 |
| HRMS for compounds progressed to in vivo studies | S27 |
| Supplementary Methods | S32 |

**Table S1: Crystallography data collection and refinement statistics.**

| | **10** | **25** | **27** |
|---|---|---|---|
| **Data collection** | | | |
| Source<br>Wavelength | ESRF ID23-1<br>0.97625 | ESRF ID23-1<br>0.97625 | Diamond I04-1<br>0.91587 |
| Space group | $P4_12_12$ | $P4_12_12$ | $P4_12_12$ |
| Cell dimensions | | | |
| *a, c* (Å) | 85.00, 148.04 | 83.93, 147.29 | 83.58, 147.41 |
| Resolution (Å) | 2.3 (2.38-2.30) | 2.4 (2.49-2.40)* | 2.4 (2.49-2.40) |
| $R_{merge}$ | 0.117 (1.218) | 0.113 (2.311) | 0.254 (5.109) |
| $I / \sigma I$ | 10.7 (2.0) | 8.5 (0.6) | 6.5 (0.7) |
| CC1/2 | 0.999 0.960) | 0.999 (0.508) | 0.997 (0.597) |
| Completeness (%) | 92.1 (99.9)** | 96.4 (99.0)** | 100 (100) |
| Redundancy | 10.0 (11.2) | 7.6 (7.7) | 14.4 (14.1) |
| **Refinement** | | | |
| Resolution (Å) | 2.30 | 2.40 | 2.40 |
| No. reflections | 21736 | 19448 | 20063 |
| $R_{work}$ / $R_{free}$ | 25.5 / 31.1 | 25.0 / 32.0 | 23.2 / 30.8 |
| No. atoms | | | |
| Protein | 3646 | 3624 | 3595 |
| Ligand / lysine | 20 / 10 | 21 / 10 | 20 / 10 |
| Water | 11 | 26 | 58 |
| *B*-factors | | | |
| Protein | 75.8 | 91.7 | 80.8 |
| Ligand / Lysine | 54.2 / 64.4 | 69.0 / 82.0 | 60.0 / 64.0 |
| Water | 52.0 | 61.3 | 58.7 |
| R.m.s. deviations | | | |
| Bond lengths (Å) | 0.0063 | 0.0115 | 0.0053 |
| Bond angles (°) | 1.4472 | 2.0607 | 1.5189 |
| PDB codes | 9qea | 9qei | 9qbr |

*Values in parentheses are for highest-resolution shell.

** completeness is lower due to significant ice ring

**Table S1 (cont.): Crystallography data collection and refinement statistics.**

| | 32 | 37 | 42 |
|---|---|---|---|
| **Data collection** | | | |
| Source<br>Wavelength | Inhouse<br>1.54178 | Diamond I04-1<br>0.91188 | Inhouse<br>1.54178 |
| Space group | $P4_12_12$ | $P4_12_12$ | $P4_12_12$ |
| Cell dimensions | | | |
| *a*, *c* (Å) | 83.70, 147.69 | 83.67, 147.26 | 83.98, 147.32 |
| Resolution (Å) | 2.6 (2.71-2.60) | 2.28 (2.32-2.28) | 2.6 (2.72-2.60)* |
| $R_{merge}$ | 0.360 (2.404) | 0.142 (1.702) | 0.158 (0.522) |
| $I / \sigma I$ | 4.8 (0.8) | 11.1 (0.9) | 8.5 (2.3) |
| CC1/2 | 0.988 (0.589) | 0.991(0.894) | 0.988 (0.972) |
| Completeness (%) | 99.7 (98.1) | 100 (1000) | 99.7 (97.5) |
| Redundancy | 11.2 (9.6) | 12.7 (12.8) | 13.8 (11.4) |
| **Refinement** | | | |
| Resolution (Å) | 2.60 | 2.28 | 2.60 |
| No. reflections | 15953 | 23355 | 15966 |
| $R_{work}$ / $R_{free}$ | 26.0 / 31.8 | 21.4 / 27.7 | 23.2 / 29.2 |
| No. atoms | | | |
| Protein | 3654 | 3591 | 3595 |
| Ligand / lysine | 21 / 10 | 21 / 10 | 27 / 10 |
| Water | 12 | 82 | 31 |
| *B*-factors | | | |
| Protein | 65.7 | 69.8 | 56.3 |
| Ligand / Lysine | 42.1 / 46.6 | 52.2 / 48.7 | 43.3 / 40.7 |
| Water | 37.2 | 52.2 | 34.1 |
| R.m.s. deviations | | | |
| Bond lengths (Å) | 0.0044 | 0.0082 | 0.0036 |
| Bond angles (°) | 1.2122 | 1.5664 | 1.0064 |
| PDB codes | 9qdj | 9qc3 | 9qc4 |

*Values in parentheses are for highest-resolution shell.

**Table S2: Pharmacokinetic data collected throughout the study.**

| | 8 | 10 | 11 | 25 | 34 | 37 | 49 |
|---|---|---|---|---|---|---|---|
| $C_{max}$ (ng/mL) | 1,400 | 5,300 | 4,800 | 4,260 | 339 | 2,670 | 3700 |
| AUC (µg/mL.min) | 110 | 220 | 270 | 400 | 68 | 450 | 500 |
| Cl (mL/min/kg) | 28 | 19 | 16 | 23 | 29 | 22 | 15 |
| $T_{1/2}$ (h) | 0.3 | 0.3 | 0.3 | 0.4 | 1 | 2 | 1.4 |
| $V_d$ (L/kg) | 0.7 | 0.4 | 0.4 | 0.7 | 4 | 2.2 | 1.0 |
| F (%) | 31 | 43 | 43 | 91 | 16 | 100 | 74 |

Comparative murine pharmacokinetic parameters obtained across the SAR program

**Table S3: *In vivo* exposure for four most advanced compounds.**

| | 46 | 47 | 48 | 49 |
|---|---|---|---|---|
| $C_{max}$ (ng/mL) | 118,000 | 25,300 | 50,500 | 35,100 |
| AUC (µg/mL.min) | 169 | 36 | 100 | 181 |

Total blood concentrations taken at four time-points on first day of dosing from individual infected mice.

A

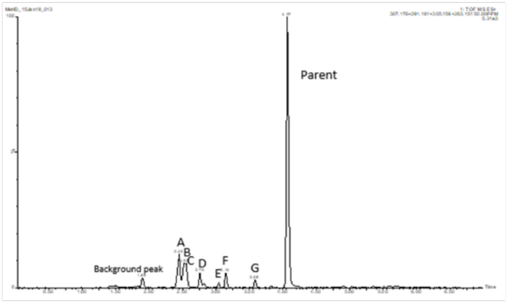


B

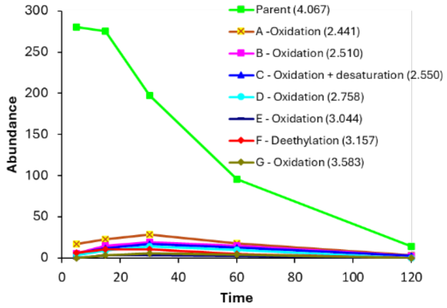


C

| Peak ID | Tentative Met. Id. | Proposed [M+H]+ ([M-H]-) | Ret. Time - min acidic (basic) | % Peak Area | Product Ions - positive (negative) |
|---|---|---|---|---|---|
| Parent (P) | Parent | 291 | 4.07 | 67 | 55,97,98,125,150,167,221,263,291 |
| A | P + O | 307 | 2.44 | 10 | 67,95,125,150,167,195 |
| B | P + O | 307 | 2.51 | 7 | 67,95,125,150,167,195,261,279 |
| C | P + O - 2H | 305 | 2.55 | 6 | 93,111,125,167,195,200 |
| D | P + O | 307 | 2.76 | 5 | 95,125,150,167,195,200,000 |
| E | P + O | 307 | 3.04 | 1 | No data |
| F | P - C2H4 | 263 | 3.16 | 4 | 125,167,263 |
| G | P + O | 307 | 3.58 | 2 | 125,167 |

D

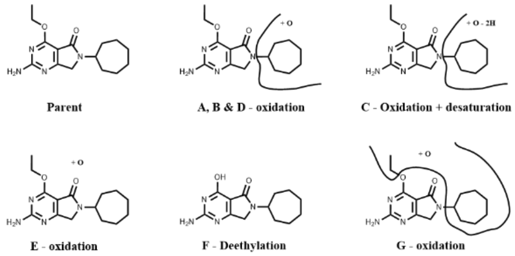


### Figure S1: *In vivo* metabolite identification for 11.

**A.** Overlaid extracted ion chromatogram (EIC) of parent compound and identified metabolites (A to G). **B.** Change in ion abundance with time for identified mass spectrometer signals for parent compound and identified metabolites. **C.** Summary table showing parent compound and identified metabolites with their precursor ion masses, retention time, % peak area (% of total peak area of the proposed [M+H]+ ions at 30 min time point) and MS2 fragment ions. **D.** Proposed location of identified metabolic transformations based upon the observed MS2 fragment ions.

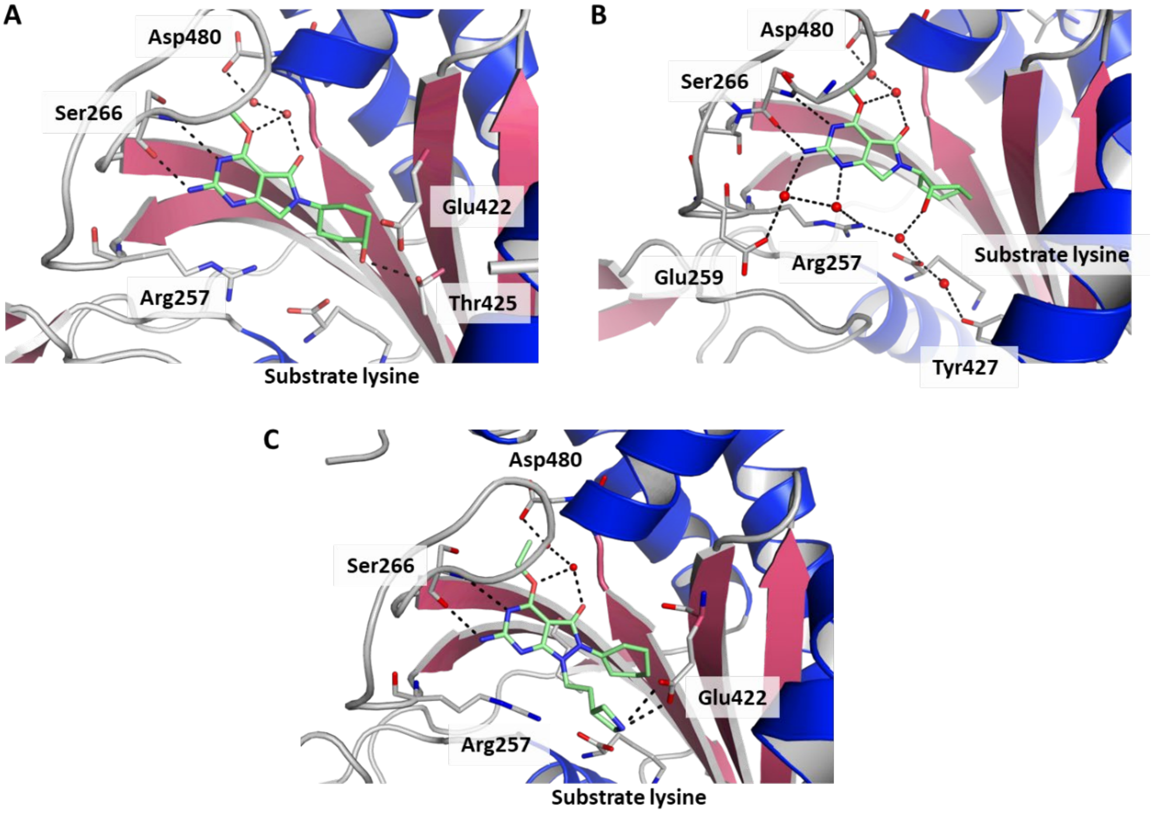


### Figure S2: Additional co-crystal structures.

The ligands are shown with mint green carbon atoms, blue nitrogen atoms and red oxygen atoms. Key water molecules are shown as red spheres and hydrogen bonds by dashed lines. Key residues involved with binding are shown. **A. 27** (PDB 9qbr); **B. 37** (PDB 9qc3) and **C. 42** (PDB 9qc4). Figures prepared using PyMol: The PyMOL Molecular Graphics System, Version 2.5.5 Schrödinger, LLC.

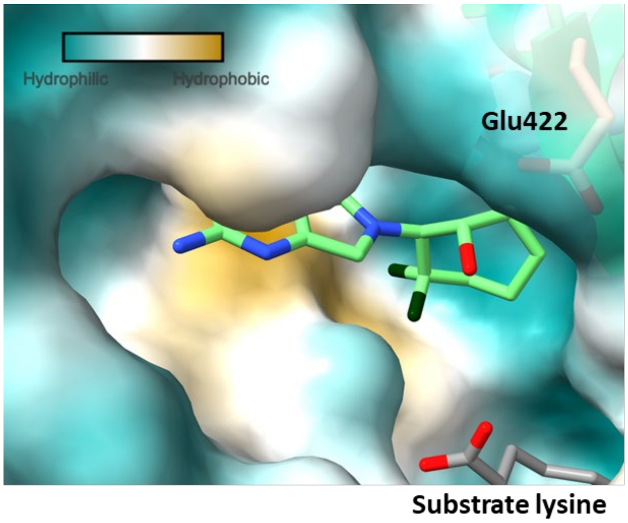


### Figure S3: Surface view of 49 in the ATP binding pocket of LysRS.

Figure prepared as Figure 3 in main article. The polar hydroxyl group of **49** sits in the more hydrophilic part of the pocket pointing towards Glu422. While the hydrophobic di-fluoro group sits in the hydrophobic area formed by the side chain of Met271 (yellow surface).

**1H NMR spectra for compounds in manuscript.**

**2**

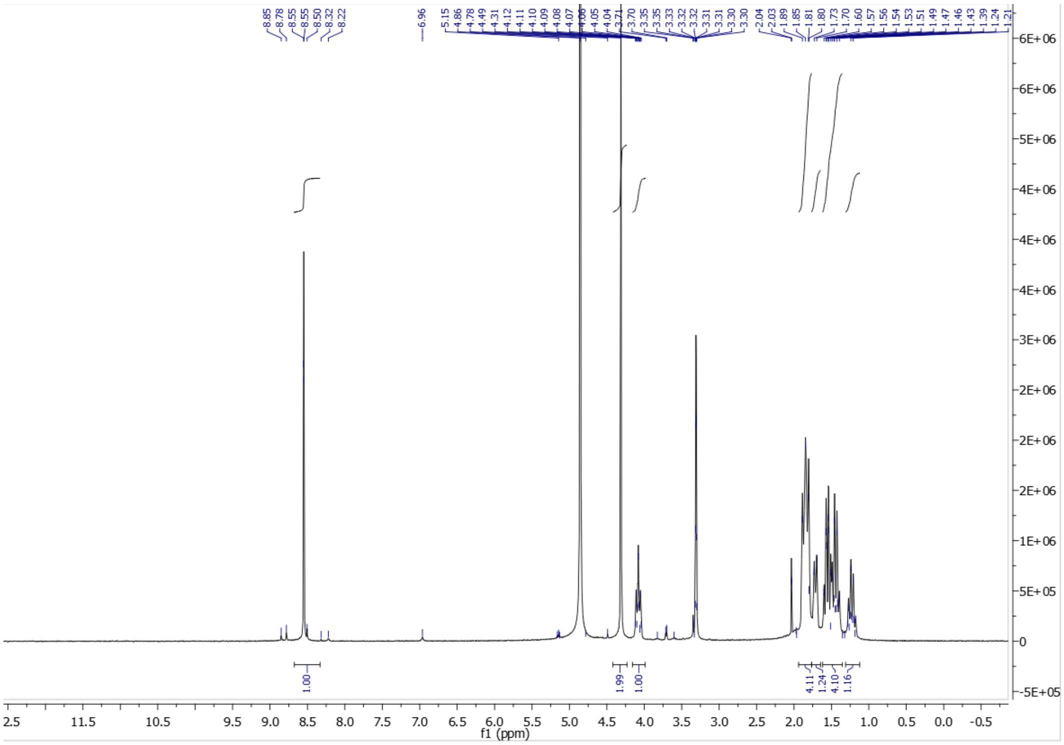

**3**

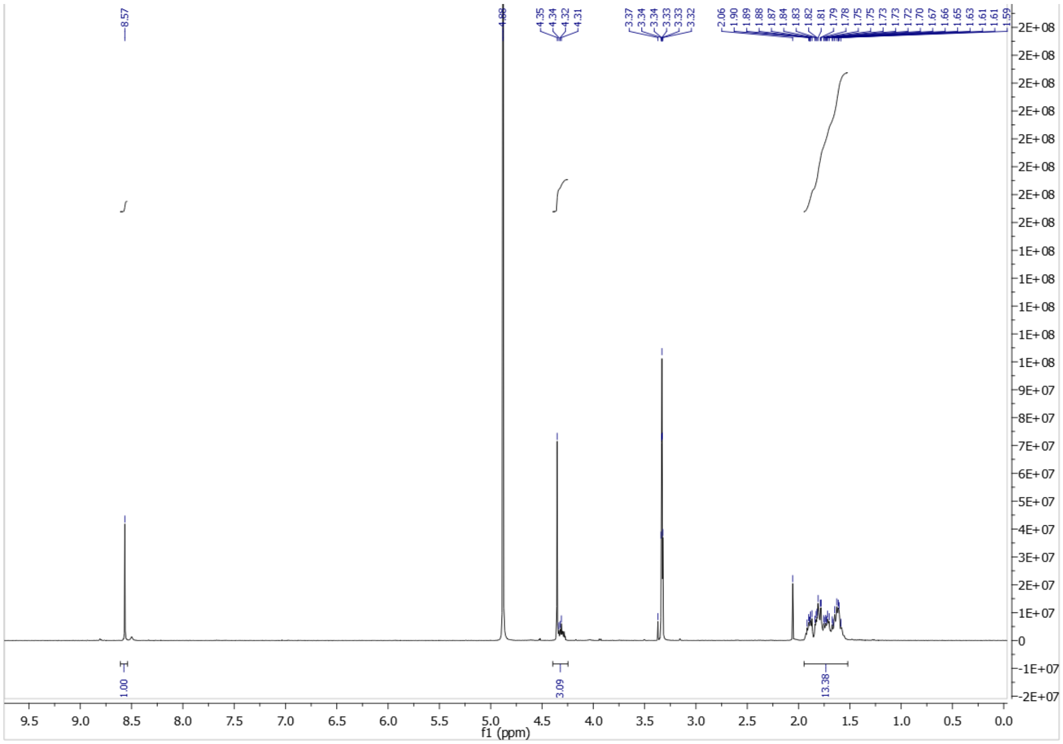

**4**

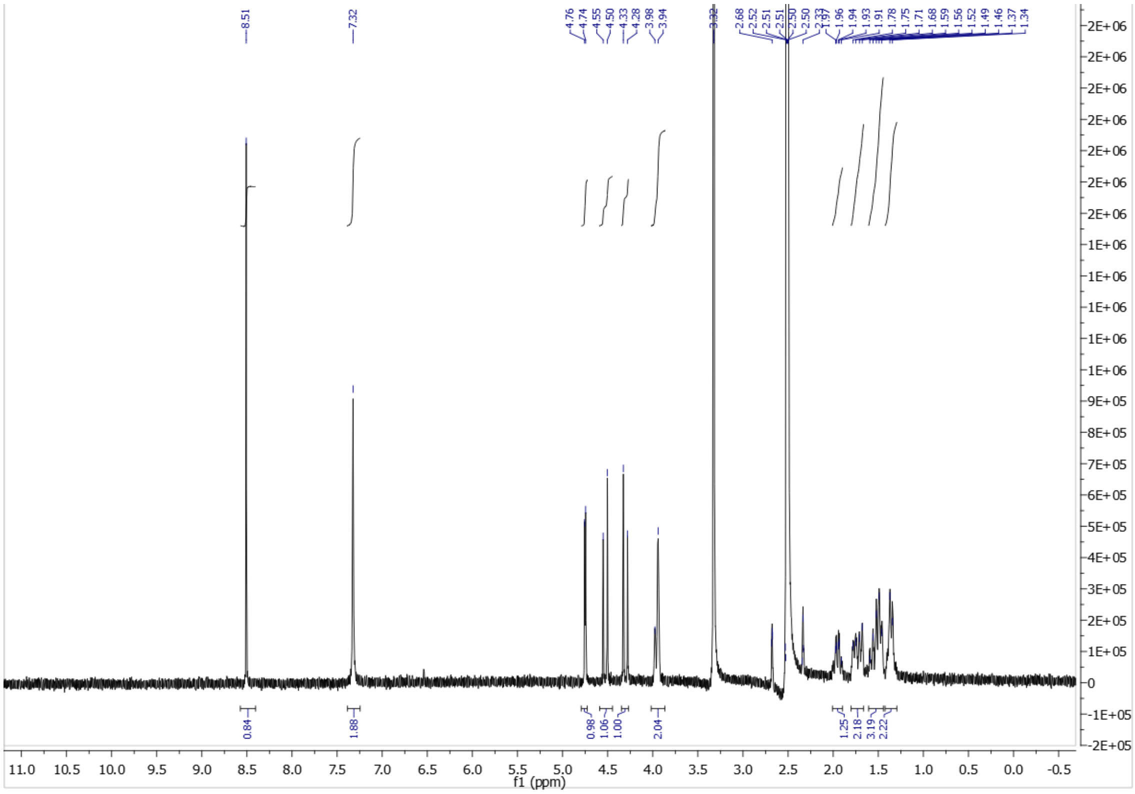

**5**

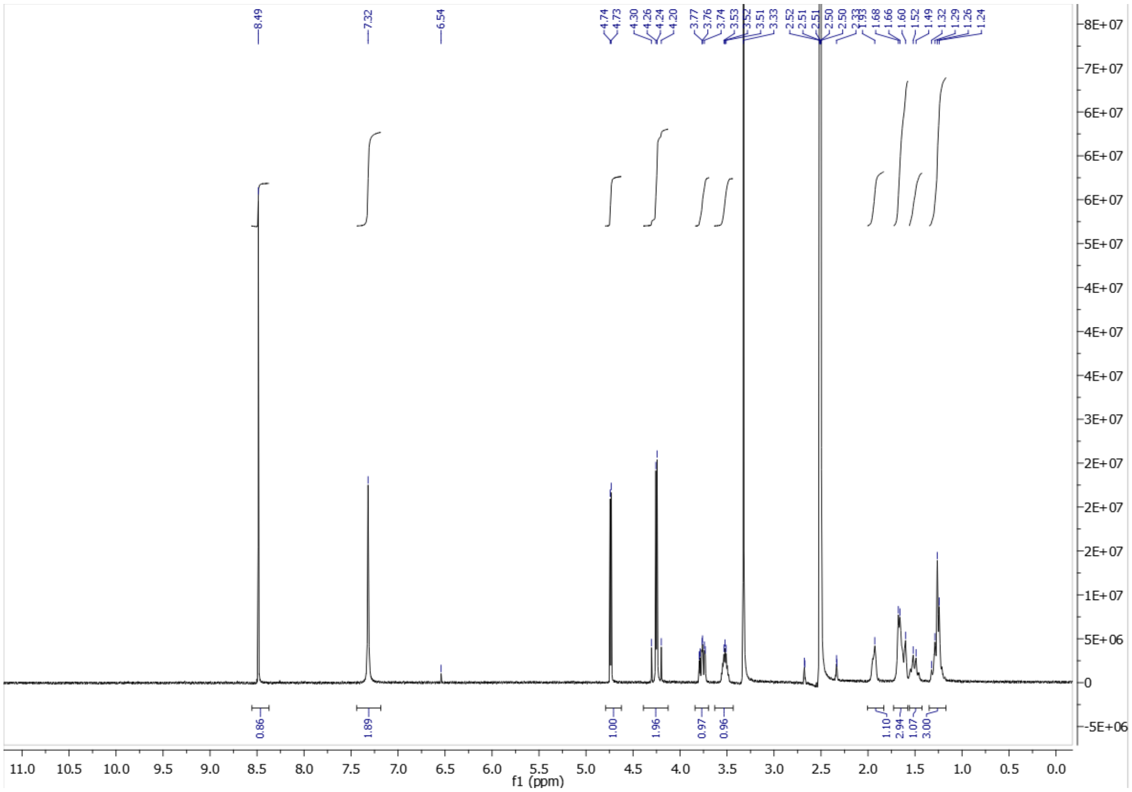

**6**

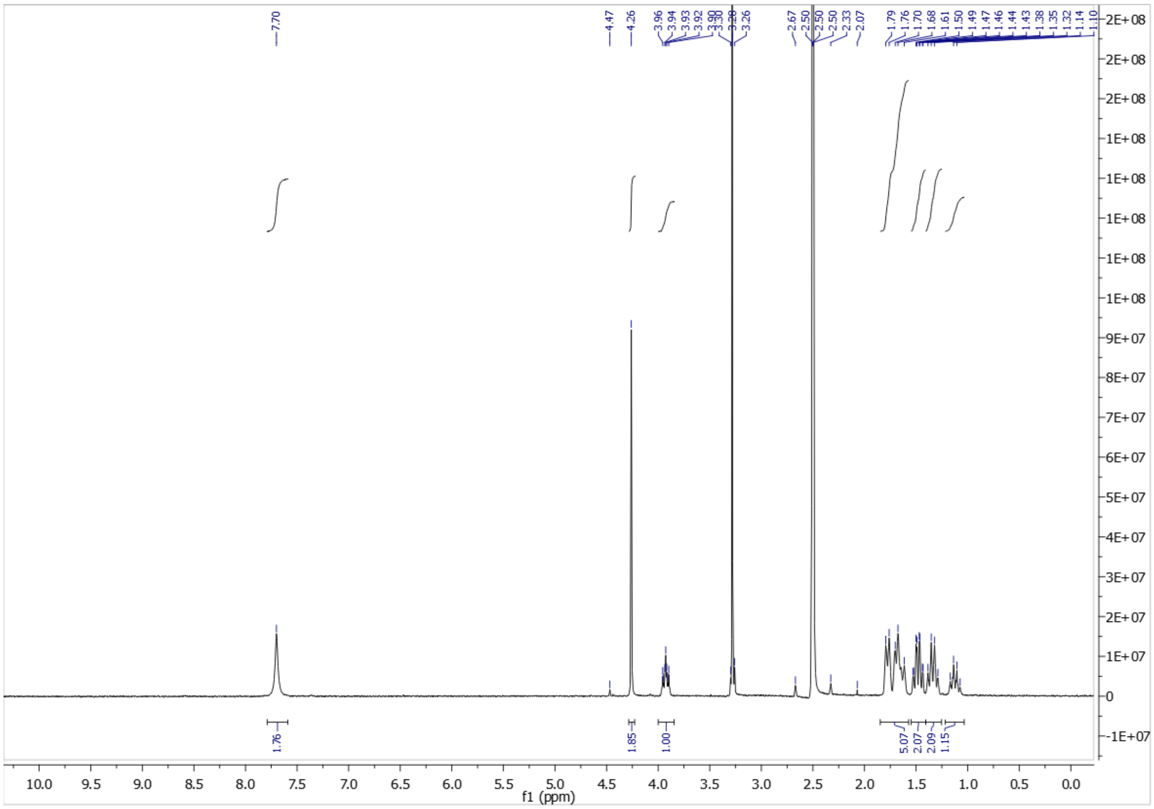

**7**

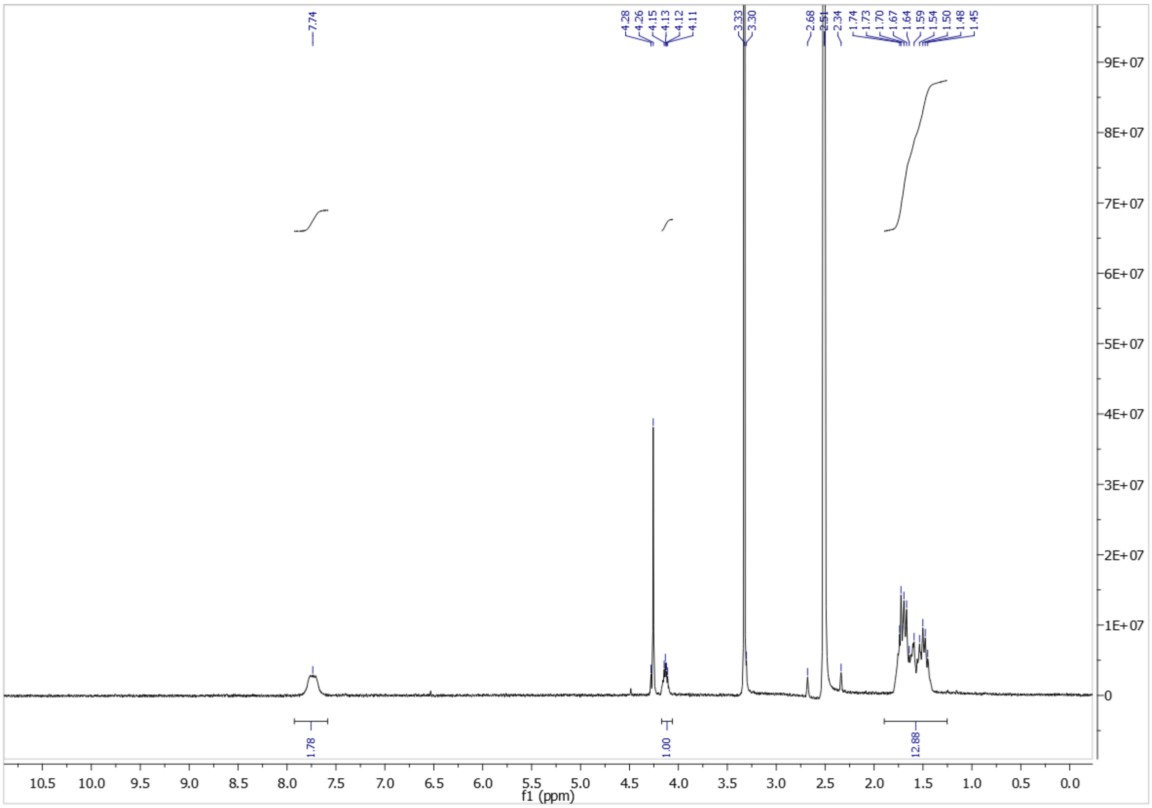

8

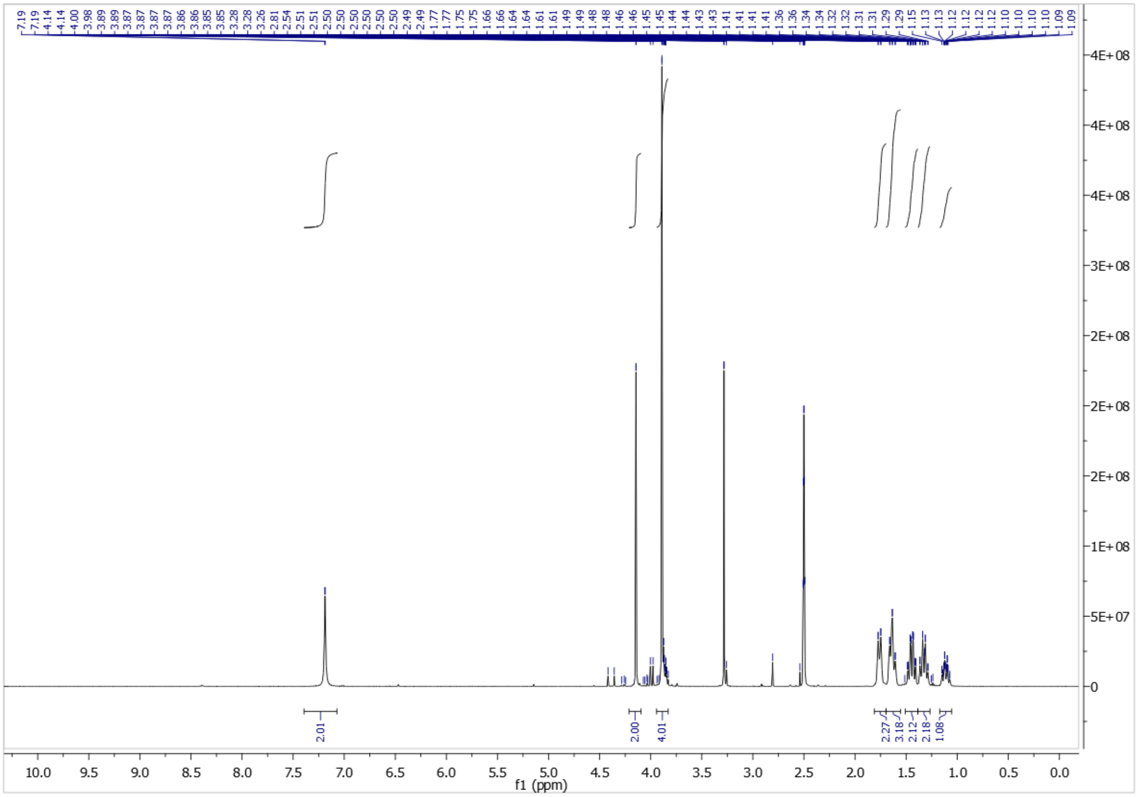

9

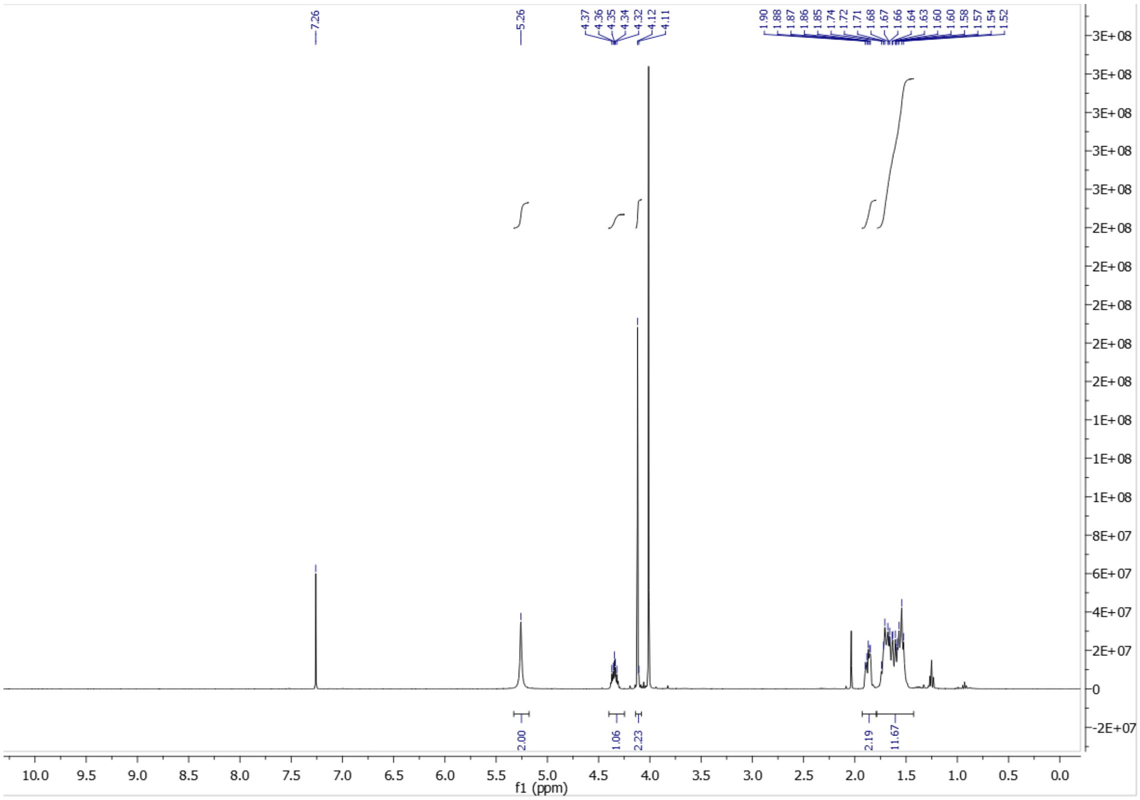

**12**

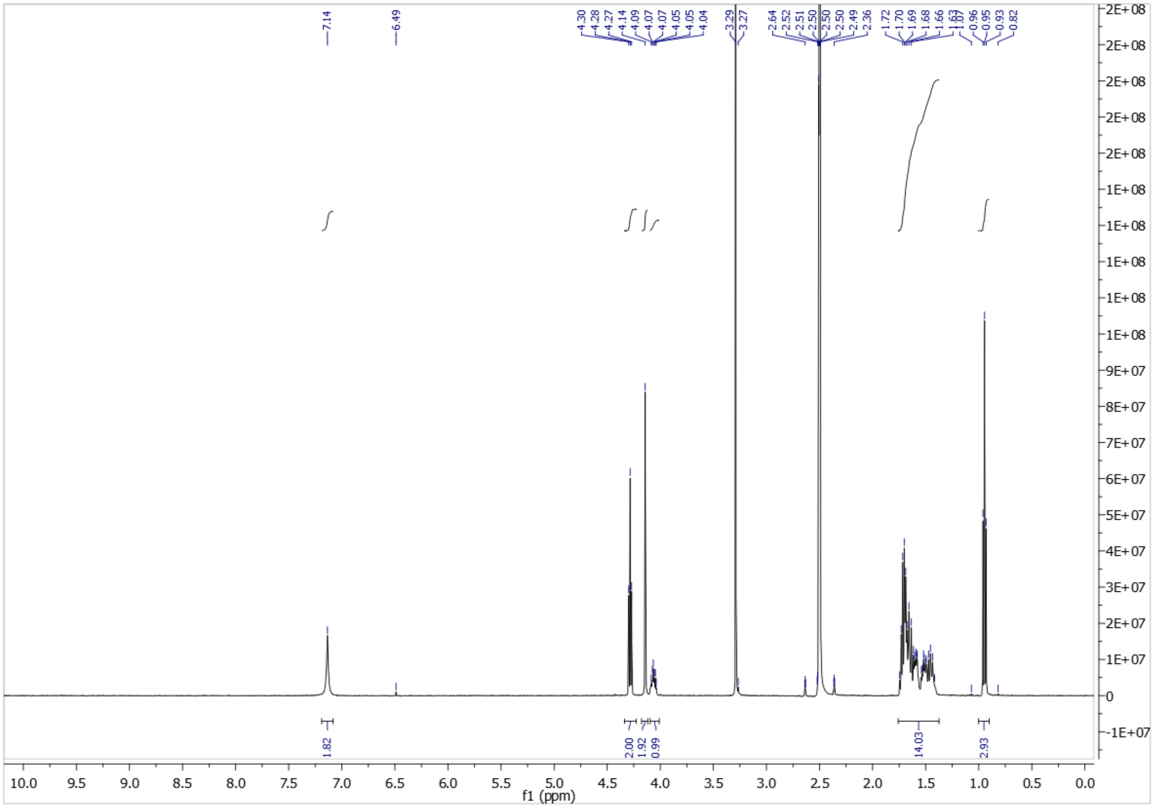

**13**

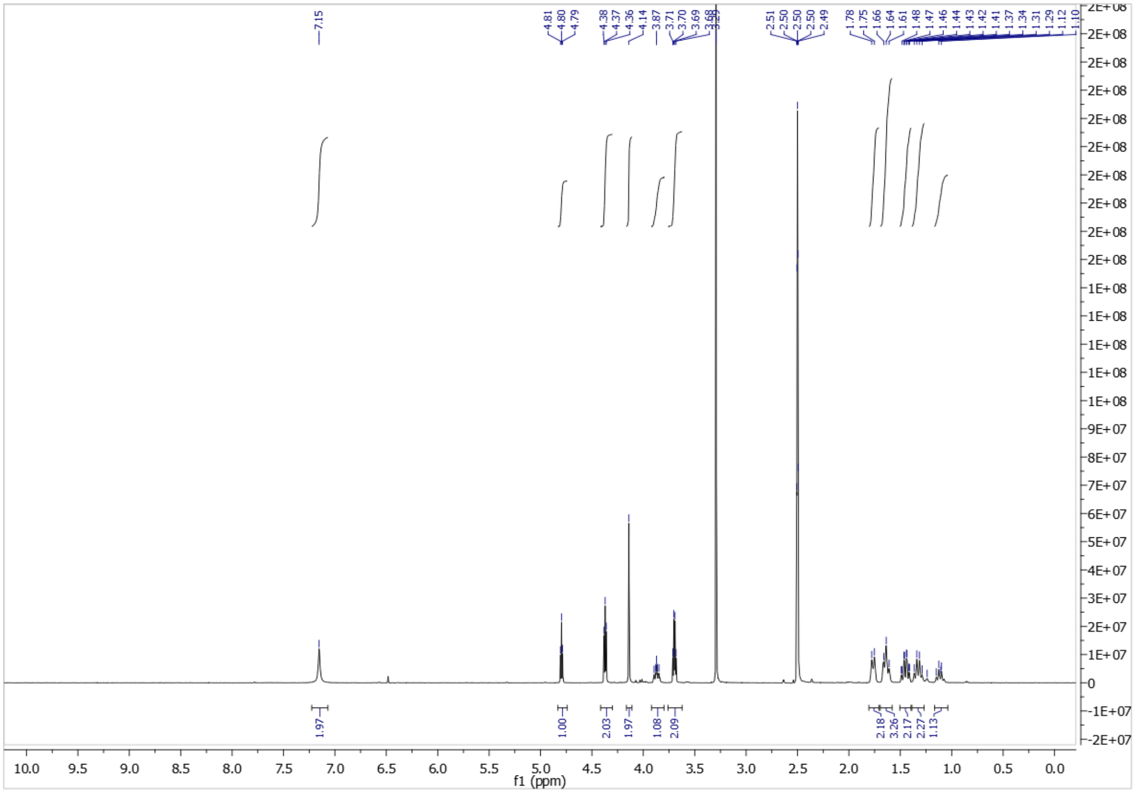

**14**

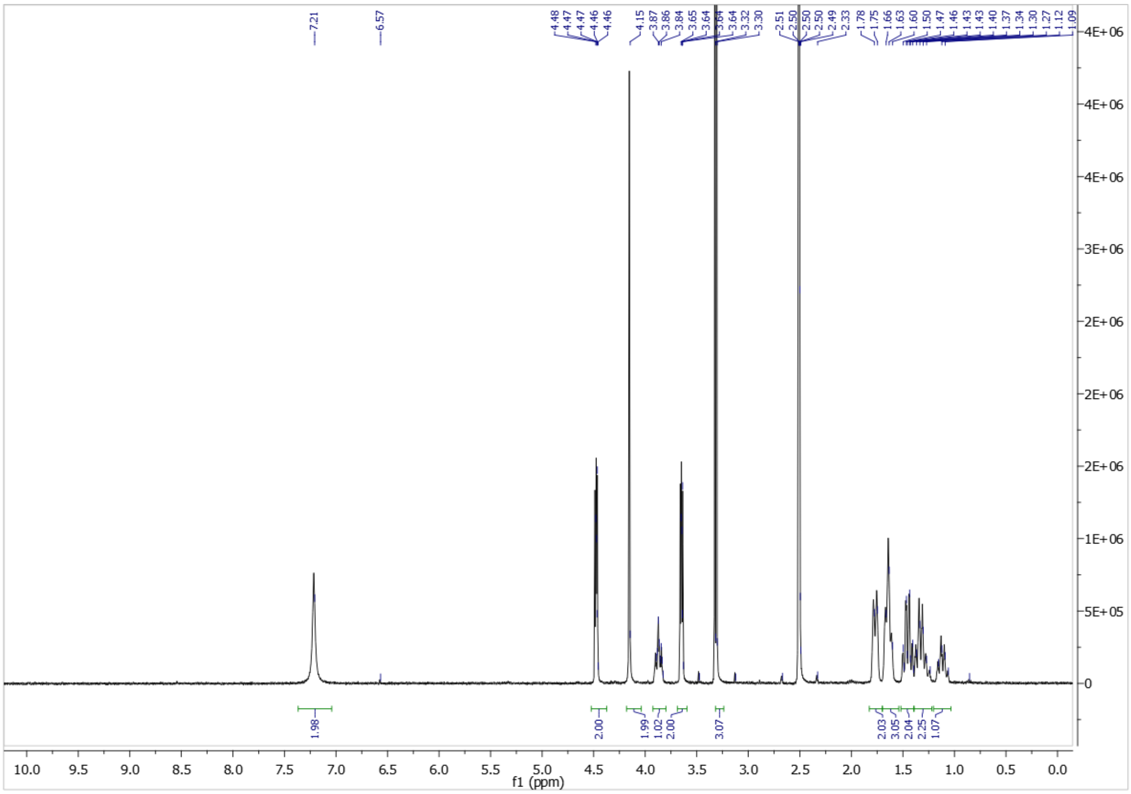
7.21
6.57
4.48
4.47
4.47
4.46
4.46
4.15
3.87
3.86
3.84
3.65
3.64
3.64
3.64
3.32
3.30
2.51
2.50
2.50
2.49
2.33
1.78
1.75
1.66
1.63
1.60
1.50
1.47
1.46
1.43
1.43
1.40
1.37
1.34
1.30
1.27
1.12
1.09
1.98
2.00
1.99
1.02
2.00
3.07
2.03
3.05
2.04
2.25
1.07
4E+06
3E+06
2E+06
1E+06
5E+05
0
10.0 9.5 9.0 8.5 8.0 7.5 7.0 6.5 6.0 5.5 5.0 4.5 4.0 3.5 3.0 2.5 2.0 1.5 1.0 0.5 0.0
f1 (ppm)

**15**

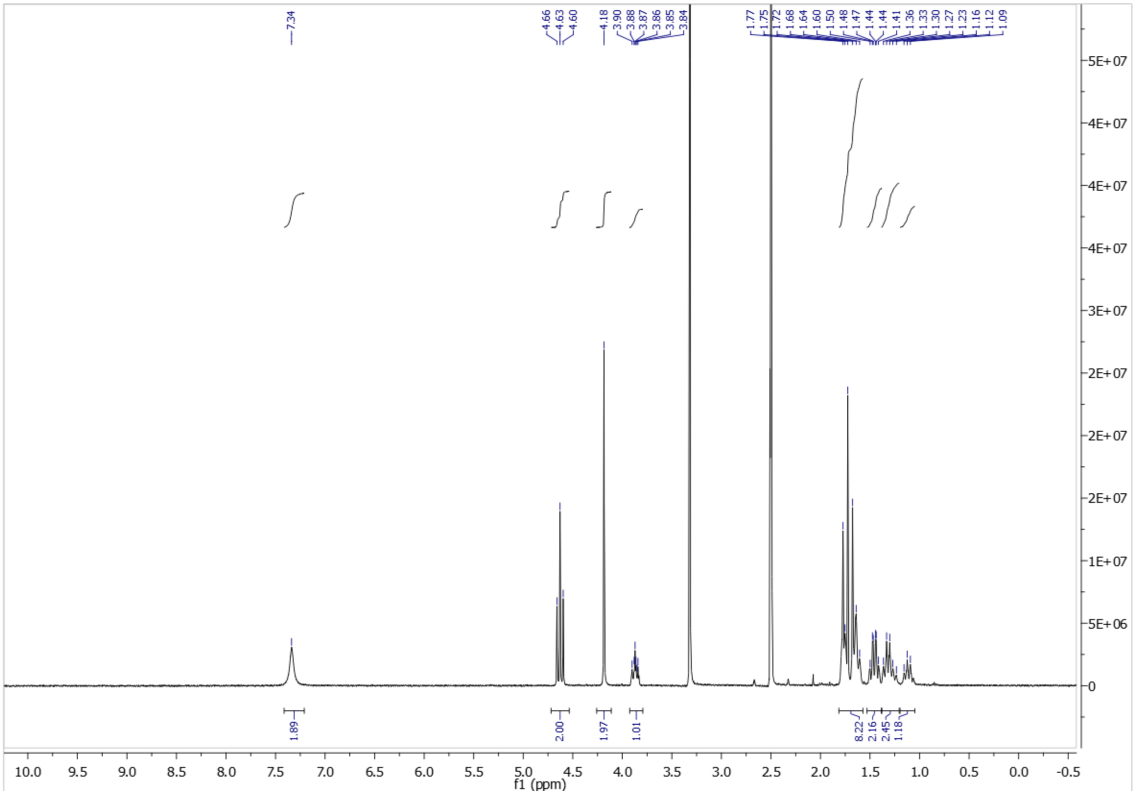
7.34
4.66
4.63
4.60
4.18
3.90
3.88
3.87
3.86
3.85
3.84
1.77
1.75
1.72
1.68
1.64
1.60
1.50
1.48
1.47
1.44
1.41
1.36
1.33
1.30
1.27
1.23
1.16
1.12
1.09
1.89
2.00
1.97
1.01
8.22
2.16
2.45
1.18
5E+07
4E+07
3E+07
2E+07
1E+07
5E+06
0
10.0 9.5 9.0 8.5 8.0 7.5 7.0 6.5 6.0 5.5 5.0 4.5 4.0 3.5 3.0 2.5 2.0 1.5 1.0 0.5 0.0 -0.5
f1 (ppm)

**16**

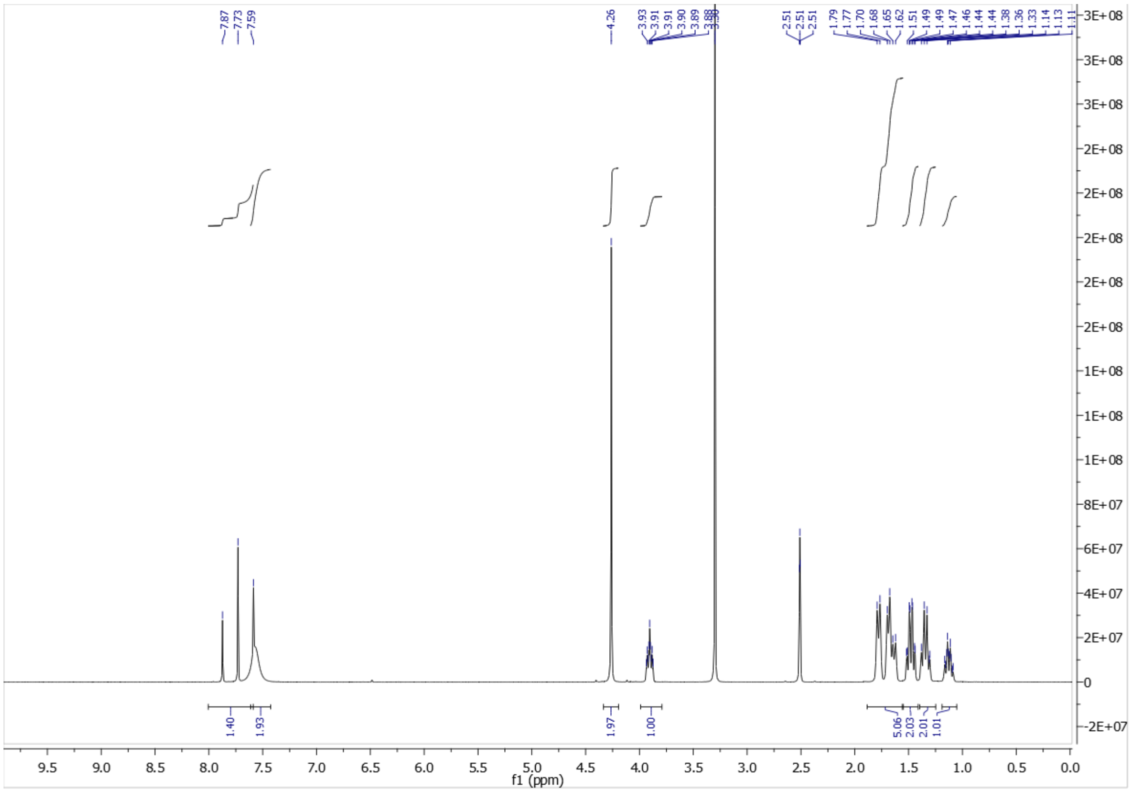

**17**

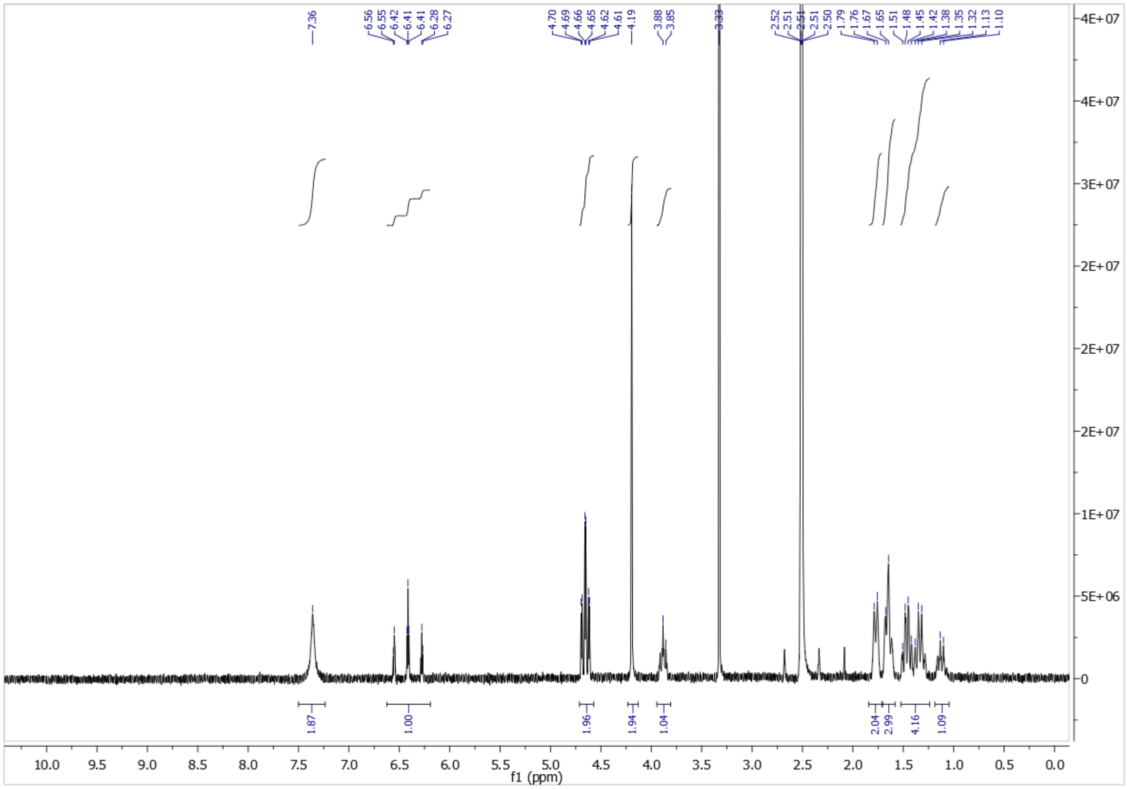

**18**

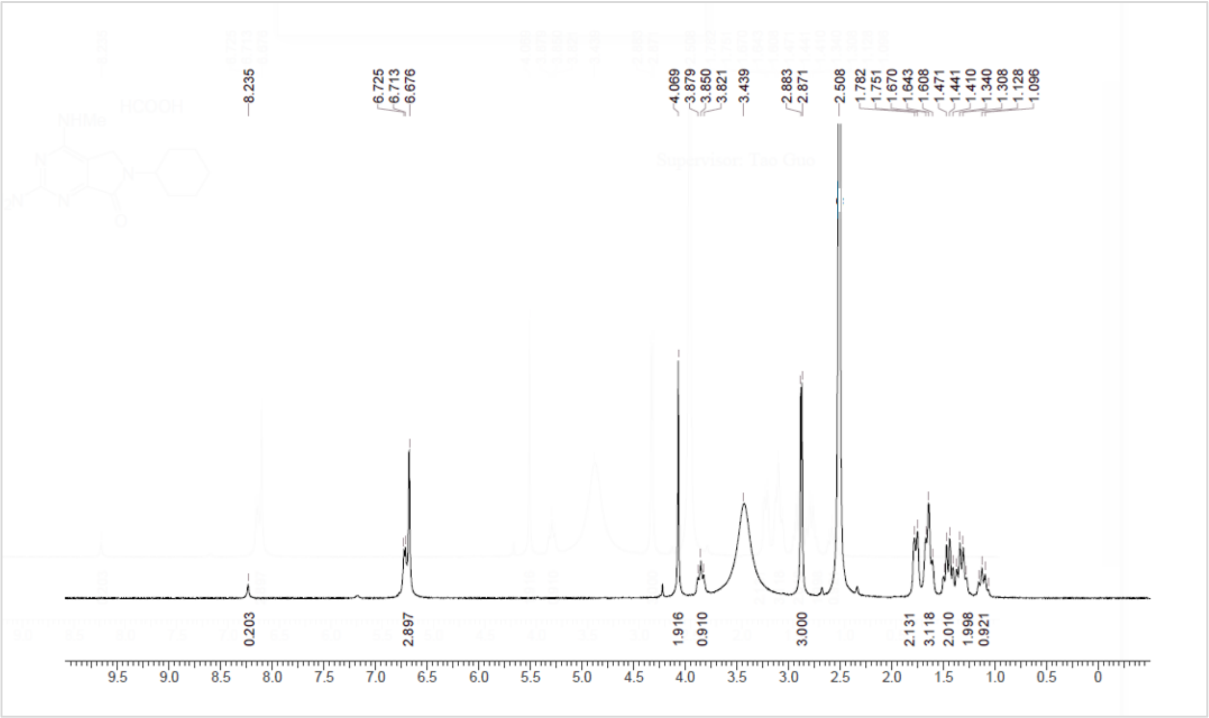

**19**

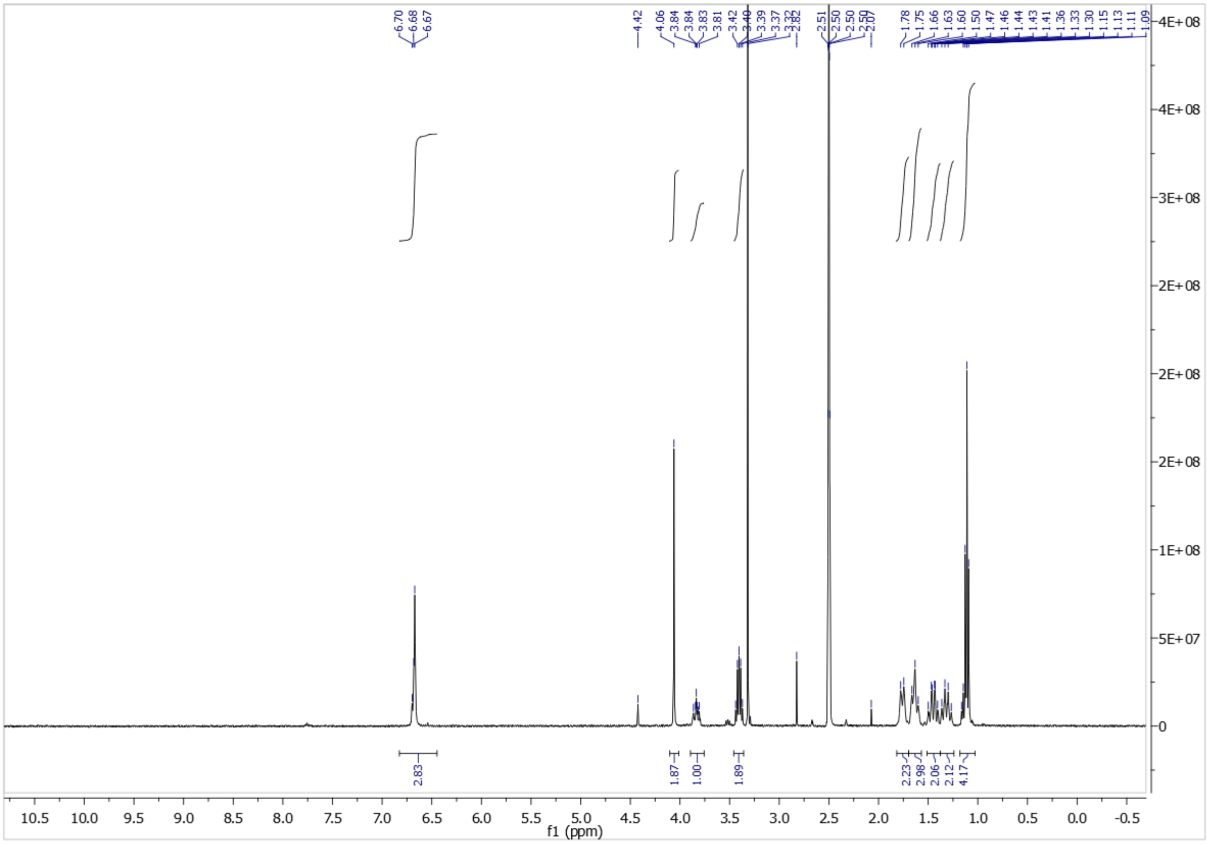

**20**

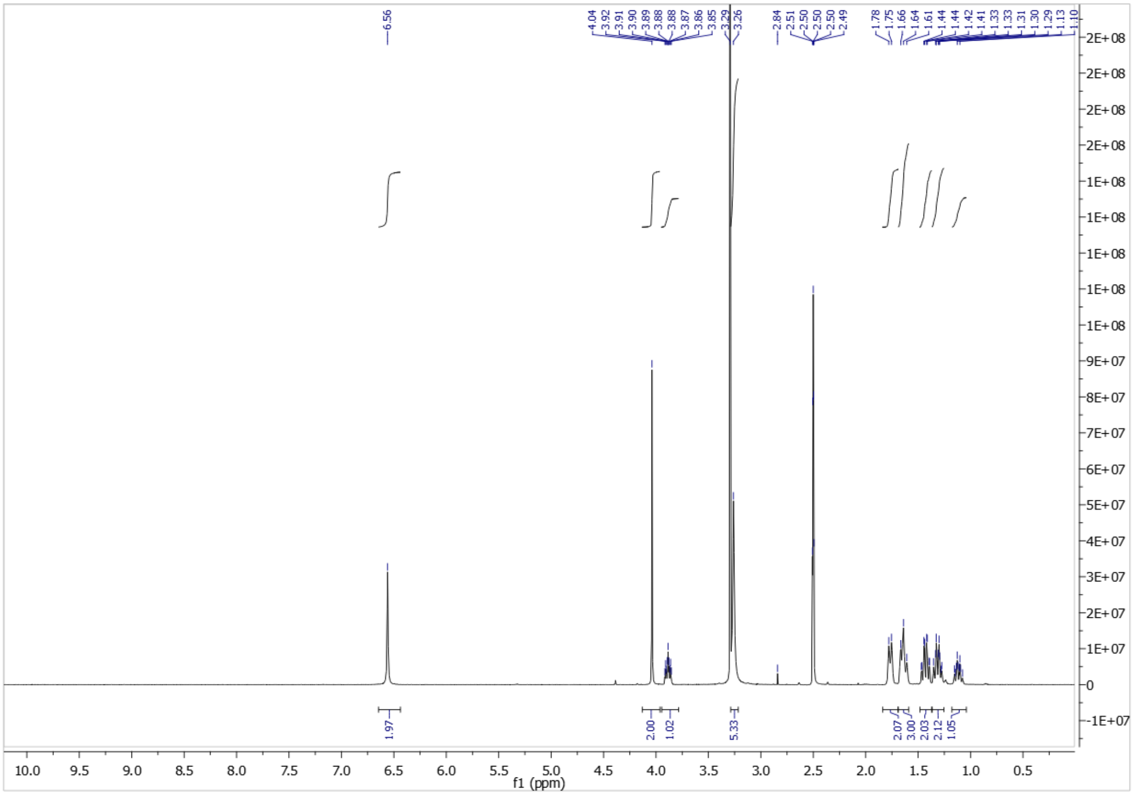

**21**

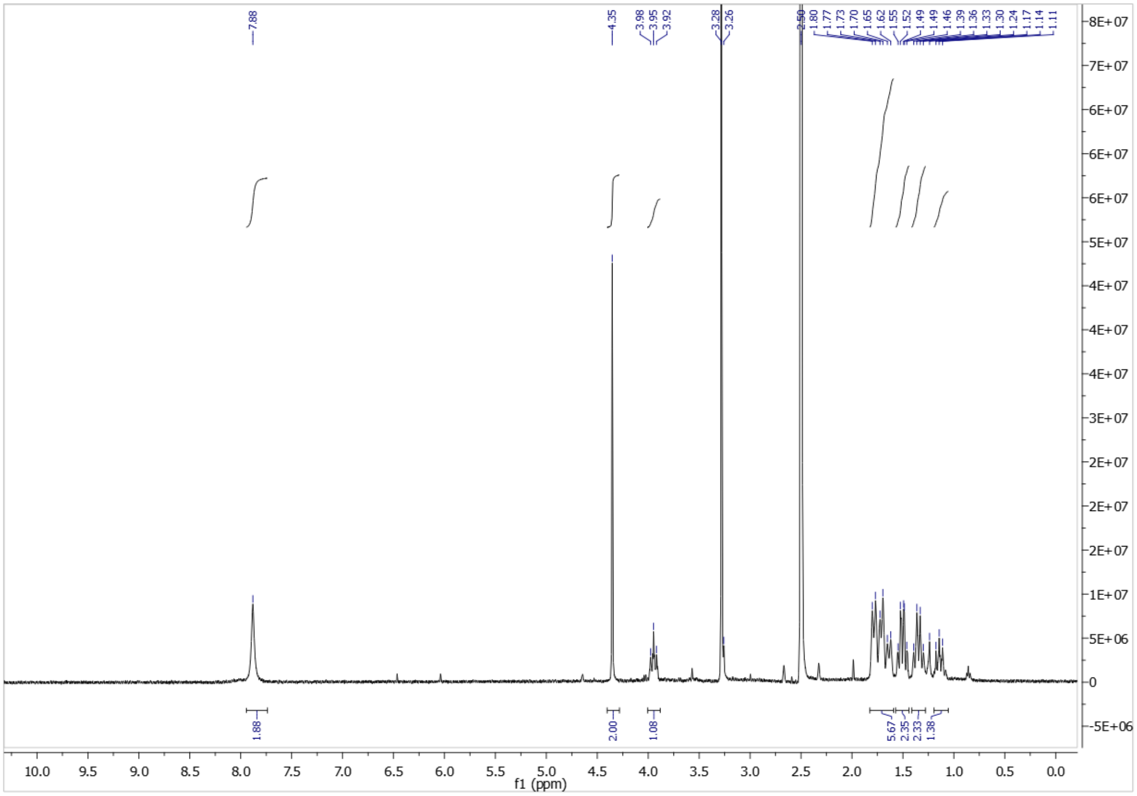

**22**

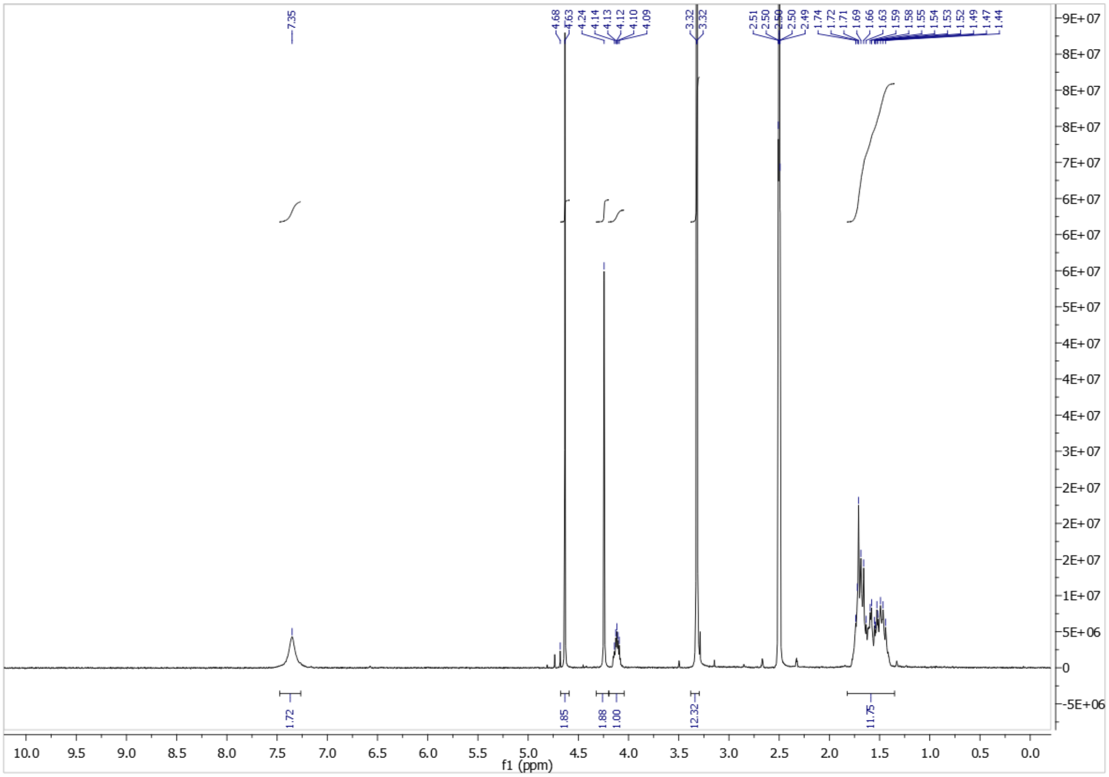


**23**

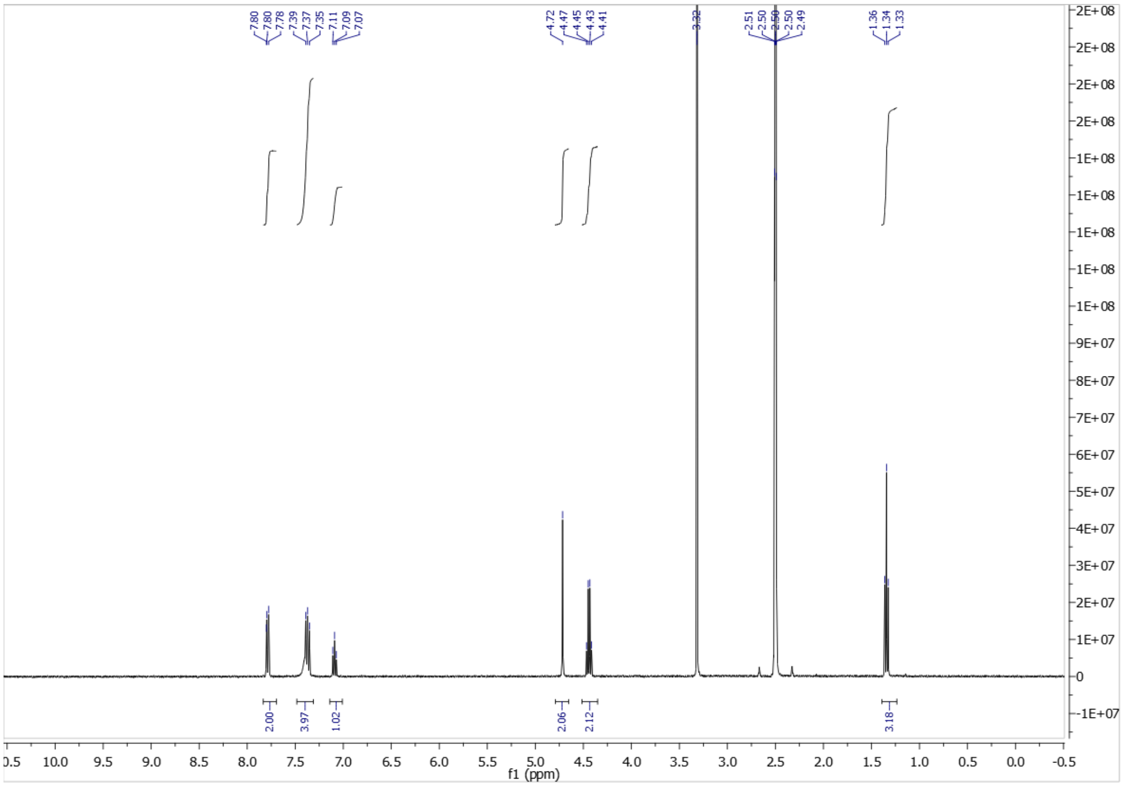

**26**

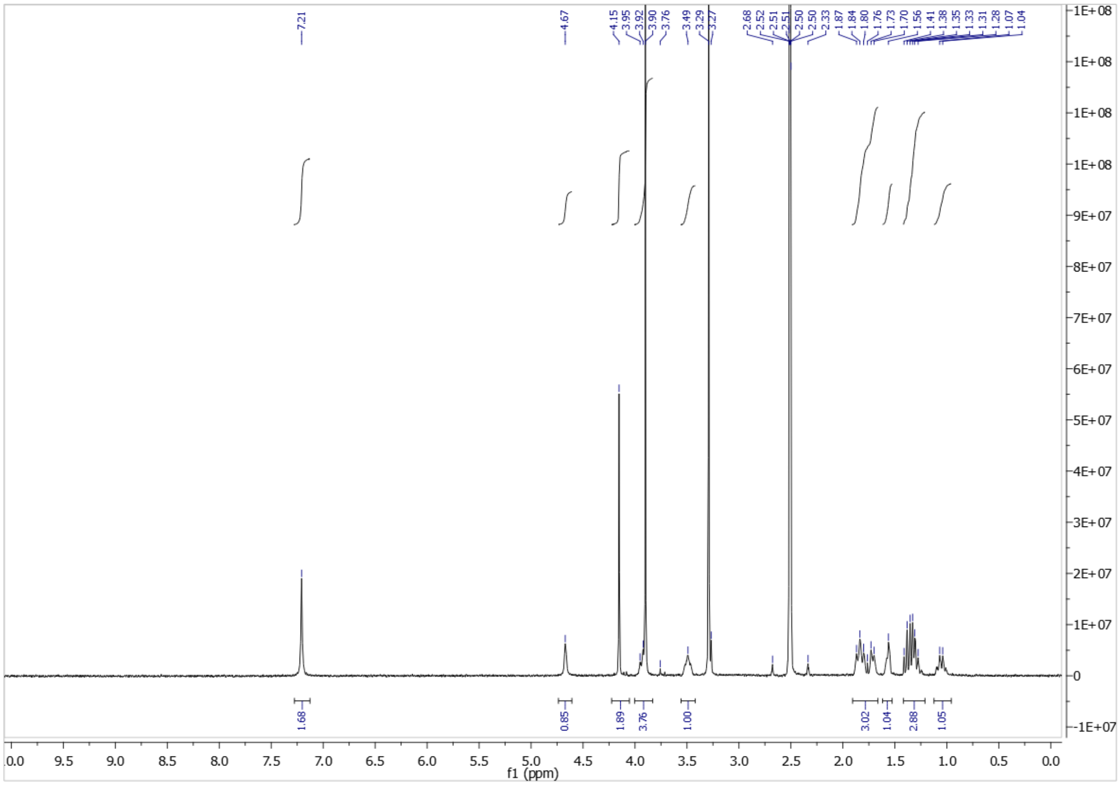


**27**

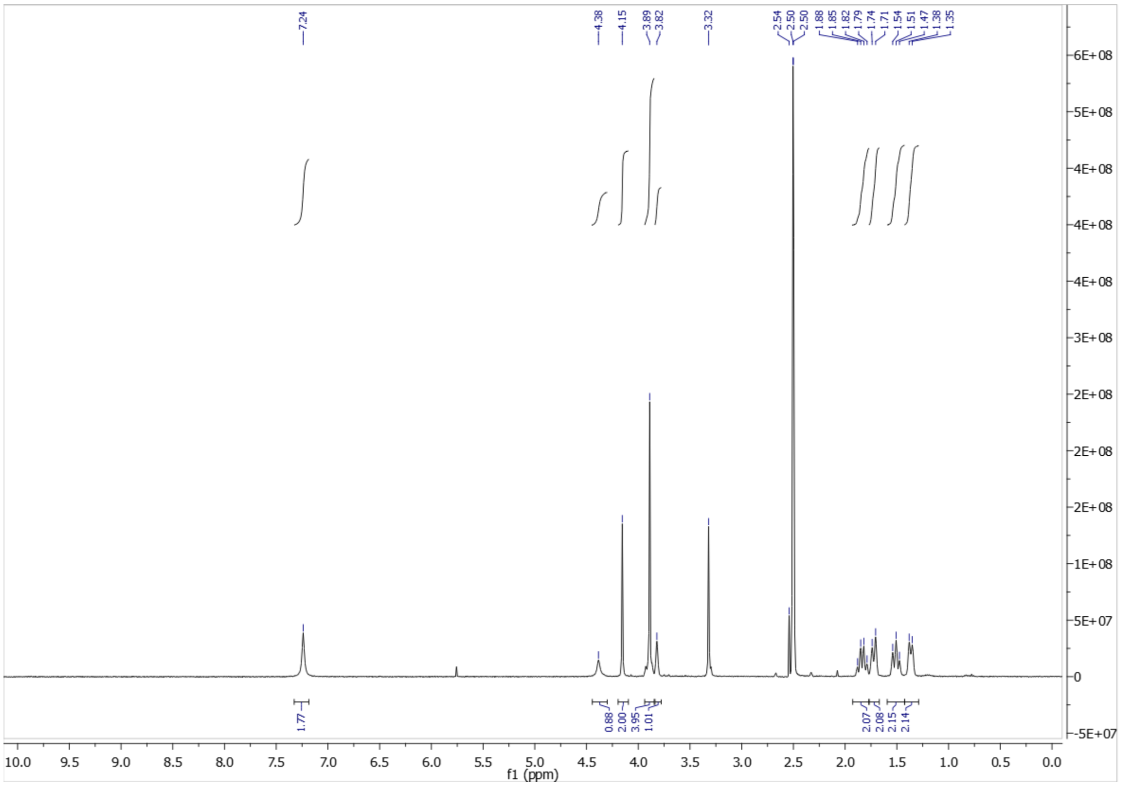

**28**

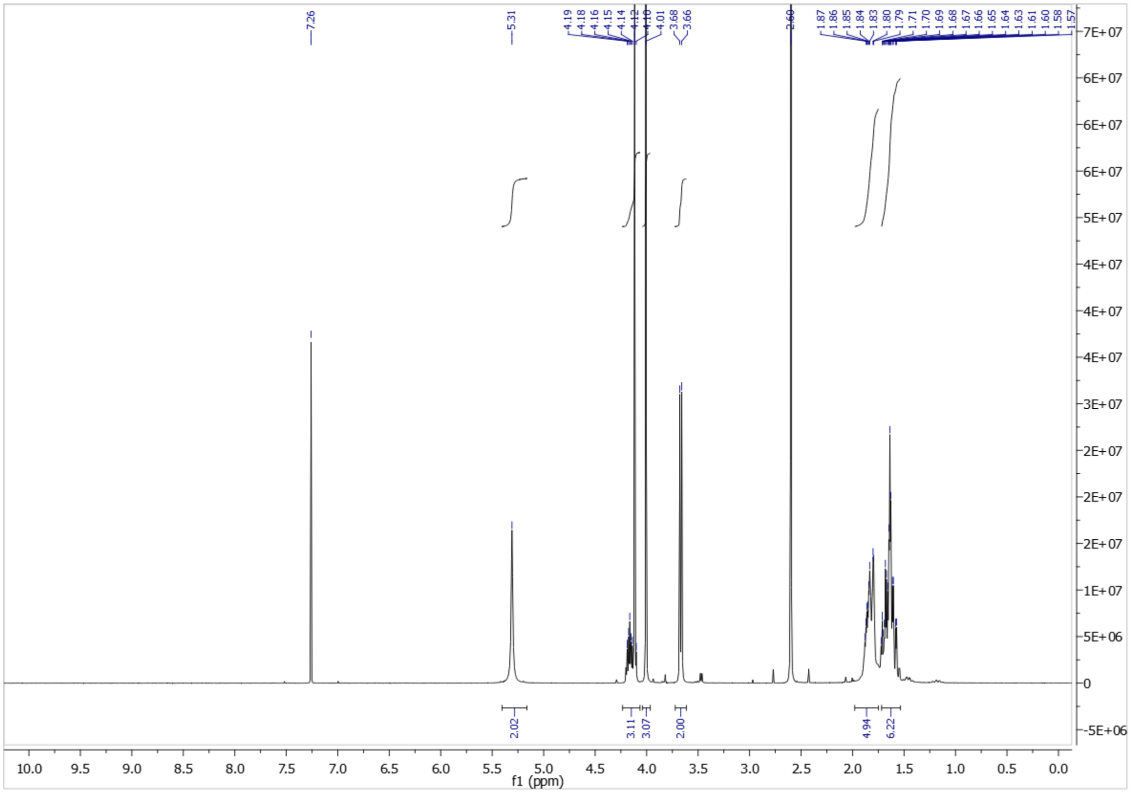

**29**

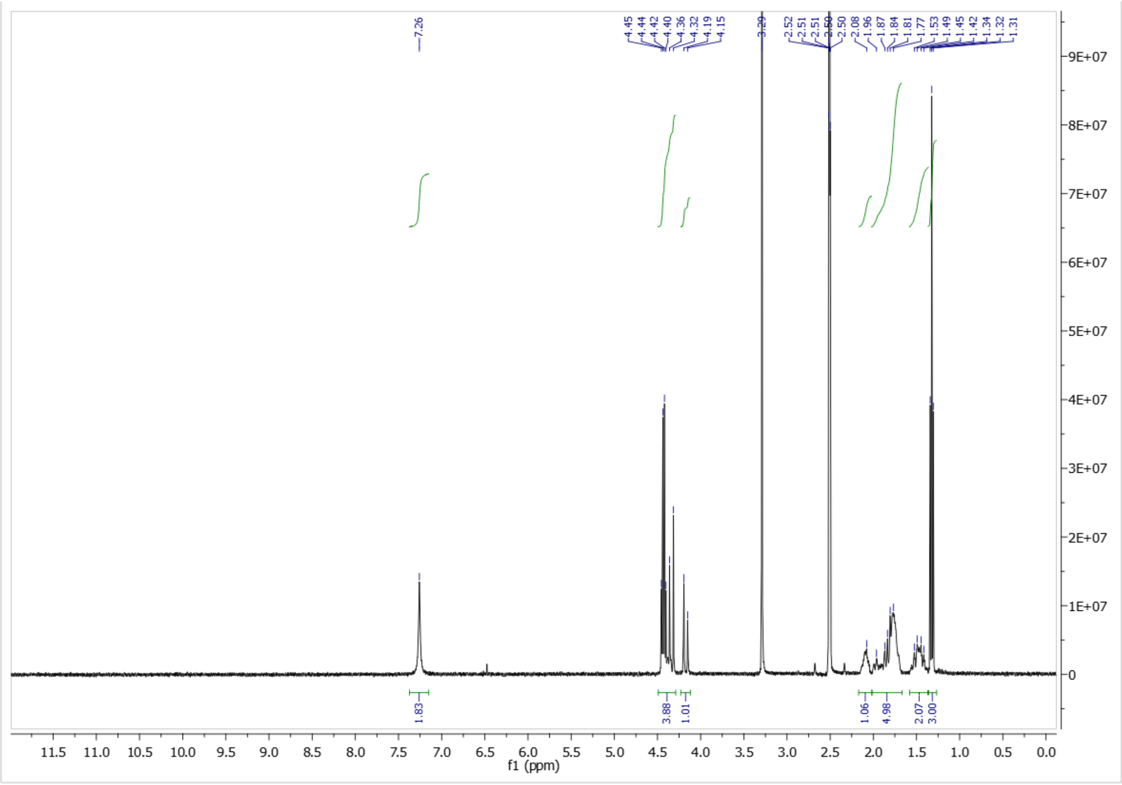

**30**

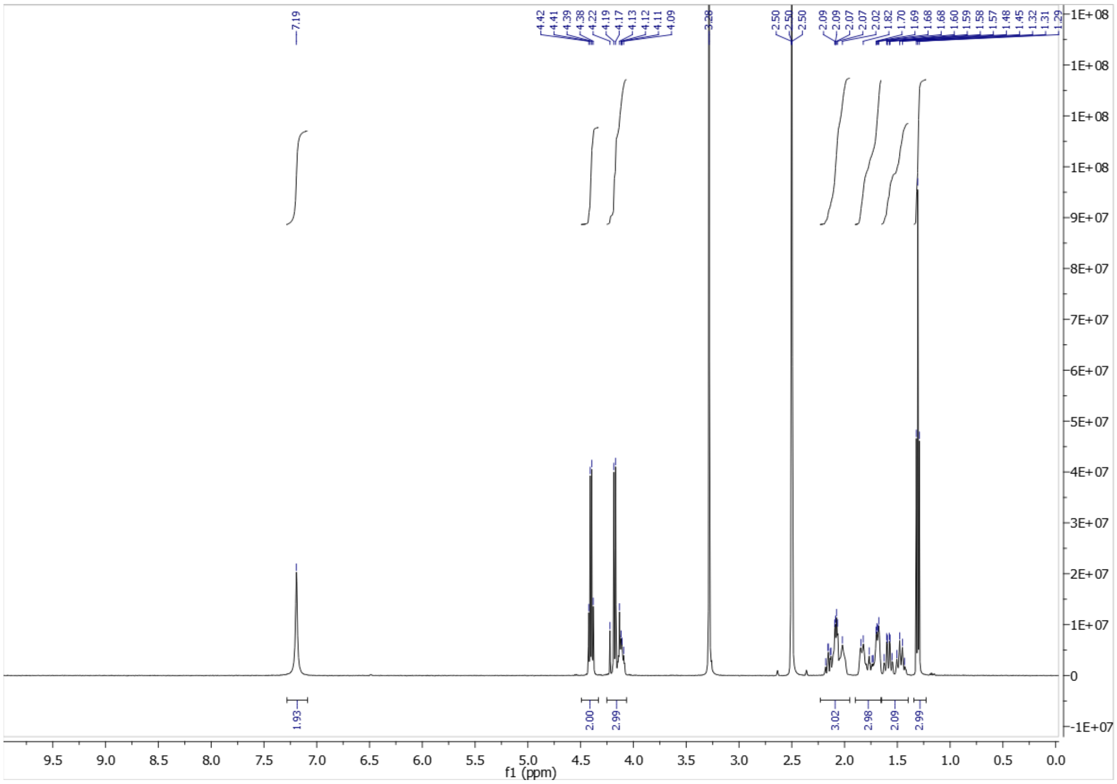
7.19
4.42 4.41 4.39 4.38 4.22 4.19 4.17 4.13 4.12 4.11 4.09
3.28
2.50 2.50 2.50
2.09 2.09 2.07 2.07 2.02 1.82 1.70 1.69 1.68 1.60 1.59 1.58 1.57 1.48 1.45 1.32 1.31 1.29
1.93 2.00 2.99 3.02 2.98 2.09 2.99
f1 (ppm)

**31**

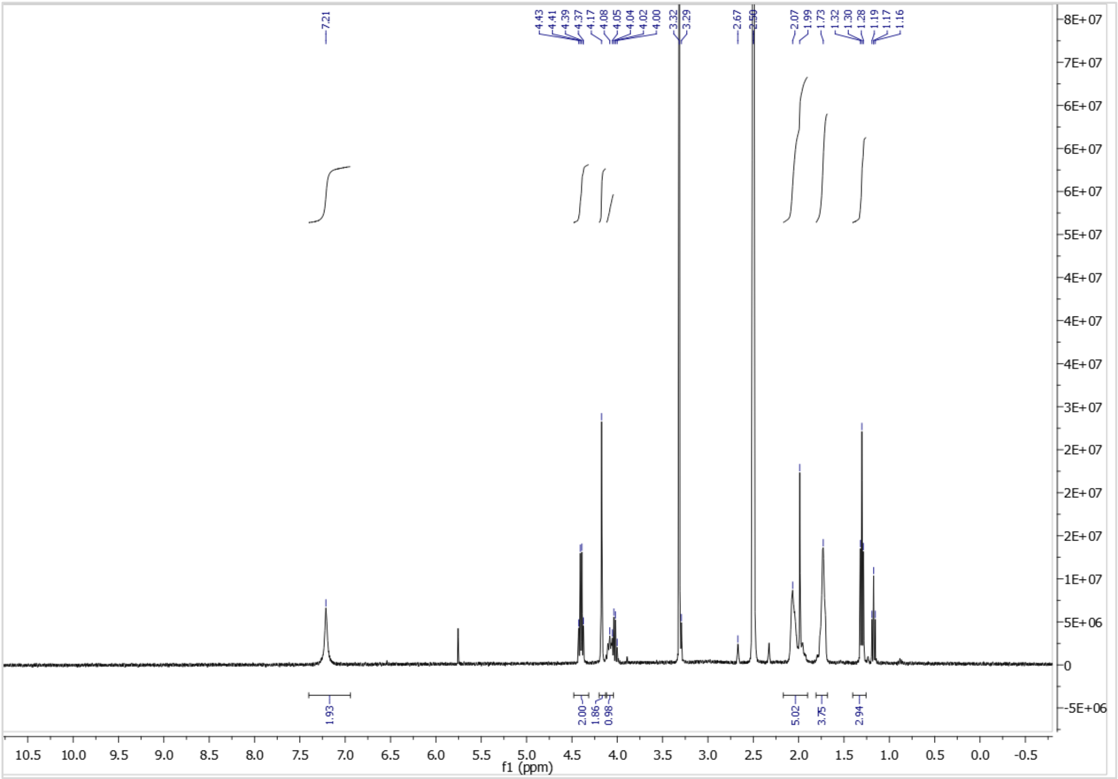
7.21
4.43 4.41 4.39 4.37 4.17 4.08 4.05 4.04 4.02 4.00
3.32 3.29
2.67 2.50
2.07 1.99 1.73 1.32 1.30 1.28 1.19 1.17 1.16
1.93 2.00 1.86 0.98 5.02 3.75 2.94
f1 (ppm)

**33**

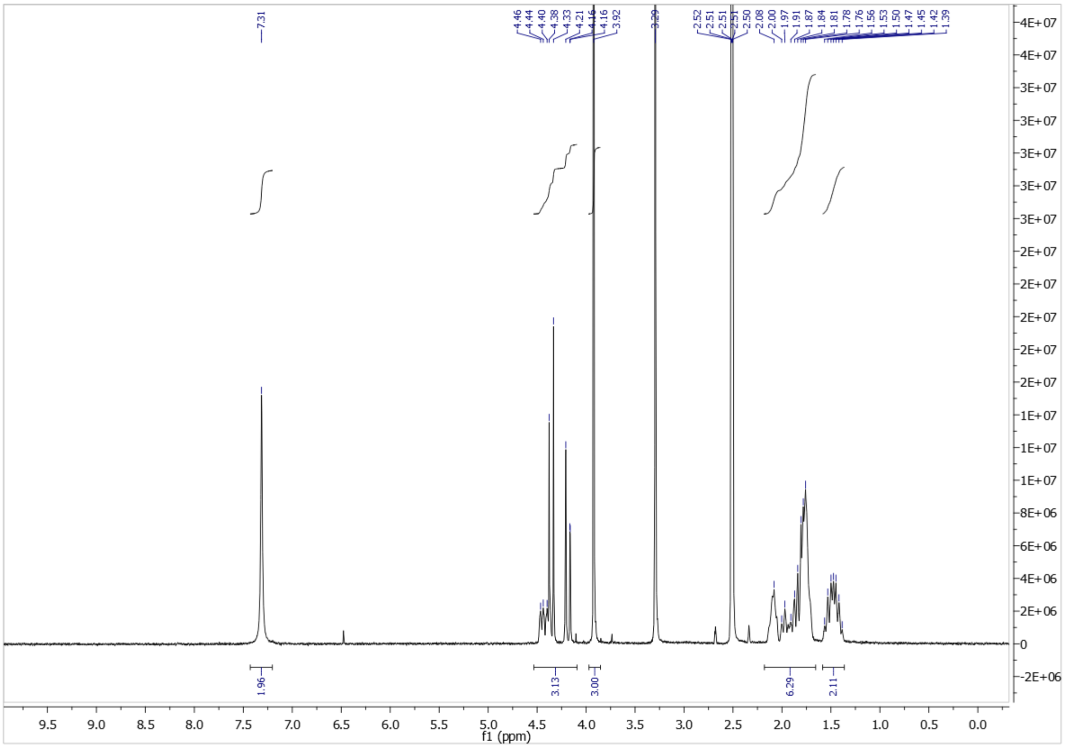

**34**

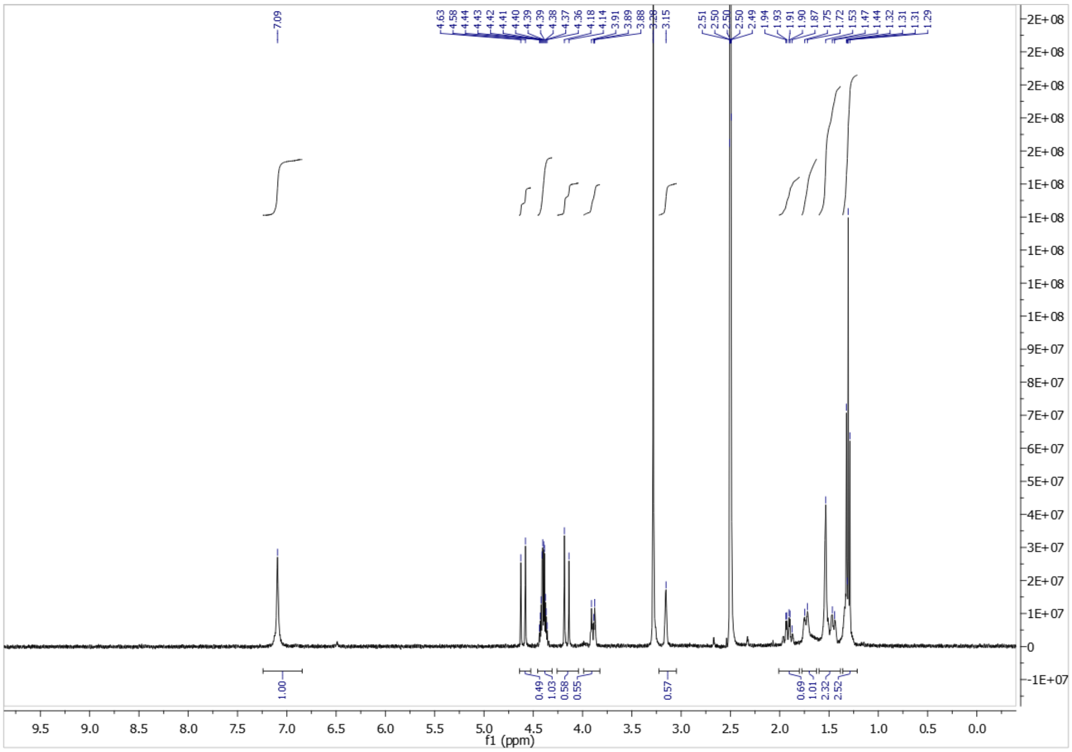

**35**

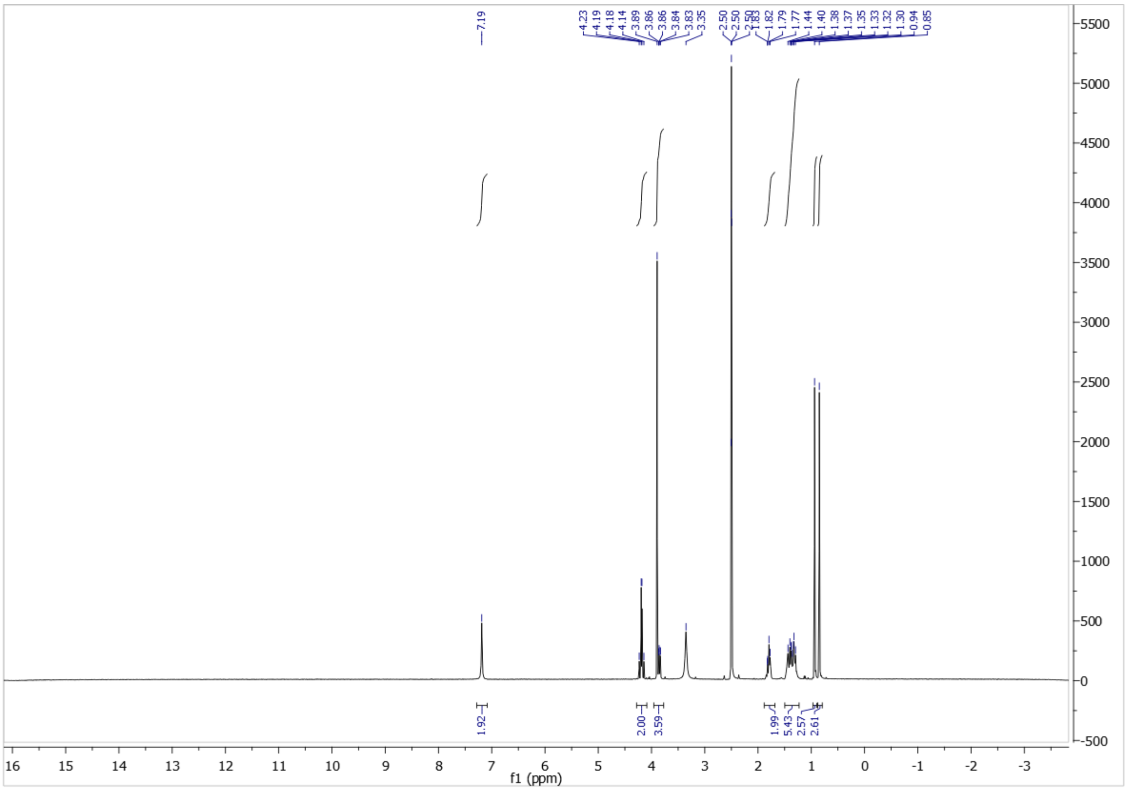

**38**

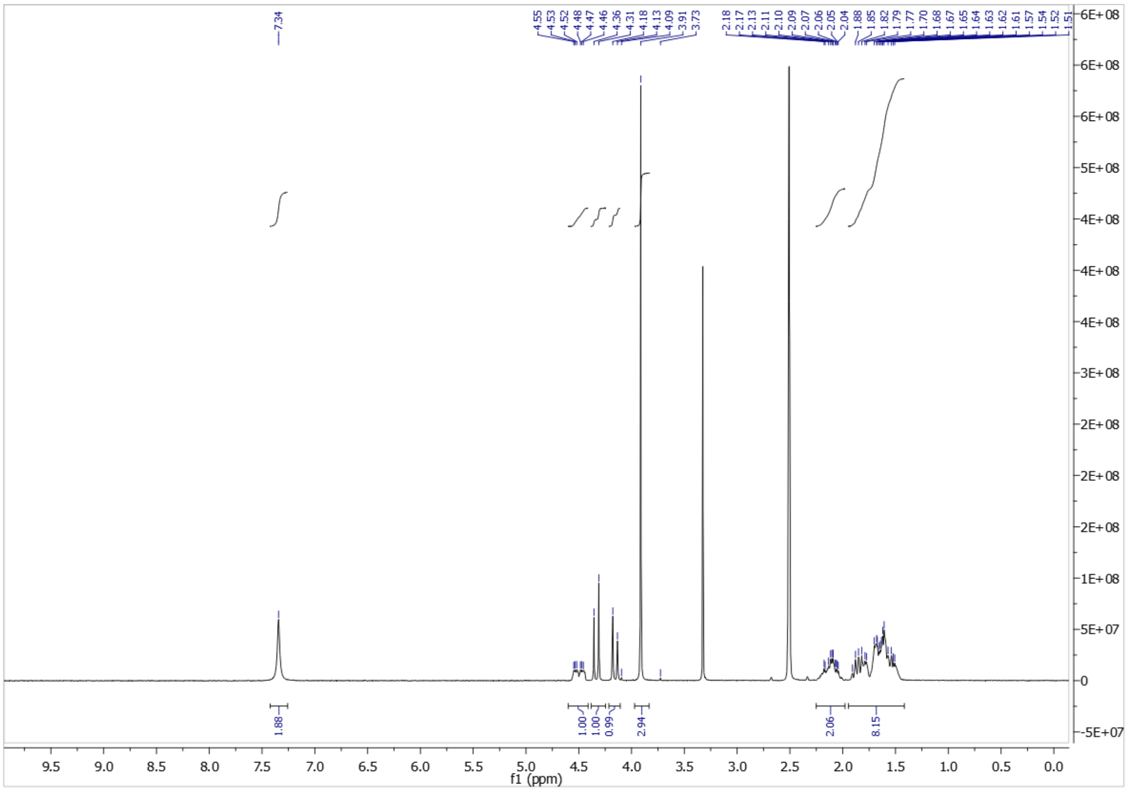

**39**

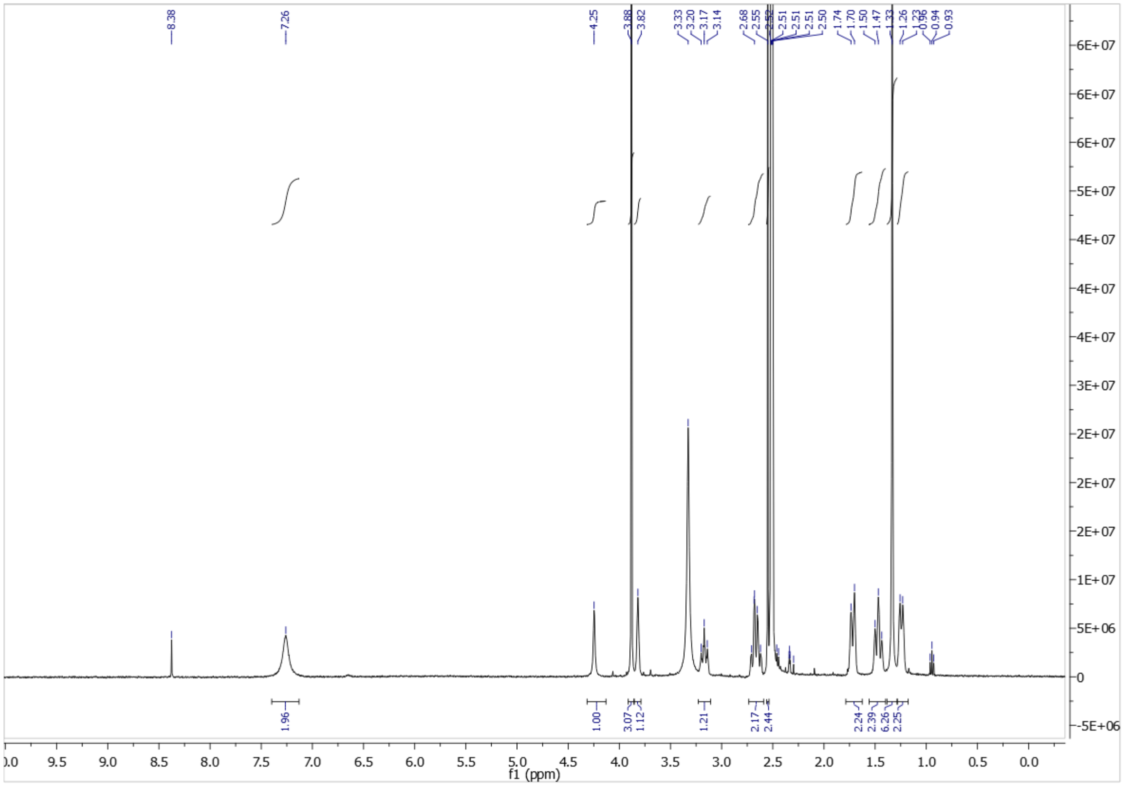

**40**

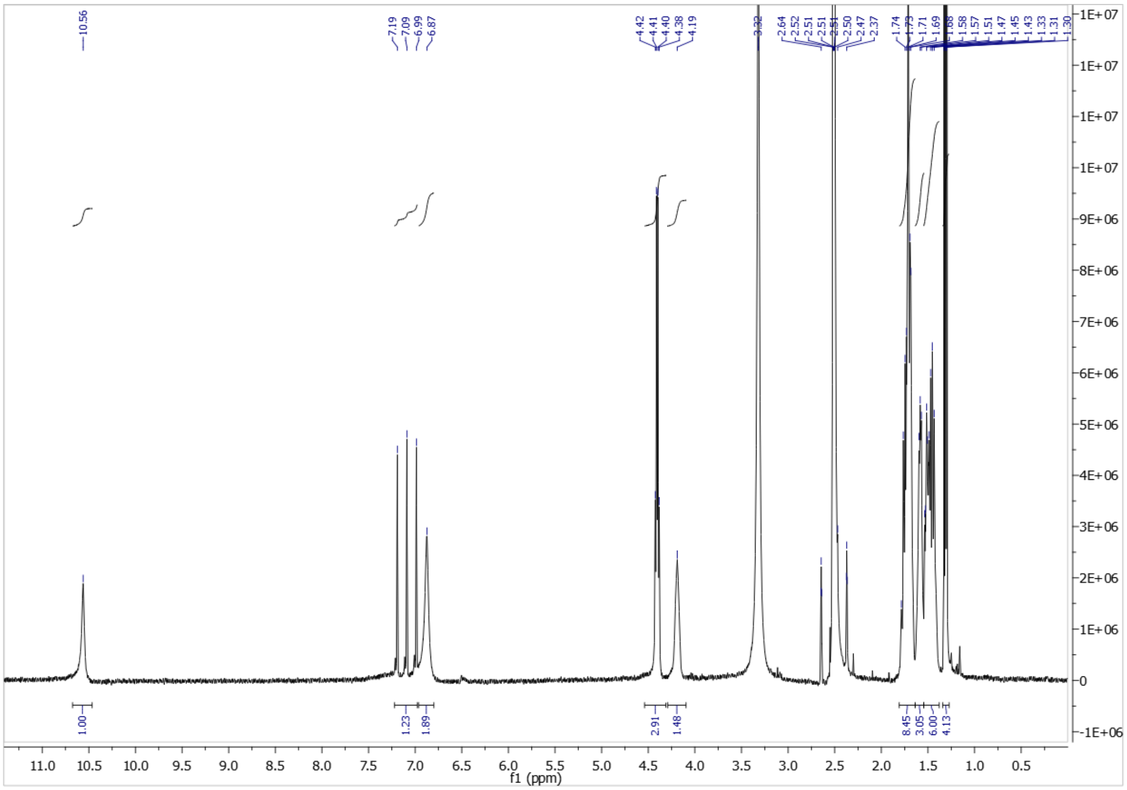

**41**

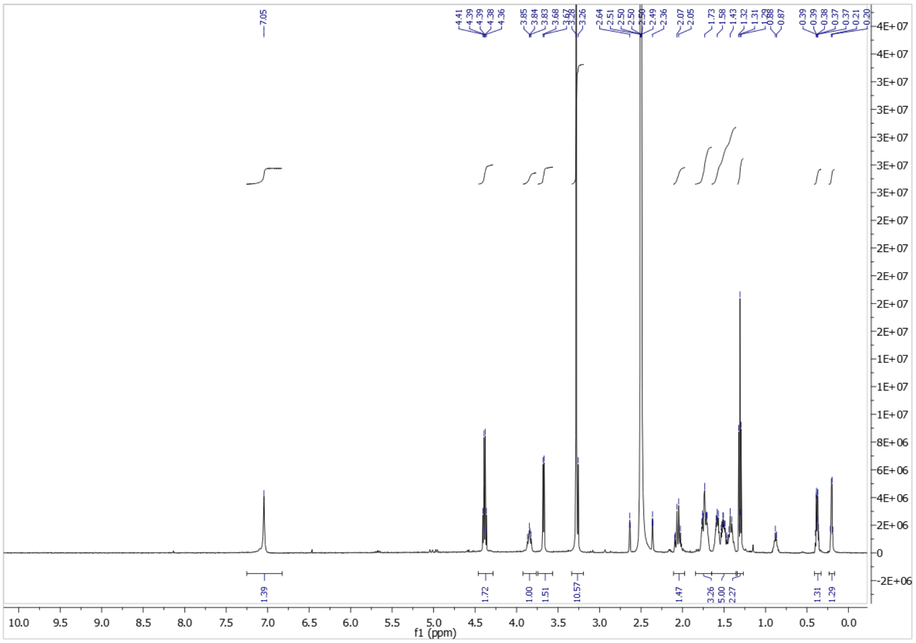

Integration did not correlate as expected.  HRMS included to confirm purity level was acceptable for initial profiling.

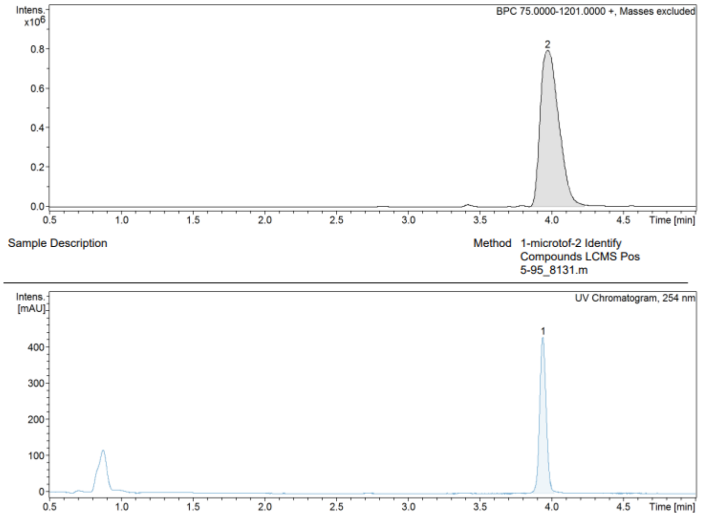


| RT [min] | Area Frac. % | Chromatogram |
|---|---|---|
| 3.9 | 100.00 | UV Chromatogram, 254 nm |
| 4.0 | 100.00 | BPC 75.0000-1201.0000 +, Masses excluded |

**Cmpd 2, 4.0 min**

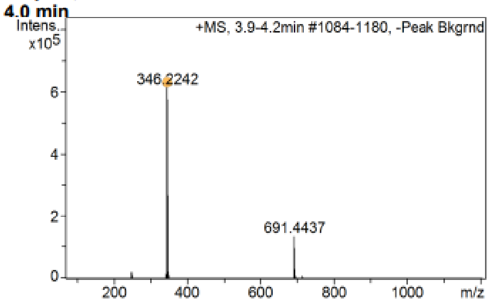

**42**

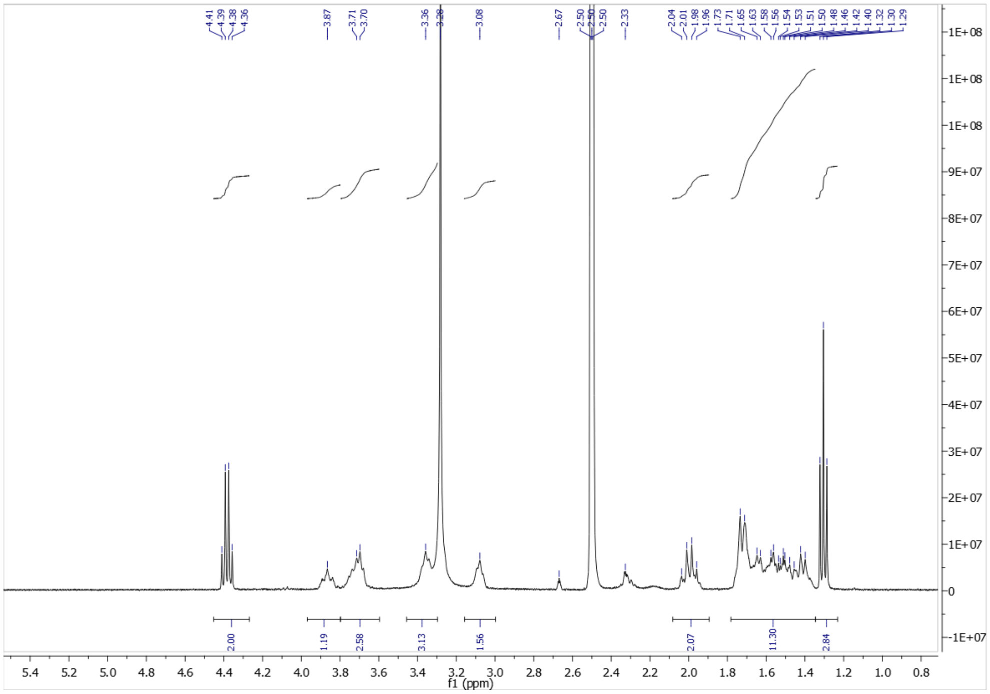


Integration did not correlate as expected. HRMS included to confirm purity level was acceptable for initial profiling.

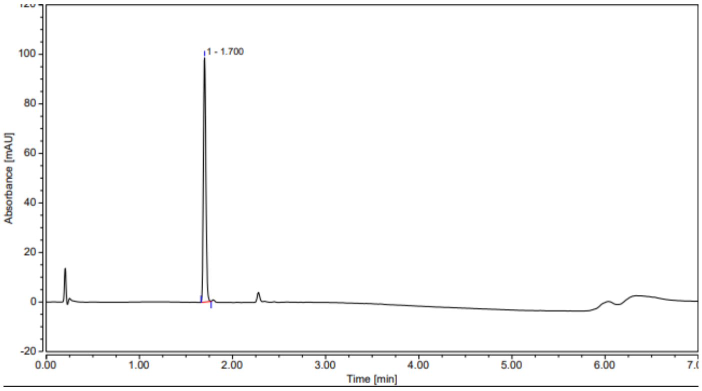


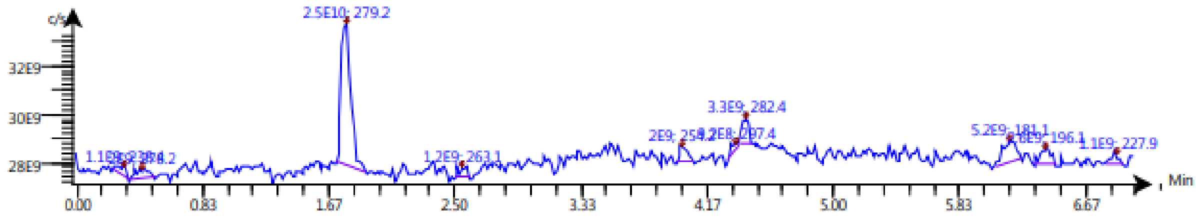


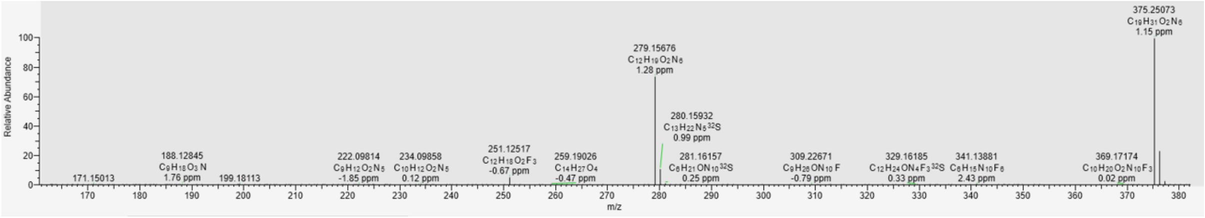

**43**

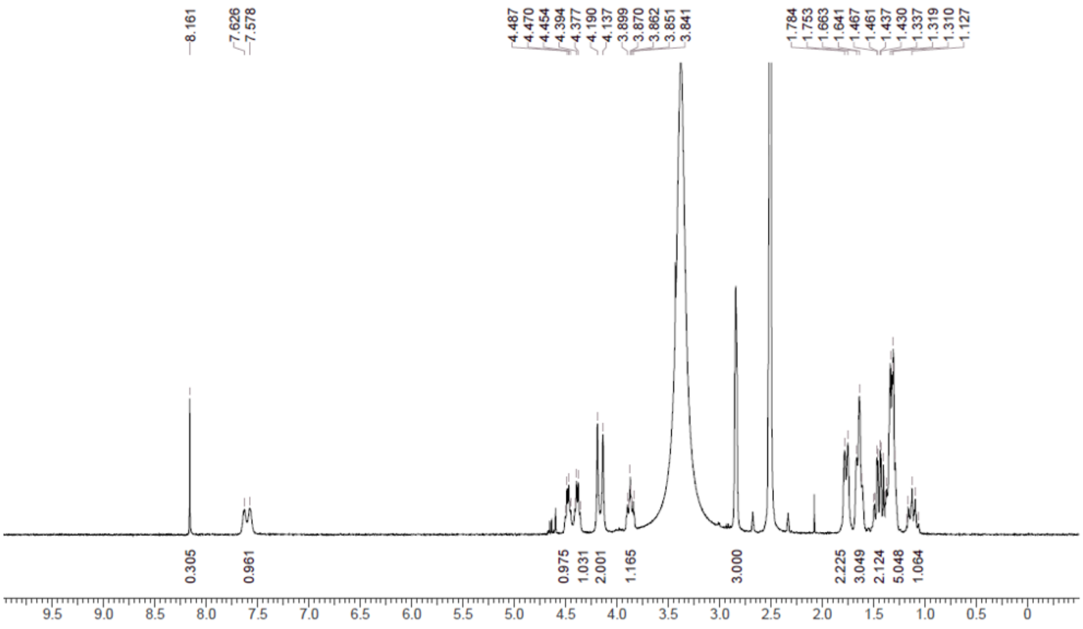

**44**

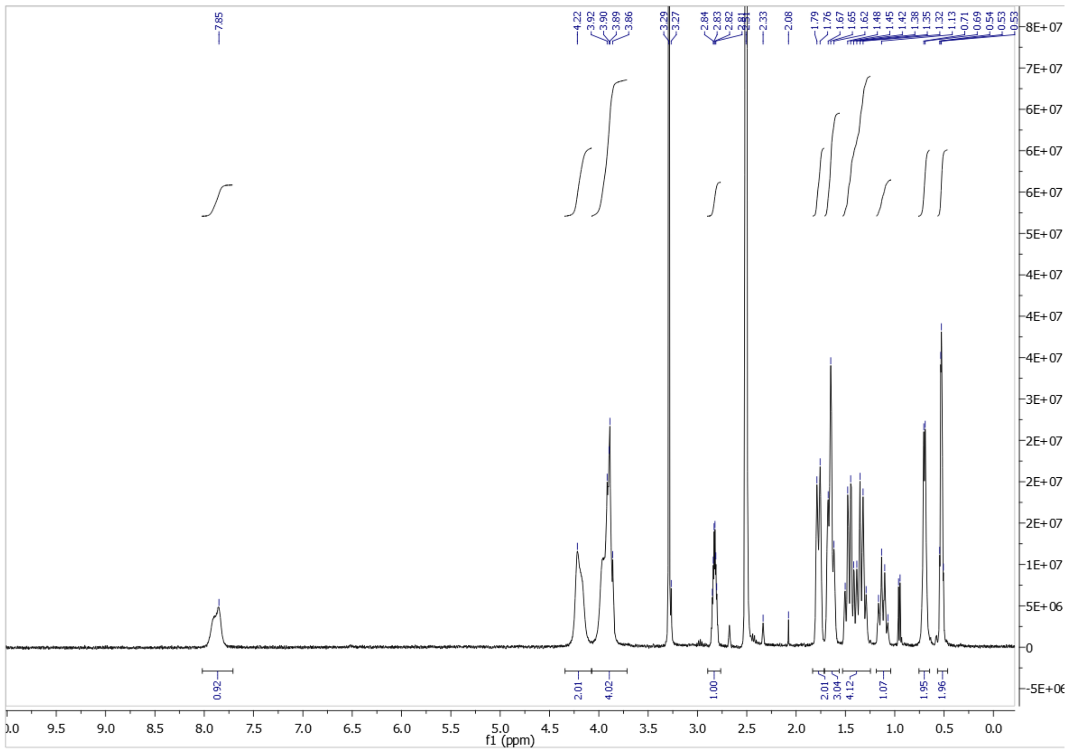

**45**

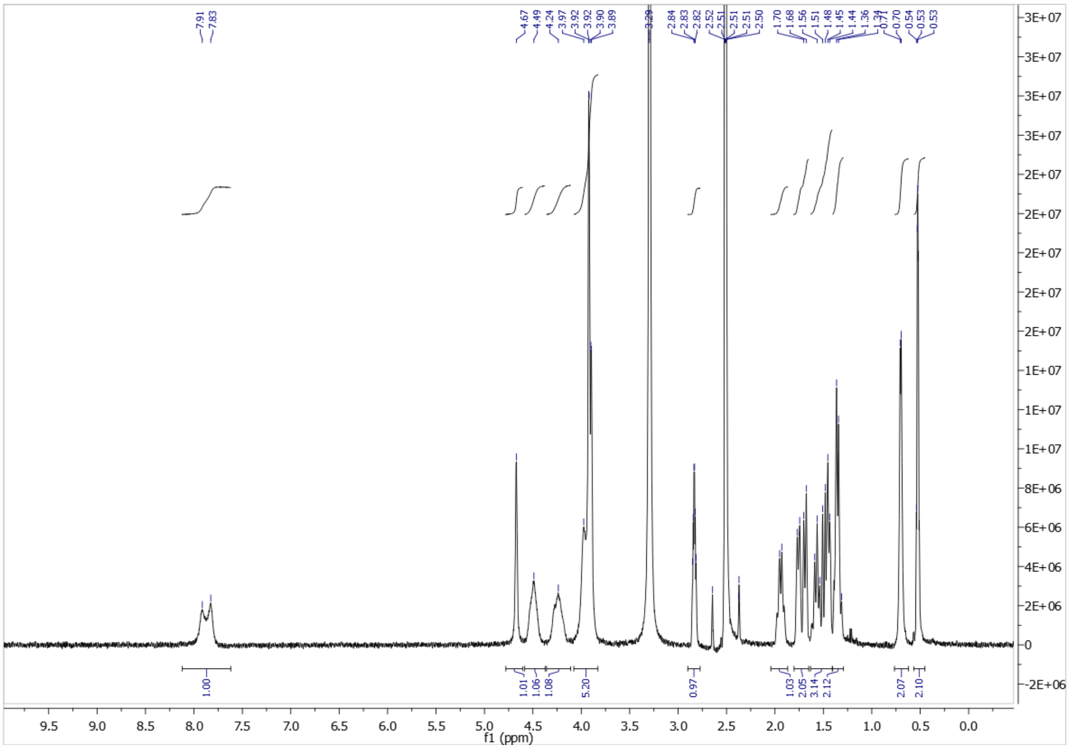


**48**

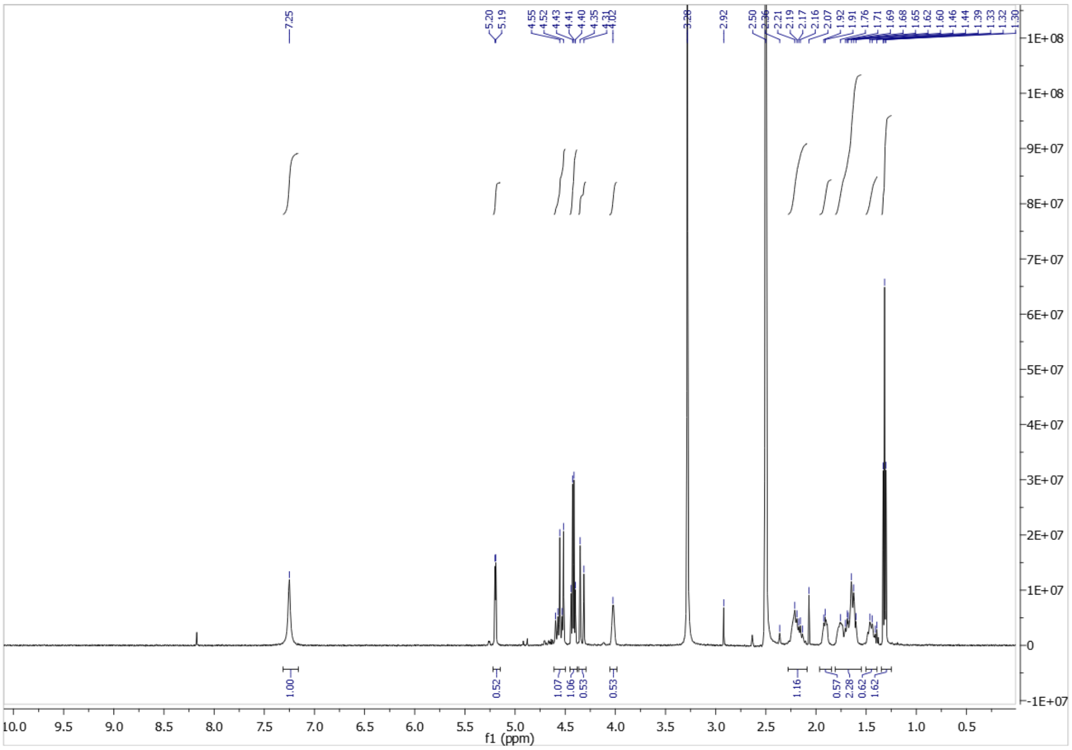

### HRMS Spectra for compounds progressed to in vivo studies

**8**

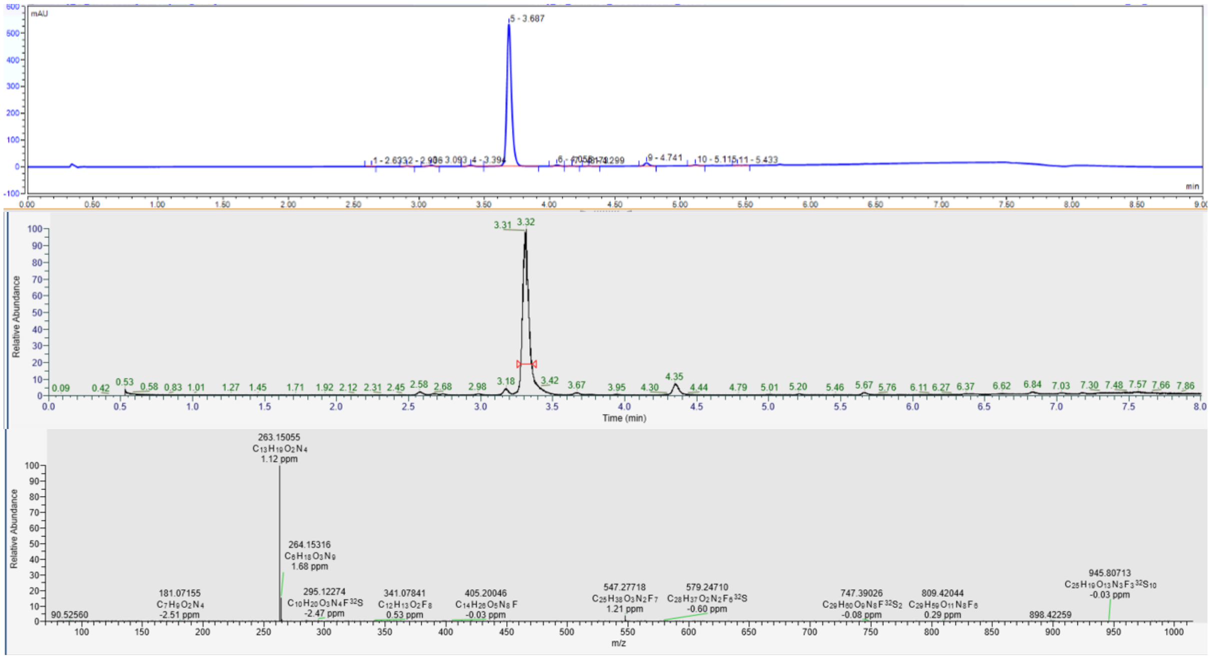


**10**

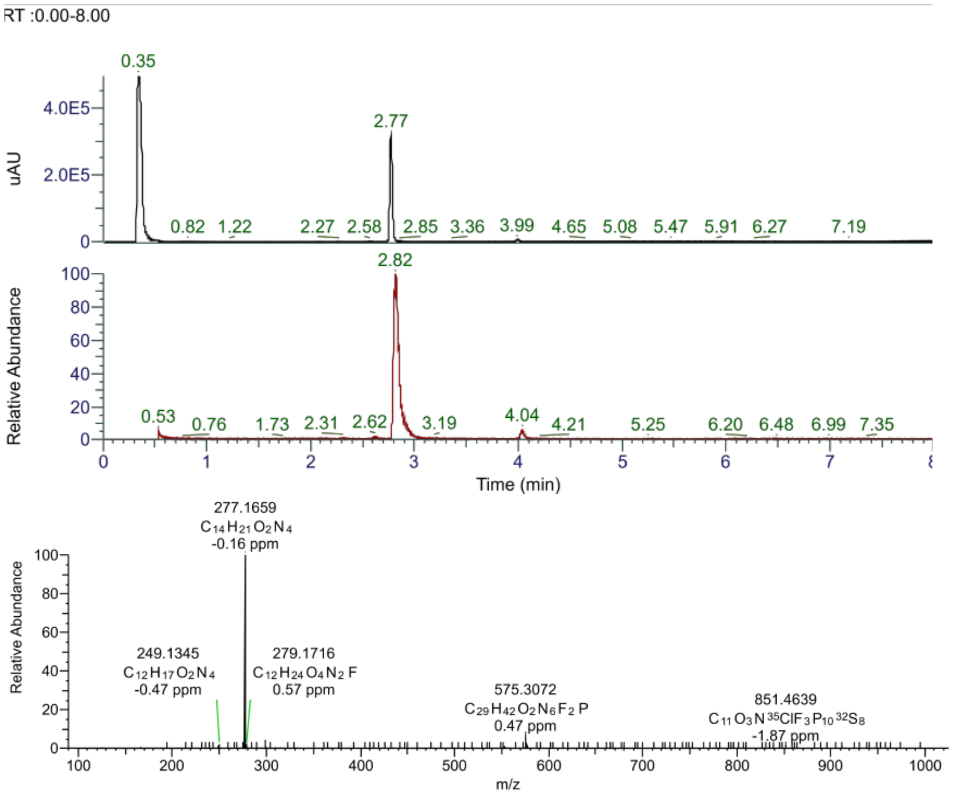

**11**

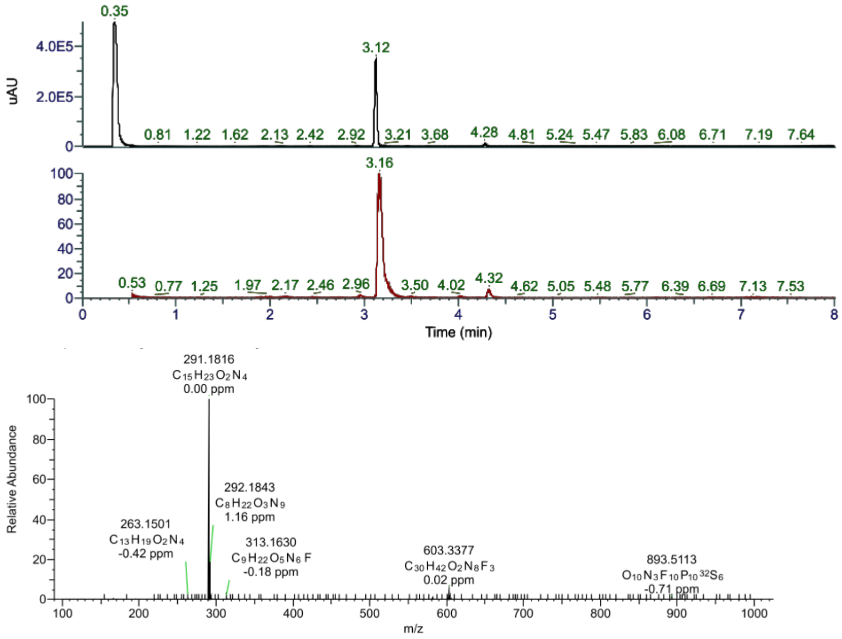

0.35
3.12
uAU
4.0E5
2.0E5
0.81 1.22 1.62 2.13 2.42 2.92 3.21 3.68 4.28 4.81 5.24 5.47 5.83 6.08 6.71 7.19 7.64
3.16
0.53 0.77 1.25 1.97 2.17 2.46 2.96 3.50 4.02 4.32 4.62 5.05 5.48 5.77 6.39 6.69 7.13 7.53
Time (min)
291.1816
C15H23O2N4
0.00 ppm
Relative Abundance
292.1843
C8H22O3N9
1.16 ppm
263.1501
C13H19O2N4
-0.42 ppm
313.1630
C9H22O5N6F
-0.18 ppm
603.3377
C30H42O2N8F3
0.02 ppm
893.5113
O10N3F10P10 32S6
-0.71 ppm
m/z


**25**

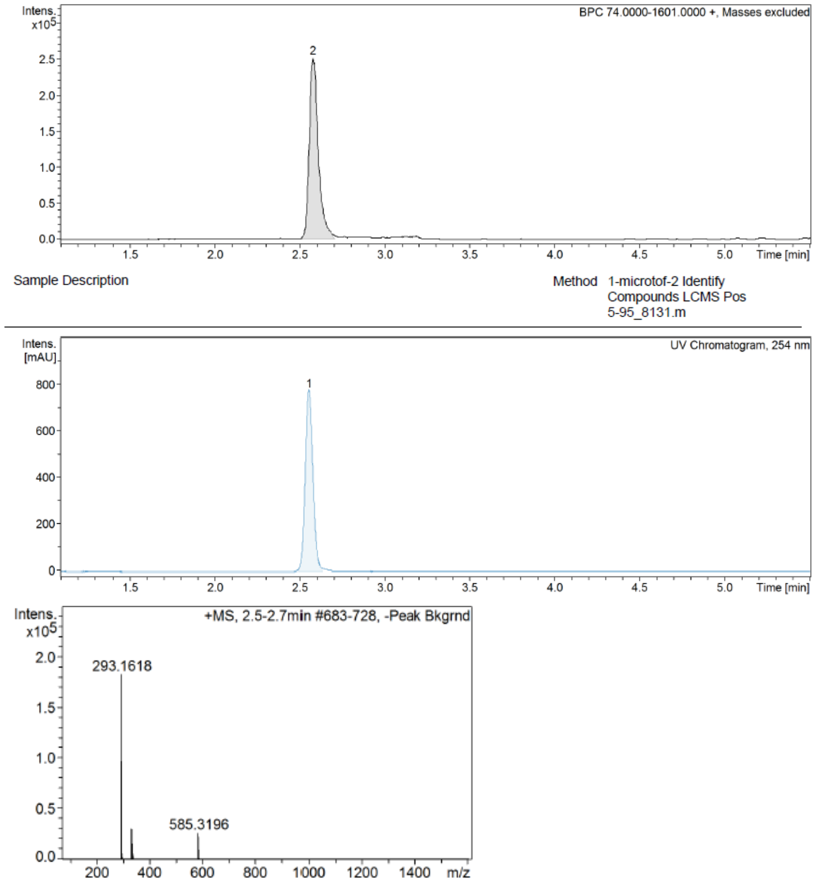

Intens. x10^5
BPC 74.0000-1601.0000 +, Masses excluded
2
Time [min]
Sample Description
Method 1-microtof-2 Identify Compounds LCMS Pos 5-95_8131.m
Intens. [mAU]
UV Chromatogram, 254 nm
1
Time [min]
Intens. x10^5
+MS, 2.5-2.7min #683-728, -Peak Bkgrnd
293.1618
585.3196
m/z

**32**

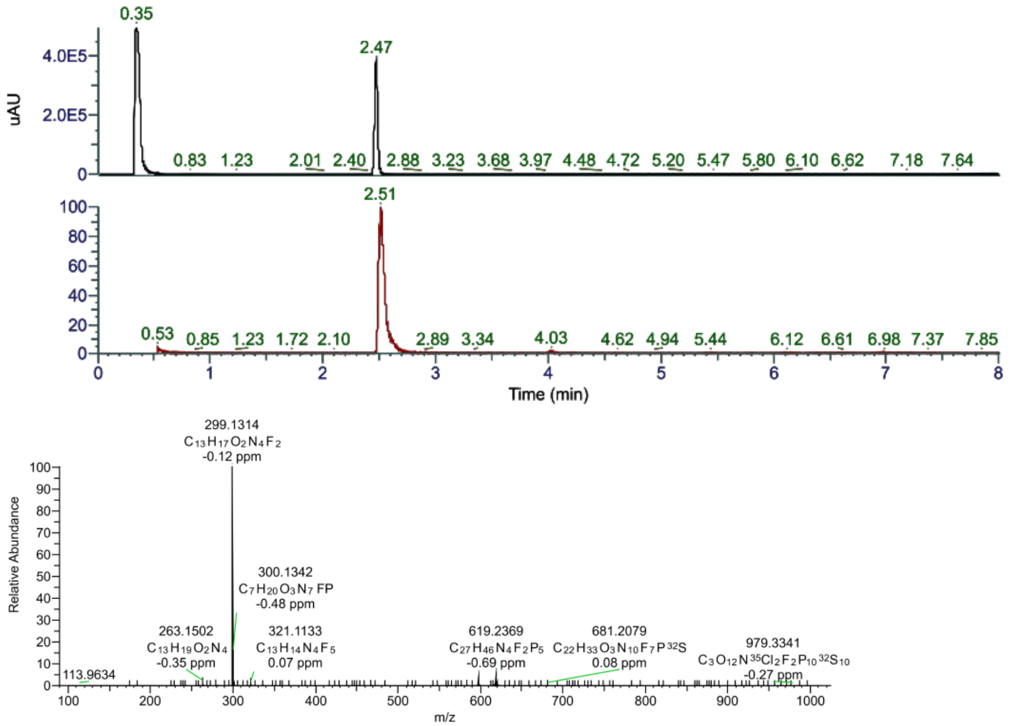

0.35
2.47
4.0E5
2.0E5
uAU
0.83 1.23 2.01 2.40 2.88 3.23 3.68 3.97 4.48 4.72 5.20 5.47 5.80 6.10 6.62 7.18 7.64
2.51
0.53 0.85 1.23 1.72 2.10 2.89 3.34 4.03 4.62 4.94 5.44 6.12 6.61 6.98 7.37 7.85
Time (min)
299.1314
C13H17O2N4F2
-0.12 ppm
300.1342
C7H20O3N7FP
-0.48 ppm
263.1502
C13H19O2N4
-0.35 ppm
321.1133
C13H14N4F5
0.07 ppm
619.2369
C27H46N4F2P5
-0.69 ppm
681.2079
C22H33O3N10F7P32S
0.08 ppm
979.3341
C3O12N35Cl2F2P10 32S10
-0.27 ppm
113.9634
Relative Abundance
m/z


**37**

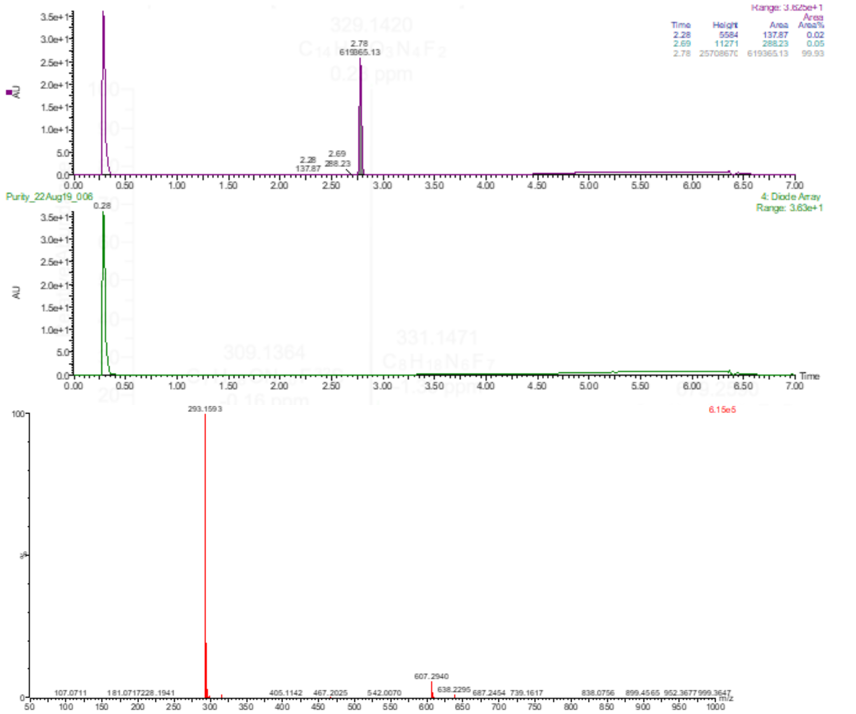

2.78
Purity_22Aug19_006
4: Diode Array
Range: 3.63e+1
AU
Time
293.1593
6.15e5
607.2940

**38**

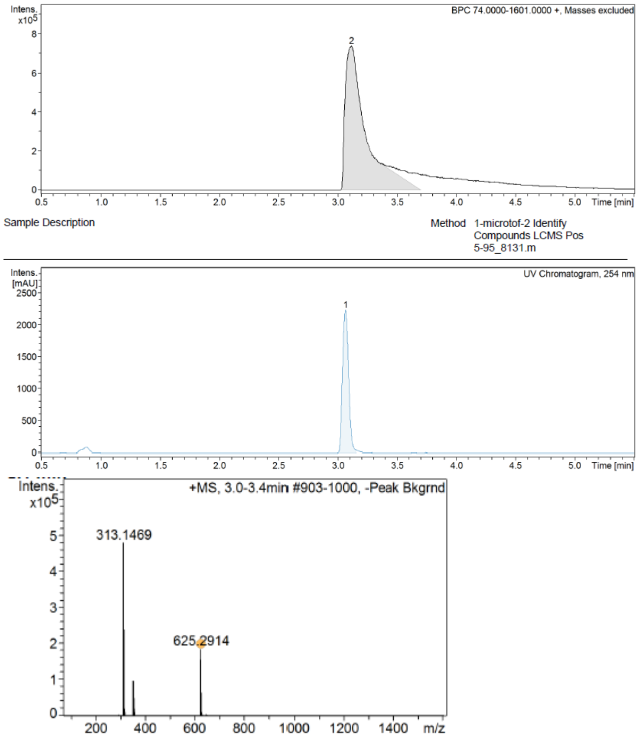
BPC 74.0000-1601.0000 +, Masses excluded
Intens. x10^5
Time [min]
Sample Description
Method 1-microtof-2 Identify Compounds LCMS Pos 5-95_8131.m
UV Chromatogram, 254 nm
Intens. [mAU]
+MS, 3.0-3.4min #903-1000, -Peak Bkgrnd
313.1469
625.2914
m/z

**46**

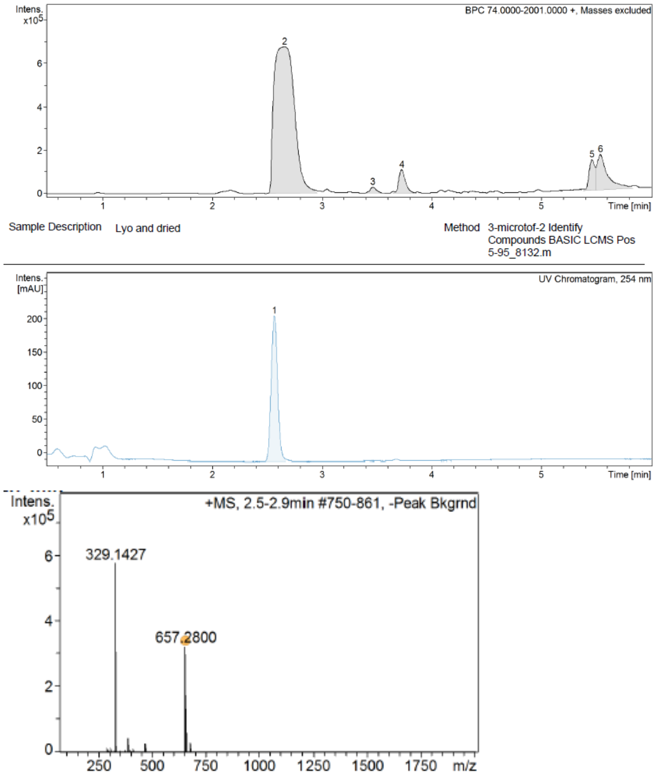
BPC 74.0000-2001.0000 +, Masses excluded
Intens. x10^5
Time [min]
Sample Description Lyo and dried
Method 3-microtof-2 Identify Compounds BASIC LCMS Pos 5-95_8132.m
UV Chromatogram, 254 nm
Intens. [mAU]
+MS, 2.5-2.9min #750-861, -Peak Bkgrnd
329.1427
657.2800
m/z

**47**

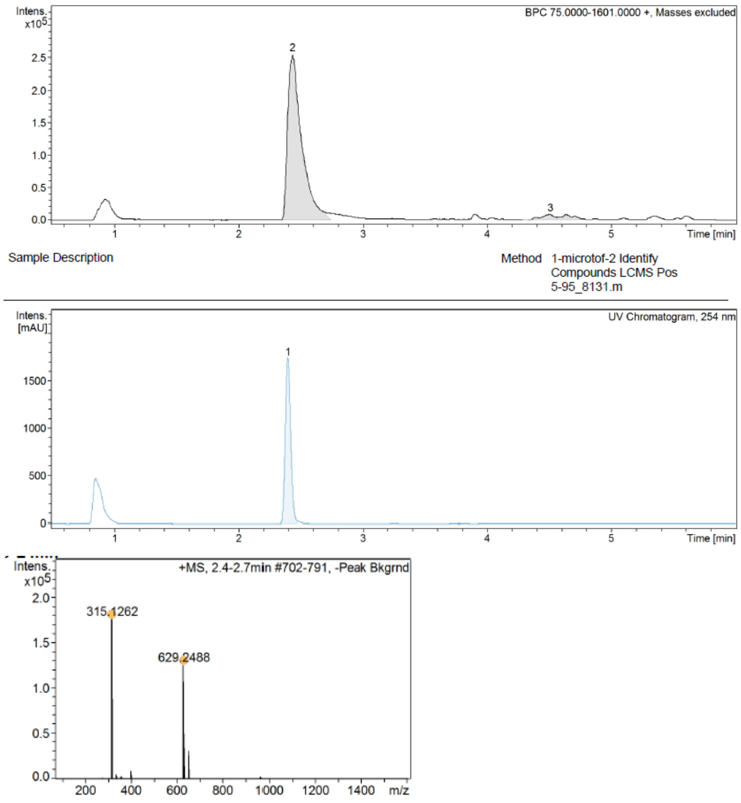
Intens. x10^5
BPC 75.0000-1601.0000 +, Masses excluded
Time [min]
Sample Description
Method 1-microtof-2 Identify Compounds LCMS Pos 5-95_8131.m
Intens. [mAU]
UV Chromatogram, 254 nm
Time [min]
Intens. x10^5
+MS, 2.4-2.7min #702-791, -Peak Bkgrnd
315.1262
629.2488
m/z

**48**

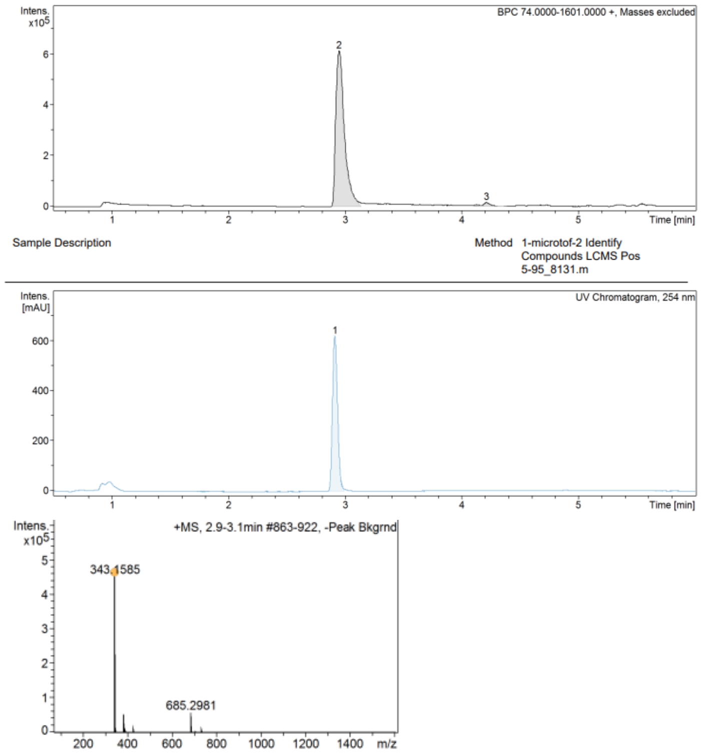
Intens. x10^5
BPC 74.0000-1601.0000 +, Masses excluded
Time [min]
Sample Description
Method 1-microtof-2 Identify Compounds LCMS Pos 5-95_8131.m
Intens. [mAU]
UV Chromatogram, 254 nm
Time [min]
Intens. x10^5
+MS, 2.9-3.1min #863-922, -Peak Bkgrnd
343.1585
685.2981
m/z

**49**

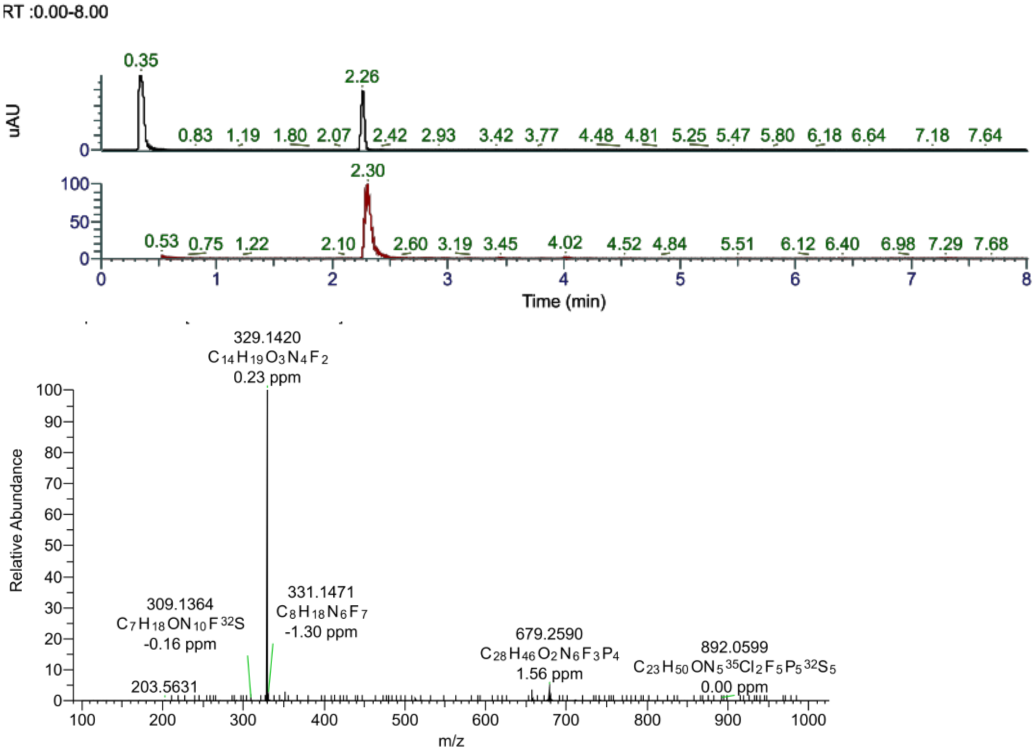


### Supplementary Methods

#### *In vivo* metabolite identification of 11.

**Sample Preparation**: Blood samples (30 µL) from the 10 mg/kg orally dosing PK study were vortexed with LCMS grade acetonitrile (90 µL) and centrifuged (13k rpm for 5 min) to remove any precipitated proteins. Supernatant (80 µL) from each sample was then added to 80 µL water (Milli-Q) and vortexed in preparation for LCMS analysis.

**Analysis**: Samples were analysed using a Waters Acquity UPLC with a diode array detector and Waters Xevo Q-TOF mass spectrometer. Reversed phase separation of the metabolites was accomplished using gradient elution method. The following elution profile was used: Eluent A, water plus 0.01% formic acid. Eluent B, acetonitrile plus 0.01% formic acid. Hold at 5% A for 0.5 min. Linearly increase to 40% B over 3.5 min. Then linearly increase to 95% B over another 2 min. One minute re-equilibration. Total run time was 7 min. Flow rate was 0.5 mL/min. Separation was achieved using a Waters BEH C18 column (50 x 2.1 mm, 1.7 µm particle size. 130 Å pore size, Cat. No. 186002350). The column temperature was 40°C. The UV response was monitored from 200-400 nm but not used in this study.

The mass spectrometer was operated using an electrospray (ESi) source in positive mode only. The source temperature was set to 120°C and the capillary voltage to 1.5 kV. The sample cone had an applied potential of 40V. For initial metabolite identification $MS^2$ (ms/ms) spectra were obtained using the $MS^e$ mode on the Waters

QToF spectrometer. Possible metabolites were then confirmed by re-injecting the test sample and collecting targeted $MS^2$ spectra, using the retention time and mass of the metabolite. A ramped collision energy (20, 30 and 40 V) was used for fragmentation of the metabolites by CID. These spectra were used for metabolite identification.
