## Supplementary material for "Design and development of lysyl tRNA synthetase inhibitors, for the treatment of tuberculosis": Data and PAINS summary

| Paper Number | LysRS $IC_{50}$ (μM) | KARS1 $IC_{50}$ (μM) | HepG2 (μM) | MIC (μM) OD | MIC (μM) Resazurin | Intra IC50 (μM) | Mics | Heps | SMILES | PAINS alert |
|---|---|---|---|---|---|---|---|---|---|---|
| **1** | 42 | >100 | >50 | 20 | 36 | 20 | <0.5 | ND | Nc1ncc2c(n1)CN(C1CCCC1)C2=O | No |
| **2** | 6.0 | 44 | >50 | ND | 4.9 | 2.5 | 0.7 | ND | Nc1ncc2c(n1)CN(C1CCCCC1)C2=O | No |
| **3** | 2.0 | 73 | >50 | ND | 2.7 | 0.40 | 0.8 | ND | Nc1ncc2c(n1)CN(C1CCCCCC1)C2=O | No |
| **4** | 21 | >100 | >50 | 7.9 | 20 | 13 | 0.7 | ND | Nc1ncc2c(n1)CN([C@H]1CCCC[C@H]1O)C2=O | No |
| **5** | >50 | ND | >50 | ND | ND | ND | ND | ND | Nc1ncc2c(n1)CN([C@H]1CCCC[C@@H]1O)C2=O | No |
| **6** | 0.3 | 1.8 | 1.4 | 0.16 | ND | ND | 1.2 | 5.3 | Nc1nc(c2c(n1)CN(C1CCCCC1)C2=O)Cl | No |
| **7** | 0.2 | 1.1 | 3.3 | 0.05 | ND | ND | 3.2 | 18 | Nc1nc(c2c(n1)CN(C1CCCCCC1)C2=O)Br | No |
| **8** | 0.3 | 60 | 46 | 0.40 | 0.3 | 0.16 | <0.5 | 0.9 | COc1nc(nc2c1C(=O)N(C2)C1CCCCC1)N | No |
| **9** | 0.2 | 56 | 26 | 0.13 | ND | 0.25 | 0.6 | 0.5 | COc1nc(nc2c1C(=O)N(C2)C1CCCCCC1)N | No |
| **10** | 0.4 | >100 | >100 | ND | 0.9 | 0.40 | 1.6 | 5.4 | CCOc1nc(nc2c1C(=O)N(C2)C1CCCCC1)N | No |
| **11** | 0.2 | >100 | >100 | 1.3 | 0.6 | 0.25 | 1.4 | 3.9 | CCOc1nc(nc2c1C(=O)N(C2)C1CCCCCC1)N | No |
| **12** | 0.3 | >100 | >50 | 1.3 | 1.3 | 0.40 | 4.4 | ND | CCCOc1nc(nc2c1C(=O)N(C2)C1CCCCCC1)N | No |
| **13** | 7.0 | >100 | >100 | ND | 60 | ND | <0.5 | ND | Nc1nc2c(c(n1)OCCO)C(=O)N(C2)C1CCCCC1 | No |
| **14** | 3.2 | >100 | >100 | ND | 30 | ND | 0.8 | ND | COCCOc1nc(nc2c1C(=O)N(C2)C1CCCCC1)N | No |
| **15** | 35 | >100 | >100 | ND | 80 | ND | 1.4 | 6 | CC(F)(F)COc1nc(nc2c1C(=O)N(C2)C1CCCCC1)N | No |
| **16** | 0.3 | 27 | 36 | ND | 0.6 | ND | 1.1 | ND | Nc1nc2c(c(n1)OC(F)F)C(=O)N(C2)C1CCCCC1 | No |
| **17** | 0.9 | 67 | >50 | 7.9 | 5 | 2.5 | 1.5 | 3.8 | Nc1nc2c(c(n1)OCC(F)F)C(=O)N(C2)C1CCCCC1 | No |
| **18** | 0.6 | 33 | 41 | 1.6 | 2.5 | 1.3 | 1.1 | 3.1 | CNc1nc(nc2c1C(=O)N(C2)C1CCCCC1)N | No |
| **19** | 1.1 | 66 | >100 | 6.3 | 5 | 2.5 | 6.4 | 16 | CCNc1nc(nc2c1C(=O)N(C2)C1CCCCC1)N | No |
| **20** | 50 | >100 | >100 | ND | ND | ND | 10 | 14 | CN(C)c1nc(nc2c1C(=O)N(C2)C1CCCCC1)N | No |
| **21** | 0.3 | 14 | 15 | 0.20 | <0.16 | 0.50 | 1.2 | 3.3 | Nc1nc2c(c(n1)C#N)C(=O)N(C2)C1CCCCC1 | No |
| **22** | 0.6 | >100 | >100 | 1.6 | ND | 2.5 | 0.5 | 3.2 | COCc1nc(nc2c1C(=O)N(C2)C1CCCCCC1)N | No |
| **23** | 3.2 | >100 | >100 | 50 | 40 | ND | 19 | 8.1 | CCOc1nc(nc2c1C(=O)N(C2)c1ccccc1)N | No |
| **24** | 0.2 | 32 | 47 | 0.20 | ND | ND | <0.5 | <0.5 | COc1nc(nc2c1C(=O)N(C2)[C@@H]1CCCC[C@@H]1O)N | No |
| **25** | 0.3 | >100 | >100 | 0.50 | 1.3 | 0.50 | <0.5 | 0.6 | CCOc1nc(nc2c1C(=O)N(C2)[C@@H]1CCCC[C@@H]1O)N | No |
| **26** | 1.3 | >100 | >100 | 1.6 | ND | 4.0 | <0.5 | <0.5 | COc1nc(nc2c1C(=O)N(C2)[C@@H]1CCC[C@H](O)C1)N | No |
| **27** | 0.4 | >100 | >100 | 1.0 | ND | 1.3 | <0.5 | <0.5 | COc1nc(nc2c1C(=O)N(C2)[C@H]1CC[C@@H](O)CC1)N | No |
| **28** | 0.9 | >100 | >100 | 1.3 | ND | ND | <0.5 | <0.5 | COc1nc(nc2c1C(=O)N(C2)[C@H]1CC[C@@H](CO)CC1)N | No |
| **29** | 0.4 | >100 | >100 | 2.0 | 2.5 | 1.0 | <0.5 | 1.2 | CCOc1nc(nc2c1C(=O)N(C2)C1CCCCC1(F)F)N | No |
| **30** | 0.5 | >100 | >100 | 5.0 | 1.9 | 0.79 | <0.5 | 1 | CCOc1nc(nc2c1C(=O)N(C2)C1CCCC(F)(F)C1)N | No |
| **31** | 1.5 | >100 | >100 | 13 | 15 | ND | <0.5 | 1.5 | CCOc1nc(nc2c1C(=O)N(C2)C1CCC(F)(F)CC1)N | No |
| **32** | 0.2 | >100 | 93 | 0.32 | ND | 0.25 | <0.5 | <0.5 | COc1nc(nc2c1C(=O)N(C2)[C@H]1CCCCC1(F)F)N | No |
| **33** | 0.9 | 64 | 32 | 3.2 | ND | 1.6 | <0.5 | <0.5 | COc1nc(nc2c1C(=O)N(C2)[C@@H]1CCCCC1(F)F)N | No |
| **34** | 2.2 | >100 | >100 | >100 | >80 | ND | <0.5 | 0.9 | CCOc1nc(nc2c1C(=O)N(C2)[C@@H]1CCCC[C@@H]1N)N | No |
| **35** | 0.3 | 27 | 15 | 2.5 | ND | ND | 3.4 | 6.2 | COc1nc(nc2c1C(=O)N(C2)C1CCCCC1(C)C)N | No |
| **36** | 0.2 | >100 | >100 | 0.63 | ND | 0.40 | <0.5 | 0.6 | CCOc1nc(nc2c1C(=O)N(C2)C1CCCCCC1O)N | No |
| **37** | 0.1 | 60 | 60 | 0.25 | ND | 0.50 | <0.5 | <0.5 | COc1nc(nc2c1C(=O)N(C2)C1CCCCCC1O)N | No |

| | | | | | | | | | | |
|---|---|---|---|---|---|---|---|---|---|---|
| **38** | 0.1 | >100 | >100 | 0.16 | ND | 0.16 | <0.5 | 0.8 | COc1nc(nc2c1C(=O)N(C2)C1CCCCCC1(F)F)N | No |
| **39** | 21 | >100 | >100 | >100 | ND | ND | <0.5 | ND | COc1nc(nc2c1C(=O)N([C@H]1CC[C@@H](O)CC1)C2(C)C)N | No |
| **40** | 12 | >100 | >100 | ND | ND | ND | <0.5 | ND | CCOc1nc(nc2c1C(=O)N(N2)C1CCCCCC1)N | No |
| **41** | 8.0 | >100 | >100 | | ND | ND | 39 | ND | CCOc1nc(nc2c1C(=O)N(C1CCCCCC1)N2CC1CC1)N | No |
| **42** | 1.5 | 56 | >100 | 50 | ND | ND | <0.5 | <0.5 | CCOc1nc(nc2c1C(=O)N(C1CCCCCC1)N2CCC1CNC1)N | No |
| **43** | 1.3 | >100 | >100 | 7.9 | ND | 5.0 | 12 | ND | CCOc1nc(nc2c1C(=O)N(C2)C1CCCCC1)NC | No |
| **44** | 0.6 | >100 | >100 | 0.40 | ND | ND | 9.3 | ND | COc1nc(nc2c1C(=O)N(C2)C1CCCCC1)NC1CC1 | No |
| **45** | 0.4 | >100 | >100 | 1.6 | ND | 2.5 | 1.1 | 3.6 | COc1nc(nc2c1C(=O)N(C2)[C@@H]1CCCC[C@@H]1O)NC1CC1 | No |
| **46** | 0.18/0.62* | >100 | >100 | 0.40 | ND | 0.32 | <0.5 | <0.5 | CCOc1nc(nc2c1C(=O)N(C2)[C@H]1[C@@H](O)CCCC1(F)F)N | No |
| **47** | 0.10/0.28* | >100 | 68 | 0.13 | ND | 0.13 | <0.5 | <0.5 | COc1nc(nc2c1C(=O)N(C2)[C@H]1[C@@H](O)CCCC1(F)F)N | No |
| **48** | 0.11/0.08* | >100 | 54 | 0.13 | ND | 0.10 | <0.5 | 2.6 | CCOc1nc(nc2c1C(=O)N(C2)[C@H]1[C@@H](O)CCCCC1(F)F)N | No |
| **49** | 0.14/0.05* | 1219 | >100 | 0.04 | ND | 0.03 | <0.5 | 1.2 | COc1nc(nc2c1C(=O)N(C2)[C@H]1[C@@H](O)CCCCC1(F)F)N | No |
